# Predicting chemotherapy-induced thrombotoxicity by NARX neural networks and transfer learning

**DOI:** 10.1101/2024.08.06.606816

**Authors:** Marie Steinacker, Yuri Kheifetz, Markus Scholz

## Abstract

**Background:** Thrombocytopenia is a common side effect of cytotoxic chemotherapies, which is often dose-limiting. Predicting an individual’s risk is of high clinical importance, as otherwise, a small subgroup of patients limits dosages for the overall population for safety reasons.

**Methods:** We aim to predict individual platelet dynamics using non-linear auto-regressive networks with exogenous inputs (NARX). We consider different architectures of the NARX networks, namely feed-forward networks (FNN) and gated recurrent units (GRU). To cope with the relative sparsity of individual patient data, we employ transfer learning (TL) approaches based on a semi-mechanistic model of hematotoxicity. We use a large data set of patients with high-grade non-Hodgkin’s lymphoma to learn the respective models on an individual scale and to compare prediction performances with that of the semi-mechanistic model.

**Results:** Of the examined network models, the NARX with GRU architecture performs best. In comparison to the semi-mechanistic model, the network model can result in a substantial improvement of prediction accuracy for patients with irregular dynamics, given well-spaced measurements. TL improves individual prediction performances.

**Conclusion:** NARX networks can be utilized to predict an individual’s thrombotoxic response to cytotoxic chemotherapy treatment. For reasonable model learning, we recommend at least three well-spaced measurements per cycle: at baseline, during the nadir phase and during the recovery phase. We aim at generalizing our approach to other treatment scenarios and blood lineages in the future.

## 1 Introduction

Cytotoxic cancer therapies are often accompanied by major side-effects, such as thrombocytopenia, which are dose-limiting (Pfreundschuh, 2004a,b). Thrombocytopenia is associated with a higher risk for uncontrolled bleeding and shorter patient survival (Elting et al, 2001), but the extent of this side effect is highly heterogeneous among patients. Thus, for safety reasons, a possibly small subset of patients impose treatment limitations for the general patient population. Predicting an individual’s risk is therefore of high clinical importance.

To achieve this goal, a number of statistical (Wunderlich et al, 2003), semimechanistic (Friberg et al, 2002; Friberg and Karlsson, 2003; Mangas-Sanjuan et al, 2015; Henrich et al, 2017) and comprehensive mechanistic models (Scholz et al, 2010; Kheifetz and Scholz, 2019, 2021) were proposed but with limited success. Existing statistical models typically use only cross-sectional data of a patient and do not consider actually given therapies and resulting individual dynamics. This results in high residual patient heterogeneity, hampering proper risk stratification and individual treatment decisions. Semi-mechanistic and mechanistic models allow, in principle, individual predictions. But, this approach requires estimation of model parameters based on individual data under therapy which are typically sparse. (Kheifetz and Scholz, 2019, 2021).

Neural networks are a data-driven modeling approaches, which in theory can represent arbitrary dynamical systems (Hornik et al, 1989). They are widely applied to prediction tasks also in clinical settings, e.g. Choi et al (2016); Esteban et al (2016); Li et al (2020). To cope with the sparse data problem of learning individual time series, we proposed a transfer learning approach exploiting (semi-) mechanistic models of haematotoxicity (Steinacker et al, 2023). Recently, this approach has been examined in a wider range of biological modeling problems (Kleissl et al, 2023; Zabbarov et al, 2024).

To further optimize this approach, we present here a comparative analysis of different neural network architectures and learning methods, including a data augmentation approach based on a frequently used semi-mechanistic model of Friberg et al. (Friberg et al, 2002). For learning the models and comparing it’s prediction performances, we consider real patient data from a study of high-grade non-Hodgkin’s lymphoma patients (Pfreundschuh, 2004a,b).

## 2 Methods

To predict individual patient time courses we employ and compare different types of feed-forward or recurrent neural networks (RNNs). Architectures are optimized to cope with the relative sparsity of clinical data. As an alternative, we consider a simple semi-mechanistic model having a limited number of parameters which can be estimated based on individual patient data. We also employ and compare different learning methods for the neural network models including transfer learning from the semi-mechanistic model to allow training of individual networks. We explain these approaches in the following. A more comprehensive presentation is provided in the supplement, section 1.1 (Online Resource 1). To compare the approaches, we consider real patient data as explained in the following section.

### 2.1 Patient data

We consider patient data from the NHL-B study comparing CHOP-like therapies of high-grade non-Hodgkin’s lymphoma in young and elderly patients (Wunderlich et al, 2003; Pfreundschuh, 2004a,b). Patients received either 6xCHOP-14/21 or 6xCHOEP14/21. Relatively dense data of blood counts are available for most of the patients. Since we aim at predicting individual next cycle platelet toxicity, we here consider patients with platelet data available for at least four therapy cycles and more than one blood count in the first cycle. We exclude patients who received platelet transfusions. This results in a total of 360 patients eligible for analysis.

To assess the impact of data sparsity on prediction performance, patients are stratified into two groups. The first group comprises 135 patients for which at least five platelet counts are available per cycle. This group represents patients whose individual platelet dynamics is well-covered by the available data. All other selected patients are summarized in the second group, representing reduced coverage of the platelet dynamics. Due to stronger dynamics, we expect less favorable prediction performances for the 14 day CHO(E)P therapies (d14) compared to the 21 day (d21) schedules. Thus, we consider four groups in the following analyses, denoted by De14/De21 for patients with dense measurements and Sp14/Sp21 for patients with sparse coverage of their individual platelet dynamics.

### 2.2 Semi-mechanistic model of platelet dynamics

As a reference to our NARX modeling, we consider a semi-mechanistic model of platelet dynamics under chemotherapy proposed by Friberg et al (2002). This is a simple compartment model describing proliferating stem cells maturing into circulating blood cells. Maturation is modeled by three sub-compartments with first order transitions with transit time MTT. The feedback strength of the circulating compartment towards the proliferating compartment is represented by parameter *γ*. Individual steady-states can be defined by parameter *C*_0_.

The treatment effect of chemotherapy is modeled by a step-wise toxicity function acting on the proliferation compartment. We assume a simple step-function returning to zero after one day. The step height represents the dose of the chemotherapy normalized to the body surface area. The individual toxic effect is represented by parameter *E_eff_* multiplied to this dose level. Model equations and parameter ranges are presented in the supplement, section 1.2 (Online Resource 1). Individual parameters can be derived based on likelihood optimization of the agreement of model and data penalizing strong deviations from the population average. Details are explained in the supplement, section 1.2 (Online Resource 1).

### 2.3 NARX modeling

Nonlinear auto-regressive models with exogenous inputs (NARX) are a class of models suitable to describe dynamics of discrete time series data affected by external factors. Dynamics are captured by learning a functional relationship between observations or predictions at previous time steps, an externally imposed input function, which in our situation is the applied therapy, and the current state. In our situation, this function is learned via neural networks, further called NARX neural networks. All NARX neural networks are recurrent neural networks (RNNs), and the recurrent connection between the output of the network and the input is formed by a so-called tapped delay line (Siegelmann et al, 1997). In this work, we compare different network architectures regarding their learning abilities and prediction performances.

To assess and improve prediction performances, we consider different architectures of the NARX modeling framework. First, we consider a simple feed-forward network for coupling as proposed by Steinacker et al (2023). We call this architecture ARX-FNN in the following. In such a network, information only flows in one direction through the network, from the input nodes through potential so-called hidden layers of nodes to the output nodes of the network. We compare this simple framework with more complex alternatives. Specifically, we consider a so called gated-recurrent unit (GRU) network (Cho et al, 2014), which replaces the FNN by an RNN (ARX-GRU). In addition to the FNN weights, RNNs have hidden states which store the past contextual information of an input sequence. This architecture allows the network to memorize relevant information for a longer period of time or neglect such information if not required to improve prediction. The choice of the specific GRU architecture is justified in the supplement, section 1.1 (Online Resource 1).

### 2.4 Prediction approach using NARX models

We aim at predicting individual time courses of platelet dynamics. The exogenous input represents applications of cytotoxic drugs. Essentially, the network predicts daily platelet counts based on measured or predicted values at previous days considering the therapy input. Since patients are untreated at study inclusion, we assume that platelet histories of patients are in steady-states equal to the respective baseline measurements of patients. Figure 1, illustrates our modeling concept.

**Fig. 1.**
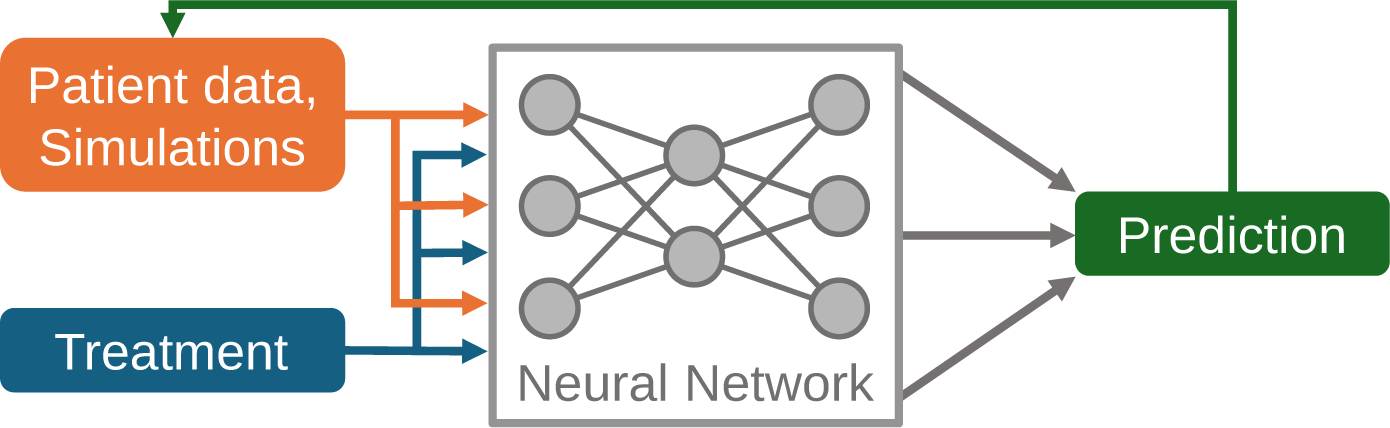
NARX neural network models for the purpose of predicting individual platelet dynamics under chemotherapy. The neural network is used to predict future platelet counts of a patient based on previous measured or predicted counts and the provided therapy. The architecture of the neural network can be modified and learned based on individual time series data of patients under treatment.

For predicting individual patient dynamics with our NARX frameworks, it is necessary to scale and transform the patient data. First, we log-transform platelet counts due to their log-normal shape of distribution. We further scale the data such that the log-transformed range between degree four thrombocytopenia, 25 *×* 10^9^cells*/*l and the average initial state in the Sp group, 300 *×* 10^9^cells*/*l (see below), corresponds to the range [*−*0.5, 0.5] for ARX-GRU models, and [*−*1, 1.5] for ARX-FNN models. These ranges are the result of a hyperparameter optimization step. Of note, this scaling does not affect the comparability of objective functions between models.

### 2.5 Transfer Learning approach for neural network models

Transfer learning (TL) techniques are developed to cope with sparse data problems in machine-learning (Pan and Yang, 2010). We here adopt a transfer from (semi-)mechanistic models as proposed recently by us and others, e.g. Steinacker et al (2023); Kleissl et al (2023); Zabbarov et al (2024). In more detail, we parameterize the semi-mechanistic model for given individual patient data and simulate dynamics for different therapy scenarios including the actually given therapy schedule (Steinacker et al, 2023). These simulated dynamics were used to pre-train our neural network for the respective individual patient. After pre-training, the weights of model inputs and hidden layers remained unchanged. Only weights of the linear output function are recalibrated based on patient’s real data. This step ensures that learned dynamics from the semi-mechanistic model are preserved while transferring the model to the real patient dynamics. Finally, a fine-tuning step is employed, re-training the whole model with the patient’s real data with a low learning rate. Details on regularization during the transfer learning procedure and the fine-tuning can be found in the supplement, section 1.3 (Online Resource 1). To assess the impact of pre-training on prediction performances, we compared this approach with a standard approach without pre-training. Scenarios are denoted “with TL” respectively “without TL” in the following.

### 2.6 Methods to reduce overfitting

Over-parametrization is a well-known problem of neural network models. We employ multiple methods to reduce overfitting during training of the individual networks. Depending on network architecture and training strategy, we combine different standard methods as explained in the following. Details of the specific methods are given in the supplement, section 1.3 (Online Resource 1).

Methods acting on the network structure comprised pruning and weight regularization. *Pruning* describes the removal of nodes or weights from the network to reduce the number of free parameters and to improve generalization performance (LeCun et al, 1990). We employ pruning for all training processes considered. The *dropout method* is a form of non-permanent pruning, where weights are randomly removed from a network with a specified frequency (Srivastava et al, 2014). We apply this method only for training modes without TL. *Weight regularization* adds a penalty to the objective function of the model, which consists of the squared sum of the network weights. It is also known as weight decay (Krogh and Hertz, 1991). Again, this technique is applied for all training process.

Still, neural networks suffer from over-parametrization as they tend to assume noisy dynamics in case of larger time distances between measurements. To avoid this behavior, it is necessary to regularize such noisy dynamics. We adopt here *exponential smoothing* (Brown, 1956; Brown and Meyer, 1961; Holt, 2004), which acts as a low-pass noise filter. We apply this method to the network output in the following way:

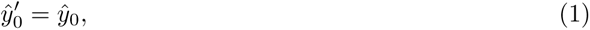

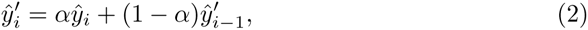

with smoothing factor *α ∈* [0, 1], implemented as a trainable hyperparameter of the network model. Here, the model prediction corresponds to *ŷ^′^* while the regressed variable *ŷ* is used in the auto-regressive feedback loop of the network. To implement exponential smoothing, we include a respective penalty term in the objective function.

Another established method to avoid overfitting due to small data-sets is the addition of a noise term to the training data, which can be seen as a form of data augmentation (Goodfellow et al, 2016). It encourages the neural network output to be a smooth function of its input and weights, respectively (An, 1996). We add here a Gaussian noise to the scaled network inputs. This augmentation is only active in the training phase, where dynamics of real patient data are learned.

### 2.7 Comparison of prediction performances

We use two fitness measures to assess and compare prediction performances between our learning frameworks. First, we consider the modified mean-squared error (*SMSE*), which is also employed during network training on real patient data. Since time points of measurements are not exactly known, we use a weighted average of the mean squared error over three days (weight 0.3 for the previous or next day, weight 1.0 for the actual day). This performance measure is useful for training individual models and comparing them in terms of accuracy of platelet predictions.

For a clinically more useful and interpretable measure of prediction performance, we determine the difference of predicted and observed thrombocytopenia degrees (*DD_c,t_*) at the nadir phase of each therapy cycle. To determine the thrombocytopenia degree, we adhere to the criteria of the National Cancer Institute (National Cancer Institute (NCI), 2009). The degree is classified as follows for cell count *C*:

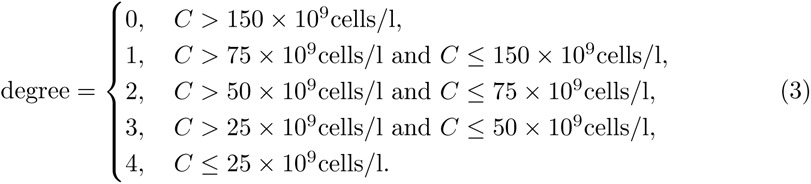

The *DD_c,t_* is calculated as follows,

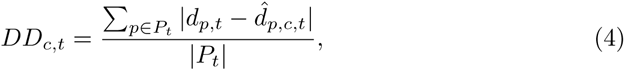

where *c* denotes the number of calibration cycles utilized, *t* denotes the cycle for which the prediction is made, *P_t_* is the set of patients with measurements for that cycle, *d* denotes the observed thrombocytopenia degree in that cycle for patient *p*, and *d̂* is the respective prediction at this time point. This fitness measure was proposed by Kheifetz and Scholz (2021) to evaluate and compare predictive performances between models. The advantage of this fitness measure is that it determines clinically relevant differences between predicted and observed platelet counts in terms of toxicity grades.

## 3 Results

First, we analyze our patient population regarding thrombotoxicity and compare the sets of patients with dense vs. sparse time series data. Next, we compare our learning approaches with respect to their prediction performances for the different patient groups. For this purpose, we learn models of single patients using different numbers of calibration cycles.

### 3.1 Distribution of thrombotoxicity across analyzed patients

To test the dependence of model performances on data sparsity, we divide the selected patients into two groups of dense (De) vs. sparse (Sp) spacing of available measurements. Respective patient characteristics are displayed in table 1.

**Table 1.** Comparison of patient groups of different data sparsity. We present distributions of risk factors and therapies for all patients and the subgroups of patients with dense (De) and sparse (Sp) time series data. We compare the latter by *χ*^2^-tests for categorical data and *t*-Test for continuous data to assess structural differences between these groups.

| Risk factor |  | All | De | Sp | p-value |
| --- | --- | --- | --- | --- | --- |
| mean initial platelet count $\times 10^9 \text{ L}^{-1}$ | | 291 | 276 | 300 | 0.096 |
| age | young ($\leq 60$ years) | 207 | 59 | 148 | $6.6 \times 10^{-5}$ |
| | old ($> 60$ years) | 153 | 76 | 77 | |
| sex | male | 202 | 65 | 137 | 0.025 |
|  | female | 158 | 70 | 88 |  |
| treatment | CHOP-14 | 71 | 25 | 46 | 0.76 |
|  | CHOEP-14 | 87 | 28 | 50 |  |
|  | CHOP-21 | 94 | 37 | 66 |  |
|  | CHOEP-21 | 108 | 45 | 63 |  |
| **total** |  | **360** | **135** | **225** |  |

It reveals that the subgroup of patients with denser measurements is older (*p* = 6.6 *×* 10*^−^*^5^) and contains more females (p=0.025) compared to the subgroup of patients with sparser measurements, which reflects the more intense monitoring of high risk patients. In contrast, distributions of treatments (p=0.76) and initial platelet counts are comparable (p=0.096) between groups.

Next, we analyze the degree of thrombotoxicity between these groups and between d21 vs. d14 schedules. Results are shown in figure 2. As expected, toxicity is higher at later therapy cycles and higher for d14 schedules compared to d21 schedules. Moreover, the toxicity is also higher in the De vs. the Sp group. The latter is expected, on one hand, due to the shift in risk factors we observe between these groups. On the other hand, higher toxicities are expected also *per se* in the De group since the more intense monitoring increases the chance of capturing the nadir phase of patients.

**Fig. 2.**
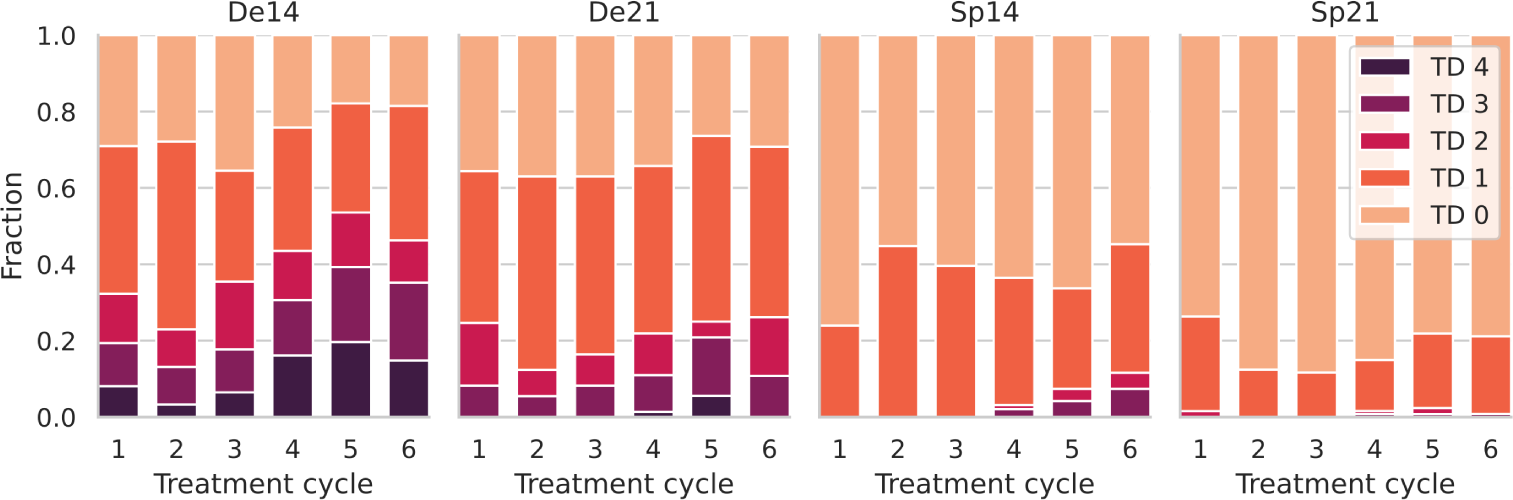
Distribution of observed thrombocytopenia degrees (TD) per treatment cycle, separated for patients with dense vs. sparse measurements and 14 vs. 21 days cycle duration. The fraction of patients with a higher thrombocytopenia degree increases with treatment cycles for all groups. As expected, higher toxicities are observed in the 14d vs. the 21d schedules. We also observe higher toxicity in the De compared to the Sp group.

### 3.2 Comparison of model performances

We apply five learning frameworks to these data, namely ARX-GRU vs. ARX-FNN with and without TL and the semi-mechanistic Friberg model as control. To learn the models, we provide data of one to five therapy cycles per patient and predict the subsequent cycles. Comparisons of learning frameworks are based on their residual sum of squares (*SMSE*) of predicted values and the deviation of observed and predicted degrees of thrombocytopenia (*DD_c,t_*). By this approach, we aim at answering the following questions: (1) Which NARX architecture performs better? (2) Is transfer learning advantageous for the NARX frameworks? (3) Do the NARX frameworks result in better prediction performances compared to the Friberg model? (4) How does data sparsity and cycle duration influence these results?

#### Evaluation of SMSE

Performances with respect to *SMSE* are shown in figure 3. As expected, prediction performance generally increases if more training cycles are provided. Regarding NARX architecture, it revealed that GRU performs generally better then FNN. Moreover, TL shows an advantage in most situations. GRU with TL performed almost always better then without TL. Comparing the ARX-GRU with TL with the Friberg model, there is a clear difference in performance depending on the patient group. For the De14 group, the ARX-GRU with TL has a better prediction performance. For Groups De21 and Sp21, the ARX-GRU with TL performs better then Friberg’s model if three respectively two or more cycles are provided to learn the model. However for group Sp14, Friberg’s model slightly outperforms the ARX-GRU with TL. Prediction performances of the d21 treatments are generally better than those of the d14 treatments. Surprisingly, prediction performances of the Sp groups are better than that of the De groups. However, this fact could be explained by the better coverage of the true dynamics within the De groups and the different risk profiles of the groups.

**Fig. 3.**
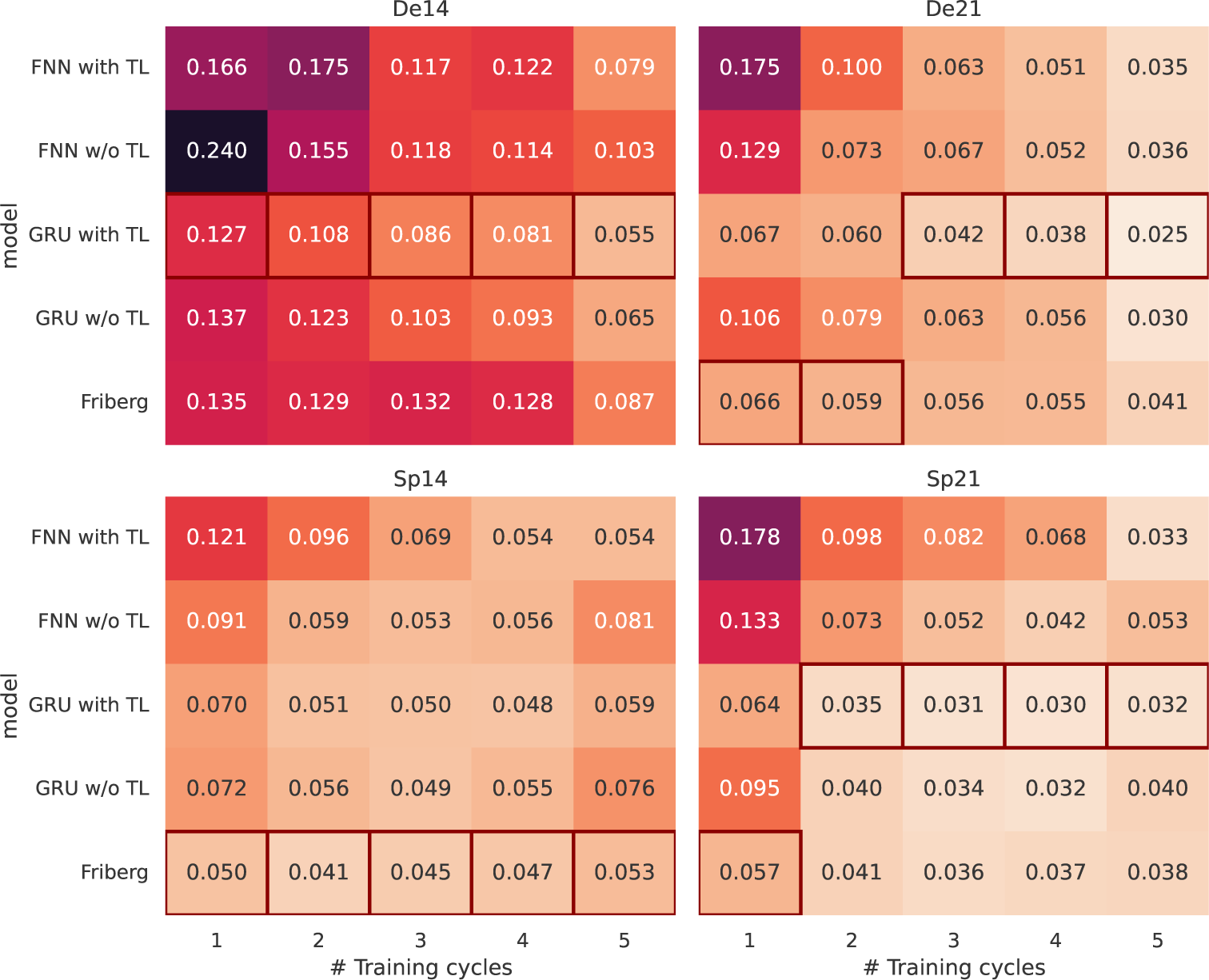
Prediction performance measured as *SMSE* for the different learning scenarios. We present the average *SMSE* for the different groups of patients (panels), number of treatment cycles used to train the model (columns) and learning approaches (rows). The *SMSE* is determined by data of subsequent cycles not used for model training. The *SMSE* scale is highlighted by coloring, with the lighter shade corresponding to lower *SMSE*, i.e. better prediction performance. Per column and panel, the scenario of the lowest *SMSE* is marked by a red rectangle.

#### Evaluation based on difference of thrombocytopenia degrees

Next, we compare model performances for the *DD_c,t_* metric representing a clinically more relevant evaluation of differing toxicity degrees. Here, we focus on the ARX-GRU with TL framework which outperformed the other NARX frameworks and compare this approach with the results of Friberg’s model. Results are shown in figure 4. Again, performance of both models tend to be better if more training cycles are provided. However, this trend is more pronounced for the ARX-GRU framework.

**Fig. 4.**
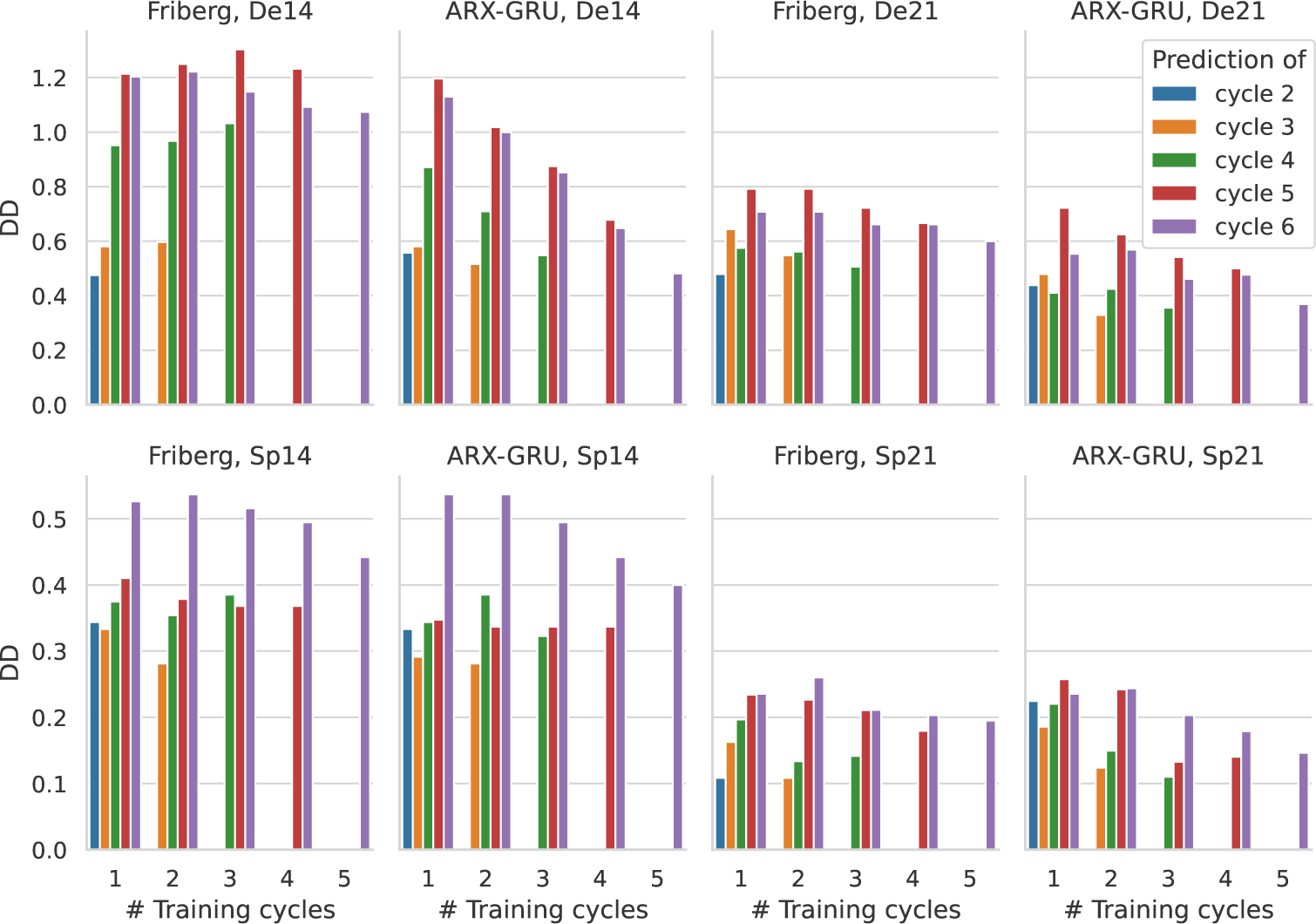
Comparison of *DD_c,t_* between ARX-GRU and Friberg model. We compare the ARX-GRU model with TL with Friberg’s model regarding *DD_c,t_* for the different patient groups. Due to generally increased toxicity, scales of y-axis differ between De (upper row) and Sp groups (lower row). Number of cycles used for model training are provided at the x-axes. Colors correspond to cycles for which *DD_c,t_* could be determined.

As expected, prediction performance decreases for later cycles. In almost all scenarios, the ARX-GRU performs better than the Friberg model. Only in cases of only one training cycle, both models perform comparable. Prediction performances differ between patient groups. *DD_c,t_* for the De group is generally higher then for the Sp group which is expected due to the better coverage of the nadir phase and the increased risk profile of the De group (see Section 3.1). *DD_c,t_* is also higher for the d14 treatments compared to the d21 treatments because the d14 treatments represent more intense therapy schedules resulting in deeper nadir phases.

### 3.3 Results of individual parameter fitting

To illustrate the learning process of the ARX-GRU with TL vs. Friberg’s model, we present in Fig. 5 a few examples featuring certain aspects of the different learning behaviors. Results of all patients are provided as supplement material (Online Resource 2).

**Fig. 5.**
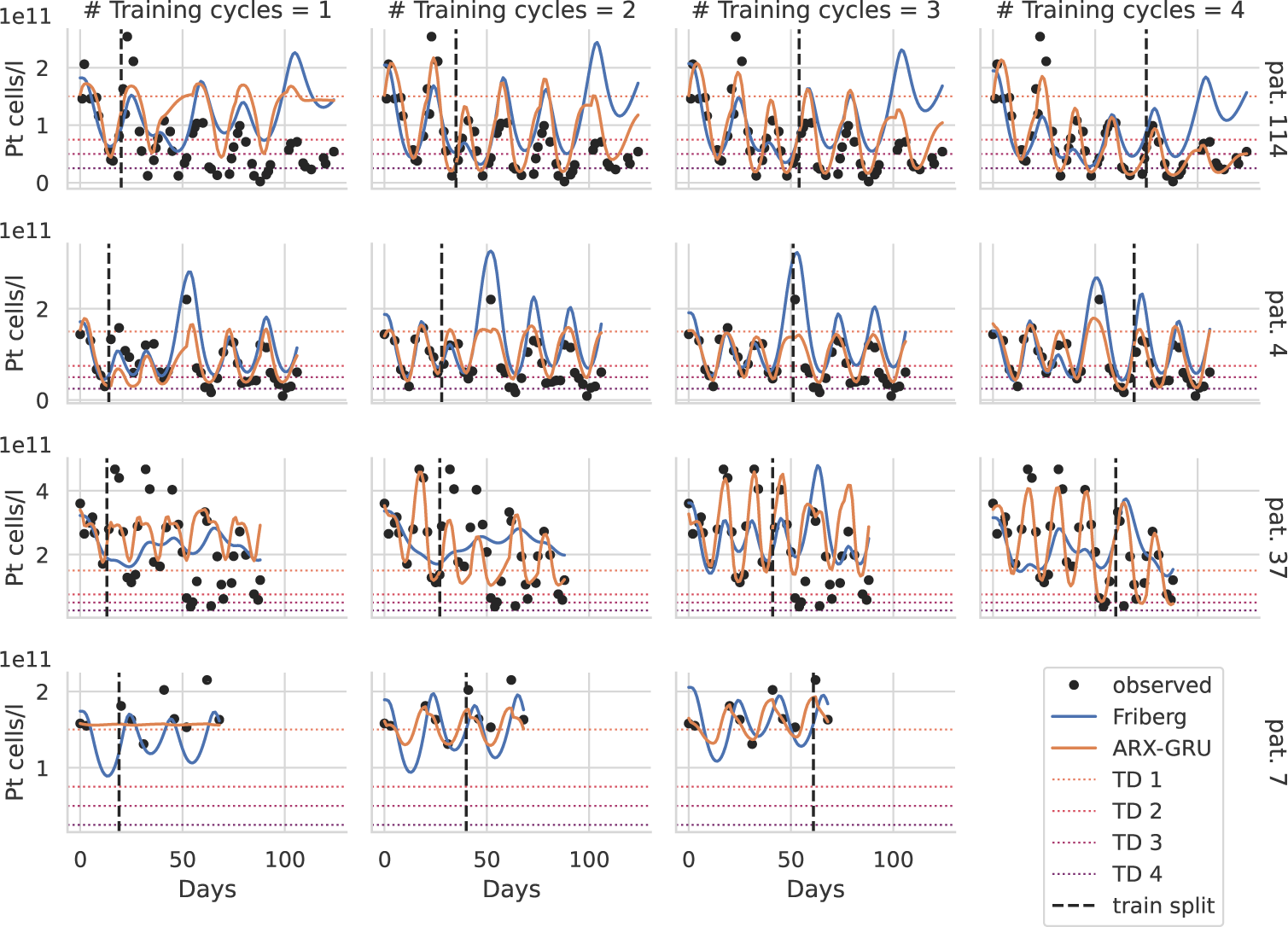
Learning dynamics of four selected patients. For four patients (4, 7, 37, 114), we present predictions of the ARX-GRU with TL (orange) and the Friberg model (blue) based on one to four training cycles (columns). The dashed lines separate the training and prediction periods. Data points are shown in black. Different scales were used for each patient (rows). For better interpretation, thrombocytopenia degrees are presented as dotted lines. While patients 4, 37 and 114 are from group De14, patient 7 belongs to Sp21.

Example patient #114 was treated with CHOEP-14 and showed thrombocytopenia degree three in the first cycle, and degree four in cycles two and later. Providing only one cycle of data resulted in inferior prediction performance for both learning frameworks. However, providing data of the first two cycles resulted in good predictions of the ARX-GRU with TL even for late cycles, which further improves if more cycles are provided for training. In contrast, the Friberg model did not improve if more cycles are provided. We conclude that in cases of strong and cumulative toxicity, the ARX-GRU outperforms the Friberg model but at least two cycles of data are required to learn these dynamics.

Patient #4 was also treated with CHOEP-14 and showed thrombocytopenia degree three in the first treatment cycle. Treatment was delayed at the fourth treatment cycle, resulting in a platelet overshoot. While the ARX-GRU performs better in predicting future nadirs, it fails to predict this overshoot, which is captured by the Friberg model. This highlights some limitations of the NARX modeling framework. Due to strong regularization constraints, the ARX-GRU under-predicts spikes favoring the prediction of nadirs, which is assumed to be more clinically relevant.

Patient #37 also receiving CHOEP-14 treatment shows an unusual pattern of toxicity starting with mild degrees of zero or one up to cycle three but strong toxicities of degree three at later cycles. Both modeling frameworks fail to accurately predict this behavior even if more than two cycles for training are provided. ARX-GRU predictions become satisfactory after four training cycles where this pattern becomes apparent while prediction performance of the Friberg model remains dismal. We resume that unusual individual patterns or changes in individual patterns can not be learned based on early cycle dynamics.

All three described patients belong to the De group. As an example from the Sp group, we show patient #7 who received CHOP-21. Measured data are very sparse and do not capture the nadir phase of the patient during the first cycle since only two measurements at the start of the cycle were available. In such cases, the ARX-GRU model initially predicts an approximately constant dynamic due to lack of data and strong regularization penalties, while the prediction of the Friberg model represents average semi-mechanistic population dynamics which always assume a nadir phase. After providing two or more cycles, the ARX-GRU improves significantly showing better agreement with the nadir phases than the Friberg model. We conclude, that at least three well-spaced measurements per cycle covering the decline, the nadir and the recovery are required for good learning performance of the ARX-GRU.

## 4 Discussion

Here, we propose a universal framework to predict the dynamical course of platelets under cytotoxic chemotherapy applications based on learning individual network models. We identified the GRU-based NARX network architecture as a promising candidate. To cope with the problem of sparse individual patient data, we applied a transfer learning approach using a semi-mechanistic model of human hematopoiesis. We applied our approach to data of patients with high-grade non-Hodgkin’s lymphoma treated with CHOP-like chemotherapies.

A comparison of FNN and GRU architectures of our NARX approach revealed that the ARX-GRU performed better. Although both approaches allow in principle to model long-term feedbacks which are characteristic for hematopoiesis recovery, this system memory is handled more efficiently in the GRU setting, allowing to keep essential memory information rather than fixed memories (Cho et al, 2014). However, other architectures such as neural ODEs (Chen et al, 2018) and physics-informed neural networks (Raissi et al, 2019) could also be of interest and should be investigated in the future.

Due to the strong between-patient heterogeneity of platelet dynamics, we aimed at learning individual models per patients. This approach also allows to consider individual therapy adaptations such as individual dosing schemes and therapy delays. On the other hand, this implies that often only a sparse data base is available to learn the model. Thus, we needed to adopt several techniques of building lean network architectures and to prevent overfitting of our model. We employed multiple rigorous regularization and reduction techniques to achieve this goal. The chosen techniques and their parameters were the result of hyperparameter optimization on a test set population of patients.

Importantly, we aimed at improving the learning process by introducing a transfer learning approach based on virtual therapy scenarios pre-simulated with the help of a semi-mechanistic model of hematotoxicity. It turned out that this approach largely improves the learning of the ARX-GRU architecture. We used the very simple but frequently applied Friberg model for that purpose. However, other alternatives (e.g. Mangas-Sanjuan et al (2015); Henrich et al (2017)) or even more complex mechanistic models such as Scholz et al (2010); Kheifetz and Scholz (2019, 2021) developed for predicting individual time series could also be considered in the future.

Prediction performance of our modeling approach depends on the structure of the provided data. First, we observed a better prediction performance for sparsely compared to densely measured time series. This seems to be contra-intuitive but could be explained by structural differences between respective patient populations. It turned out that the more densely measures time series correspond to patients of higher risk. Another issue is that denser time series improve the chance to cover the nadir phase, i.e. to measure stronger toxicities. This apparent bias between our defined De and Sp groups makes conclusions regarding the impact of measurement density difficult. Based on single learning dynamics, we conclude that having well-spaced measurements covering decline, nadir and recovery of the patient is more important than having many non-optimally distributed measurements. Second, d21 schedules resulted in better prediction performances than d14 schedules. Again, this can be explained by stronger toxicities observed for the latter accompanied by stronger, more irregular dynamics which are more difficult to predict. This is consistent with the work of Kheifetz and Scholz (2019, 2021).

Finally, we compared the prediction performance of our ARX-GRU with TL with that of the Friberg model itself. It turned out that the ARX-GRU framework performed better in the vast majority of scenarios. Strongest improvements were observed for the De14 group, where dynamics are not captured by the Friberg model but could be learned well by the ARX-GRU due to a sufficient data base. For a few individual learning scenarios with a very unfavorable spacing of measurements not covering the real patient dynamics, the ARX-GRU tend to learn a constant dynamics due to the imposed strong regularization measures required to develop a lean network architecture. Here, the Friberg model has an advantage due to the intrinsic dynamics assumed by the model. It is conceivable that this disadvantage of the ARX-GRU could be rectified by improved transfer learning methods.

With respect to clinical utility of our approach, we conclude the following. The ARX-GRU framework with transfer learning is in principle suitable to predict individual platelet dynamics with sufficient accuracy such as an average deviation of observed and predicted toxicity grades of less than 0.5. After providing at least two cycles of data, most of the individual dynamics are learned sufficiently well allowing precise predictions of toxicity grades even at later therapy cycles.

Our study has several limitations. We considered selected but frequently used network architectures of our NARX framework. It is conceivable, that refined architectures could further improve prediction performances of our model. The very simple Friberg model was used for transfer learning, because it does not make strong physiological assumptions, but other semi-mechanistic or mechanistic models could be worthwhile to investigate for transfer learning as well. We modeled time series of platelet counts under chemotherapy as an example. Modeling other hematopoietic lineages by the same framework is conceivable after some modifications. For example, growth factor applications could be considered as additional external factors imposed on our NARX network. This would allow, for example, modeling of white blood cell dynamics, which we plan to address in the near future. Finally, our analysis was based on data of a single study of therapies for high-grade non-Hodgkin’s disease. Spacing of measurements were not prescribed per protocol but was done at the discretion of the treating physicians introducing bias with respect to coverage of individual dynamics and prediction performances. We aim at testing our approach for other data sets including other diseases in the future.

## Supporting information

Online Resource 1

Online Resource 2

## Declarations

### Funding

The authors acknowledge the financial support by the Federal Ministry of Education and Research of Germany and by Sächsische Staatsministerium für Wissenschaft, Kultur und Tourismus in the programme Center of Excellence for AI-research „Center for Scalable Data Analytics and Artificial Intelligence Dresden/Leipzig”, project identification number: SCADS24B

This project was funded by “ChemoTox-AI”, a project financed by the German Federal Ministry of Education and Research (BMBF) within the Computational Life Science line of funding (BMBF 031L0261).

Computations for this work were done (in part) using resources of the Leipzig University Computing Centre.

The authors gratefully acknowledge the computing time made available to them on the high-performance computer at the Technical University Dresden (NHR center). This center is jointly supported by the Federal Ministry of Education and Research and the state governments participating in the NHR network (www.nhr-verein.de/unsere-partner).

### Competing interests

Markus Scholz received funding from Pfizer Inc. and has an ongoing cooperation with Owkin for projects not related to this research.

### Ethics approval and consent to participate

Data were obtained from studies of the German High-Grade Non-Hodgkin’s Lymphoma Study Group and were collected in the framework of the NHL-B study (Pfreundschuh, 2004a,b). All patients had given informed consent and studies were approved by responsible ethics committees and were carried out in accordance with the principles of good clinical practice and the declaration of Helsinki. Details on ethics committees and reference numbers can be found in the respective publications of the studies used (Pfreundschuh, 2004a,b).

### Consent for publication

Not applicable

### Data availability

All model relevant data are within the manuscript and its Supporting Information files.

### Materials availability

Not applicable

### Code availability

A demonstrator for learning an individual patients dynamics is provided under https://github.com/mariesteinacker/NARX-Hematox.

### Author contribution

**Marie Steinacker:** Conceived and designed the experiments, Performed the experiments, Analyzed and interpreted the data, Wrote the paper.

**Yuri Kheifetz:** Conceived and designed the experiments, Analyzed and interpreted the data.

**Markus Scholz:** Conceived and designed the experiments, Analyzed and interpreted the data, Wrote the paper.

## Notes

https://github.com/mariesteinacker/NARX-Hematox

## References

An G (1996) The Effects of Adding Noise During Backpropagation Training on a Generalization Performance. Neural Computation 8(3):643–674. 10.1162/neco.1996.8.3.643, URL https://direct.mit.edu/neco/article/8/3/643-674/5975

Brown R (1956) EXPONENTIAL SMOOTHING FOR PREDICTING DEMAND. Cambridge, Massachusetts: Arthur D Little Inc p p. 15. URL http://legacy.library.ucsf.edu/tid/dae94e00

Brown RG, Meyer RF (1961) The Fundamental Theorem of Exponential Smoothing. Operations Research 9(5):673–685. 10.1287/opre.9.5.673, URL https://pubsonline.informs.org/doi/10.1287/opre.9.5.673

Chen RTQ, Rubanova Y, Bettencourt J, et al (2018) Neural Ordinary Differential Equations. In: Bengio S, Wallach H, Larochelle H, et al (eds) Advances in Neural Information Processing Systems, vol 31. Curran Associates, Inc., URL https://proceedings.neurips.cc/paper_files/paper/2018/file/69386f6bb1dfed68692a24c8686939b9-Paper.pdf

Cho K, Van Merrienboer B, Gulcehre C, et al (2014) Learning Phrase Representations using RNN Encoder–Decoder for Statistical Machine Translation. In: Proceedings of the 2014 Conference on Empirical Methods in Natural Language Processing (EMNLP). Association for Computational Linguistics, Doha, Qatar, pp 1724–1734, 10.3115/v1/D14-1179, URL http://aclweb.org/anthology/D14-1179

Choi E, Bahadori MT, Schuetz A, et al (2016) Doctor AI: Predicting Clinical Events via Recurrent Neural Networks. JMLR Workshop Conf Proc 56:301–318

Elting LS, Rubenstein EB, Martin CG, et al (2001) Incidence, Cost, and Outcomes of Bleeding and Chemotherapy Dose Modification Among Solid Tumor Patients With Chemotherapy-Induced Thrombocytopenia. JCO 19(4):1137–1146. 10.1200/JCO.2001.19.4.1137, URL https://ascopubs.org/doi/10.1200/JCO.2001.19.4.1137

Esteban C, Staeck O, Baier S, et al (2016) Predicting Clinical Events by Combining Static and Dynamic Information Using Recurrent Neural Networks. pp 93–101, 10.1109/ICHI.2016.16

Friberg LE, Karlsson MO (2003) Mechanistic models for myelosuppression. Investigational New Drugs 21(2):183–194. 10.1023/A:1023573429626, URL http://link.springer.com/10.1023/A:1023573429626

Friberg LE, Henningsson A, Maas H, et al (2002) Model of Chemotherapy-Induced Myelosuppression With Parameter Consistency Across Drugs. JCO 20(24):4713–4721. 10.1200/JCO.2002.02.140, URL http://ascopubs.org/doi/10.1200/JCO.2002.02.140

Goodfellow I, Bengio Y, Courville A (2016) Deep Learning. MIT Press, URL https://www.deeplearningbook.org/

Henrich A, Joerger M, Kraff S, et al (2017) Semimechanistic Bone Marrow Exhaustion Pharmacokinetic/Pharmacodynamic Model for Chemotherapy-Induced Cumulative Neutropenia. J Pharmacol Exp Ther 362(2):347–358. 10.1124/jpet.117.240309, URL http://jpet.aspetjournals.org/lookup/doi/10.1124/jpet.117.240309

Holt CC (2004) Forecasting seasonals and trends by exponentially weighted moving averages. International Journal of Forecasting 20(1):5–10. 10.1016/j.ijforecast.2003.09.015, URL https://linkinghub.elsevier.com/retrieve/pii/S0169207003001134

Hornik K, Stinchcombe M, White H (1989) Multilayer feedforward networks are universal approximators. Neural Networks 2(5):359–366. 10.1016/0893-6080(89)90020-8, URL https://www.sciencedirect.com/science/article/pii/0893608089900208

Kheifetz Y, Scholz M (2019) Modeling individual time courses of thrombopoiesis during multi-cyclic chemotherapy. PLoS Comput Biol 15(3):e1006775. 10.1371/journal.pcbi.1006775, URL https://dx.plos.org/10.1371/journal.pcbi.1006775

Kheifetz Y, Scholz M (2021) Individual prediction of thrombocytopenia at next chemotherapy cycle: Evaluation of dynamic model performances. Br J Clin Pharmacol p bcp.14722. 10.1111/bcp.14722, URL https://onlinelibrary.wiley.com/doi/10.1111/bcp.14722

Kleissl M, Drews L, Heyder BB, et al (2023) SimbaML: Connecting Mechanistic Models and Machine Learning with Augmented Data. In: Maughan K, Liu R, Burns TF (eds) The First Tiny Papers Track at ICLR 2023, Tiny Papers @ ICLR 2023, Kigali, Rwanda, May 5, 2023. OpenReview.net, URL https://openreview.net/pdf?id=1wtUadpmVzu

Krogh A, Hertz J (1991) A Simple Weight Decay Can Improve Generalization. In: Moody J, Hanson S, Lippmann RP (eds) Advances in Neural Information Processing Systems, vol 4. Morgan-Kaufmann, URL https://proceedings.neurips.cc/paper/1991/file/8eefcfdf5990e441f0fb6f3fad709e21-Paper.pdf

LeCun Y, Denker JS, Solla SA (1990) Optimal brain damage. In: Advances in neural information processing systems, pp 598–605

Li K, Daniels J, Liu C, et al (2020) Convolutional Recurrent Neural Networks for Glucose Prediction. IEEE J Biomed Health Inform 24(2):603–613. 10.1109/JBHI.2019.2908488, URL https://ieeexplore.ieee.org/document/8678399/

Mangas-Sanjuan V, Buil-Bruna N, Garrido MJ, et al (2015) Semimechanistic Cell-Cycle Type–Based Pharmacokinetic/Pharmacodynamic Model of Chemotherapy-Induced Neutropenic Effects of Diflomotecan under Different Dosing Schedules. J Pharmacol Exp Ther 354(1):55–64. 10.1124/jpet.115.223776, URL http://jpet.aspetjournals.org/lookup/doi/10.1124/jpet.115.223776

National Cancer Institute (NCI) (2009) Common Terminology Criteria for Adverse Events v3.0. URL https://ctep.cancer.gov/protocolDevelopment/electronic_applications/docs/ctcaev3.pdf

Pan SJ, Yang Q (2010) A Survey on Transfer Learning. IEEE Trans Knowl Data Eng 22(10):1345–1359. 10.1109/TKDE.2009.191, URL http://ieeexplore.ieee.org/document/5288526/

Pfreundschuh M (2004a) Two-weekly or 3-weekly CHOP chemotherapy with or without etoposide for the treatment of elderly patients with aggressive lymphomas: results of the NHL-B2 trial of the DSHNHL. Blood 104(3):634–641. 10.1182/blood-2003-06-2095, URL http://www.bloodjournal.org/cgi/doi/10.1182/blood-2003-06-2095

Pfreundschuh M (2004b) Two-weekly or 3-weekly CHOP chemotherapy with or without etoposide for the treatment of young patients with good-prognosis (normal LDH) aggressive lymphomas: results of the NHL-B1 trial of the DSHNHL. Blood 104(3):626–633. 10.1182/blood-2003-06-2094, URL http://www.bloodjournal.org/cgi/doi/10.1182/blood-2003-06-2094

Raissi M, Perdikaris P, Karniadakis G (2019) Physics-informed neural networks: A deep learning framework for solving forward and inverse problems involving nonlinear partial differential equations. Journal of Computational Physics 378:686–707. 10.1016/j.jcp.2018.10.045, URL https://linkinghub.elsevier.com/retrieve/pii/S0021999118307125

Scholz M, Gross A, Loeffler M (2010) A biomathematical model of human thrombopoiesis under chemotherapy. Journal of Theoretical Biology 264(2):287–300. 10.1016/j.jtbi.2009.12.032, URL https://linkinghub.elsevier.com/retrieve/pii/S0022519310000056

Siegelmann H, Horne B, Giles C (1997) Computational capabilities of recurrent NARX neural networks. IEEE Transactions on Systems, Man, and Cybernetics, Part B (Cybernetics) 27(2):208–215. 10.1109/3477.558801, URL https://ieeexplore.ieee.org/document/558801/

Srivastava N, Hinton G, Krizhevsky A, et al (2014) Dropout: A Simple Way to Prevent Neural Networks from Overfitting. Journal of Machine Learning Research 15(56):1929–1958. URL http://jmlr.org/papers/v15/srivastava14a.html

Steinacker M, Kheifetz Y, Scholz M (2023) Individual modelling of haematotoxicity with NARX neural networks: A knowledge transfer approach. Heliyon 9(7):e17890. 10.1016/j.heliyon.2023.e17890, URL https://linkinghub.elsevier.com/retrieve/pii/S2405844023050983

Wunderlich A, Kloess M, Reiser M, et al (2003) Practicability and acute haematological toxicity of 2- and 3-weekly CHOP and CHOEP chemotherapy for aggressive non-Hodgkin’s lymphoma: results from the NHL-B trial of the GermanHigh-Grade Non-Hodgkin’s Lymphoma Study Group (DSHNHL). Annals of Oncology 14(6):881–893. 10.1093/annonc/mdg249, URL https://linkinghub.elsevier.com/retrieve/pii/S0923753419635507

Zabbarov J, Witzke S, Kleissl M, et al (2024) Optimizing ODE-derived Synthetic Data for Transfer Learning in Dynamical Biological Systems. 10.1101/2024.03.25.586390, URL http://biorxiv.org/lookup/doi/10.1101/2024.03.25.586390

