## Supplementary material for "Predicting chemotherapy-induced thrombotoxicity by NARX neural networks and transfer learning": Online Resource 1

Title: Predicting chemotherapy-induced thrombotoxicity by NARX neural networks and transfer learning
Journal: Journal of Cancer Research and Clinical Oncology
Authors: Marie Steinacker*, Yuri Kheifetz, Markus Scholz
Affiliation: *Center for Scalable Data Analytics and Artificial Intelligence (ScaDS.AI) Dresden/Leipzig, Leipzig University, Leipzig, Germany

* corresponding author

### 1 Supplemental material

#### 1.1 NARX modeling

Nonlinear auto-regressive models with exogenous inputs (NARX) are designed for discrete time series modeling. They relate previous observations, predictions and exogenous inputs to the next time step of the time series. They can be described by the general recursive formula, Eqn. 1,

$$\hat{y}_t = \mathcal{F}\left(\hat{y}_{t-1}, \ldots, \hat{y}_{t-n_y}, u_{t-1}, \ldots, u_{t-n_u}\right), \tag{1}$$

where $\hat{y}_t$ denotes the prediction made by the model at time $t$ and $u_t$ is the exogenous input at time $t$. The number of past predictions and exogenous information included in the model prediction is controlled via variables $n_y, n_u \in \mathbb{N}$. To start this recursive process, the first $n_y$ observations are required. The link function $F$ denotes an arbitrary nonlinear function of past observations $\hat{y}$ and inputs $u$.

In our work, we consider neural networks as link functions, further called NARX neural networks. The nonlinearity in these networks stems from a non-linear activation function. All NARX neural networks are recurrent networks (RNNs), and the recurrent connection between the output of the network and the input is formed by a so-called tapped delay line (Siegelmann et al., 1997). In general, any type of neural network can be used to create a NARX neural network. In our paper we compare selected network architectures.

First, we consider a simple adaptation of a feed-forward network as proposed by Steinacker et al., 2023, further called ARX-FNN. In a feed-forward network, information only flows in one direction through the network, from the input nodes through potential so-called hidden layers of nodes, which are not directly connected to the input or the output nodes of the network. In our situation, the input vector to the network comprise past measured or regressed platelet counts and past information of chemotherapy applications. We use a memory time of 21 days of past platelet counts, and 28 days of past treatment information. This input vector is processed by one hidden layer with four neurons with a sigmoid activation function, and then, passed to the output node, which is the prediction of the platelet count for the next day. This output is then re-used via the tapped delay line as a network input for the prediction of the next day. In a so-called warm-up phase of the network, initial values are required as inputs. For this purpose, we assume a stable steady state prior to treatment which is estimated by the observed values at baseline.

We compare the simple ARX-FNN with a more complex RNN model, namely a gated-recurrent unit (GRU) network (Cho et al., 2014) used as the inner network, further called ARX-GRU. In addition to the network weights, traditional RNNs have hidden states which store the past contextual information for a seen input sequence. The function applied by a GRU to its input can be described by the following equations (Cho et al., 2014),

$$z_t = \sigma(W^z x_t + U^z h_{t-1}), \tag{2}$$
$$r_t = \sigma(W^r x_t + U^r h_{t-1}), \tag{3}$$
$$h'_t = \tanh(W x_t + U(r_t \odot h_{t-1})), \tag{4}$$
$$h_t = z_t \odot h_{t-1} + (1 - z_t) \odot h'_t, \tag{5}$$

where $x_t$ is the current input to a unit and $h_t$ the current hidden state to be computed. The activation of this unit is computed with an update gate $z_t$ and a reset gate $r_t$, which depend on the input and the previous hidden state. The update gate controls the amount of previous information transferred to the current state. The reset gate is used to control the memory $h'_t$. $U$ and $W$ denote respective weight matrices to learn and $\sigma$ is the sigmoid function. This architecture allows the network to keep relevant information for a longer period of time, and respectively, discard irrelevant past information.

In the case of the ARX-GRU, Eqn. 1 can be simplified to

$$\hat{y}_t = F(\hat{y}_{t-1}, u_{t-1}), \tag{6}$$

as the kept information about the past is handled by the GRU network internally. Here, we use a network with a GRU hidden layer of 64 nodes. While this is much larger then the FNN network, the input size is much smaller. We chose hidden layer sizes via hyperparameter optimization, based on fitting a test set population . Also, we tested a simple RNN as well as an long-short term memory (LSTM) network (Hochreiter and Schmidhuber, 1997) as an inner network, but the ARX-GRU outperformed both, while being much simpler then an LSTM (results not shown).

An important distinction between both architectures is the memory handling. The ARX-FNN is the simpler architecture, where the memorized time period is fixed. Network predictions are easier to comprehend for this architecture. With the ARX-GRU, the network handles the kept memory internally, and only the regressed prediction and treatment information of the previous day is explicitly included in the prediction. Therefore, the network is larger then the ARX-FNN and has a more complex internal structure reducing its explainability.

#### 1.2 Semi-mechanistic model

We use a model of myelosuppresion proposed by Friberg et al. for transfer learning and direct competitor of our NARX approach. The model is described by the following set of differential equations,

$$\frac{d}{dt}Prol = k_{tr} \cdot Prol \cdot (1 - E_{drug}) \left(\frac{Circ_0}{Circ}\right)^{\gamma} - k_{tr} \cdot Prol, \tag{7}$$

$$\frac{d}{dt}Transit_1 = k_{tr} \cdot Prol - k_{tr} \cdot Transit_1, \tag{8}$$

$$\frac{d}{dt}Transit_2 = k_{tr} \cdot Transit_1 - k_{tr} \cdot Transit_2, \tag{9}$$

$$\frac{d}{dt}Transit_3 = k_{tr} \cdot Transit_2 - k_{tr} \cdot Transit_3, \tag{10}$$

$$\frac{d}{dt}Circ = k_{tr} \cdot Transit_3 - k_{Circ} \cdot Circ, \tag{11}$$

where the cell count of circulating blood cells to be described, is denoted by $Circ$. It's steady state value is $Circ_0$, and the cell count of stem cells in the proliferation compartment is denoted by $Prol$. The maturation of proliferating cells into circulating blood cells is modeled by three transit compartments $Transit_{1-3}$. The influence of chemotherapy is introduced by an effect function $E_{drug}$, which depends on the drug concentration in the body. The rate $k_{tr}$ describing proliferation and transition is assumed to be constant for both processes. It is obtained using the mean transit time $MTT$, i.e. $k_{tr} = 4/MTT$. The feedback between circulating cells and the proliferation compartment is controlled by parameter $\gamma$.

For the effect function $E_{drug}$, we assume a step-function. On days of cytotoxic drug applications, $E_{drug}$ is assumed as a constant on days of treatment, the dosage of treatment the patient received times an individual factor $E_s$. This dosage is normalized to what a patient of the same body surface area would receive for this treatment schedule, to be comparable between patients. On days without treatment, we assume no continued effect.

We utilize a population approach for individual calibrations of the semi-mechanistic model, ensuring that a parameter set calibrated to an individual patient does not deviate too far from the population average if only sparse data is available for this patient. The objective function $\mathcal{L}$ for the calibration has the following form

$$\mathcal{L} = \mathcal{L}_{data} + \mathcal{L}_{param}, \tag{12}$$

$$= \frac{1}{N}\sum_{i=1}^{N}(y_i - \hat{y}_i)^2 + \sum_j \frac{(\ln(p_j) - \ln(p_j^0))^2}{\sigma^2}, \tag{13}$$

with parameters $p_j$ and their population averages $p_j^0$, target data $y$ and respective model predictions $\hat{y}$. The variable $\sigma$ controls the penalty of deviating from the population average. We set $\sigma$ to 5. In the literature, the following population averages are proposed: feedback $\gamma = 0.316$, transit time $MTT = 195\,\text{h}$ (Joerger et al., 2007) and steady state $C_0 = 270 \times 10^9$cells/l (Warny et al., 2019). Based on population analysis for the remaining parameters, we assume for the linear drug effect $E_{eff} = 2$.

#### 1.3 Methods to reduce overfitting

We employ multiple methods to reduce overfitting during training of individual networks. Depending on network architecture and training regimen, we apply and combine different approaches.

*Data Augmentation:* We use dynamics simulated by the semi-mechanistic model to support pre-training of individual networks. As explained in Steinacker et al., 2023, we consider virtual therapy scenarios for this purpose varying for example doses and cycle duration. We also use such simulated scenarios for model validation. To speed up the time for pre-training, we start with network weights obtained for an average patient. This also stabilizes the learning in cases where the semi-mechanistic model proposes extreme parameter values.

*Pruning:* Over-parametrization is a well-known problem of neural network models. Pruning describes the removal of nodes or weights from the network to reduce model size and to improve generalization performance (LeCun et al., 1990). We employ pruning throughout all learning frameworks. Dropout is a form of non-permanent pruning, where weights are randomly removed from a network with a specified frequency (Srivastava et al., 2014). In our learning approach without transfer learning (TL), we apply dropout to the first training stage of the networks. Hereby, we obtain a simpler model in the first stage which roughly describes patient's time series and preventing overfitting as best as possible. However, it was necessary to fine-tune models in a second step by retraining with a lower learning rate to obtain acceptable final prediction performances.

*Regularization:* A popular way to reduce overfitting in neural networks is weight decay (Krogh and Hertz, 1991). Here, we add the squared sum of network weights times a coupling factor to the objective function to penalize non-zero weights. This can be applied also to only layers of the network, so only weights of a specified subset of network layers are included in the regularization term. Here, we apply this regularization to the output layer of the ARX-GRU models, with a coupling factor to the objective function of 0.01. For the ARX-FNN, we apply a modified regularization term, which is the sum of the root of absolute values of weights multiplied with 0.0001. Together with pruning, this specific regularization approach yield better results and more parsimonious models. Both the type of regularization and the coupling factors were determined via hyperparameter optimization.

*Exponential smoothing:* For sparse training data, neural networks tend to predict noise if the objective function primarily optimizes the agreement of predictions and data. We therefore decide to impose exponential smoothing (R. Brown, 1956; R. G. Brown and Meyer, 1961; Holt, 2004) in to our models, which acts as a low-pass noise filter. We apply it to the network output in the following way:

$$\hat{y}'_0 = \hat{y}_0, \tag{14}$$

$$\hat{y}'_i = \alpha \hat{y}_i + (1-\alpha)\hat{y}'_{i-1}, \tag{15}$$

with smoothing factor $\alpha \in [0,1]$, implemented as a trainable hyperparameter of the model. While $\hat{y}'$ denotes the prediction, the regressed variable $\hat{y}$ is used in the autoregressive feedback loop of the network.

*Smoothness penalty:* To force the model to apply exponential smoothing, we incorporate a smoothness penalty into the objective function as follows,

$$\mathcal{L}_{smooth} = \frac{1}{N}\sum_{i=1}^{N}(\hat{y}_i - \frac{\sum_{j=-k}^{k}\hat{y}_{i+j}}{2k+1})^2, \tag{16}$$

with model output $\hat{y}$ (including exponential smoothing) and $2k+1$ included neighboring outputs used for smoothing. During hyper-parameter optimization, we found

$k = 2$, and a factor of 0.01 for coupling with the overall objective function.

*Addition of a noise term to network input:* Another established method to avoid overfitting for small data-sets is the addition of a noise term to the training data, which can be seen as a form of data augmentation (Goodfellow et al., 2016). It encourages the neural network output to be a smooth function of its input and weights, respectively (An, 1996). We implement an additive Gaussian noise to the scaled network input via the tensorflow (Martín Abadi et al., 2015) function `tf.keras.layers.GaussianNoise` with a standard deviation of 0.2. This noise adding is only active in the training stages, where dynamics of real patient data are learned.

### References


An, G. (1996). The Effects of Adding Noise During Backpropagation Training on a Generalization Performance. *Neural Computation*, *8*(3), 643–674. https://doi.org/10.1162/neco.1996.8.3.643

Brown, R. (1956). EXPONENTIAL SMOOTHING FOR PREDICTING DEMAND. *Cambridge, Massachusetts: Arthur D. Little Inc.*, p. 15. http://legacy.library.ucsf.edu/tid/dae94e00

Brown, R. G., & Meyer, R. F. (1961). The Fundamental Theorem of Exponential Smoothing. *Operations Research*, *9*(5), 673–685. https://doi.org/10.1287/opre.9.5.673

Cho, K., Van Merrienboer, B., Gulcehre, C., Bahdanau, D., Bougares, F., Schwenk, H., & Bengio, Y. (2014). Learning Phrase Representations using RNN Encoder–Decoder for Statistical Machine Translation. *Proceedings of the 2014 Conference on Empirical Methods in Natural Language Processing (EMNLP)*, 1724–1734. https://doi.org/10.3115/v1/D14-1179

Friberg, L. E., Henningsson, A., Maas, H., Nguyen, L., & Karlsson, M. O. (2002). Model of Chemotherapy-Induced Myelosuppression With Parameter Consistency Across Drugs. *JCO*, *20*(24), 4713–4721. https://doi.org/10.1200/JCO.2002.02.140

Goodfellow, I., Bengio, Y., & Courville, A. (2016). *Deep Learning*. MIT Press. https://www.deeplearningbook.org/

Hochreiter, S., & Schmidhuber, J. (1997). Long Short-Term Memory. *Neural Computation*, *9*(8), 1735–1780. https://doi.org/10.1162/neco.1997.9.8.1735

Holt, C. C. (2004). Forecasting seasonals and trends by exponentially weighted moving averages. *International Journal of Forecasting*, *20*(1), 5–10. https://doi.org/10.1016/j.ijforecast.2003.09.015

Joerger, M., Huitema, A. D., Richel, D. J., Dittrich, C., Pavlidis, N., Briasoulis, E., Vermorken, J. B., Strocchi, E., Martoni, A., Sorio, R., Sleeboom, H. P., Izquierdo, M. A., Jodrell, D. I., Calvert, H., Boddy, A. V., Hollema, H., Féty, R., Van Der Vijgh, W. J., Hempel, G., … Schellens, J. H. (2007). Population Pharmacokinetics and Pharmacodynamics of Paclitaxel and Carboplatin in Ovarian Cancer Patients: A Study by the European Organization for Research and Treatment of Cancer-Pharmacology and Molecular Mechanisms Group and New Drug Development Group. *Clinical Cancer Research*, *13*(21), 6410–6418. https://doi.org/10.1158/1078-0432.CCR-07-0064

Krogh, A., & Hertz, J. (1991). A Simple Weight Decay Can Improve Generalization. In J. Moody, S. Hanson, & R. P. Lippmann (Eds.), *Advances in Neural Information Processing Systems* (Vol. 4). Morgan-Kaufmann. https://proceedings.neurips.cc/paper/1991/file/8eefcfdf5990e441f0fb6f3fad709e21-Paper.pdf

LeCun, Y., Denker, J. S., & Solla, S. A. (1990). Optimal brain damage. *Advances in neural information processing systems*, 598–605.

Martín Abadi, Ashish Agarwal, Paul Barham, Eugene Brevdo, Zhifeng Chen, Craig Citro, Greg S. Corrado, Andy Davis, Jeffrey Dean, Matthieu Devin, Sanjay Ghemawat, Ian Goodfellow, Andrew Harp, Geoffrey Irving, Michael Isard, Jia, Y., Rafal Jozefowicz, Lukasz Kaiser, Manjunath Kudlur, … Xiaoqiang Zheng. (2015). TensorFlow: Large-Scale Machine Learning on Heterogeneous Systems. https://www.tensorflow.org/

Siegelmann, H., Horne, B., & Giles, C. (1997). Computational capabilities of recurrent NARX neural networks. *IEEE Transactions on Systems, Man, and Cybernetics, Part B (Cybernetics)*, *27*(2), 208–215. https://doi.org/10.1109/3477.558801

Srivastava, N., Hinton, G., Krizhevsky, A., Sutskever, I., & Salakhutdinov, R. (2014). Dropout: A Simple Way to Prevent Neural Networks from Overfitting. *Journal of Machine Learning Research*, *15*(56), 1929–1958. http://jmlr.org/papers/v15/srivastava14a.html

Steinacker, M., Kheifetz, Y., & Scholz, M. (2023). Individual modelling of haematotoxicity with NARX neural networks: A knowledge transfer approach. *Heliyon*, *9*(7), e17890. https://doi.org/10.1016/j.heliyon.2023.e17890

Warny, M., Helby, J., Birgens, H. S., Bojesen, S. E., & Nordestgaard, B. G. (2019). Arterial and venous thrombosis by high platelet count and high hematocrit: 108 521 individuals from the Copenhagen General Population Study. *Journal of Thrombosis and Haemostasis*, *17*(11), 1898–1911. https://doi.org/10.1111/jth.14574
