## Supplementary material for "Predicting chemotherapy-induced thrombotoxicity by NARX neural networks and transfer learning": Online Resource 2

Journal: Journal of Cancer Research and Clinical Oncology

Authors: Marie Steinacker*, Yuri Kheifetz, Markus Scholz
Affiliation: *Center for Scalable Data Analytics and Artificial Intelligence (ScaDS.AI) Dresden/Leipzig, Leipzig University, Leipzig, Germany

* corresponding author

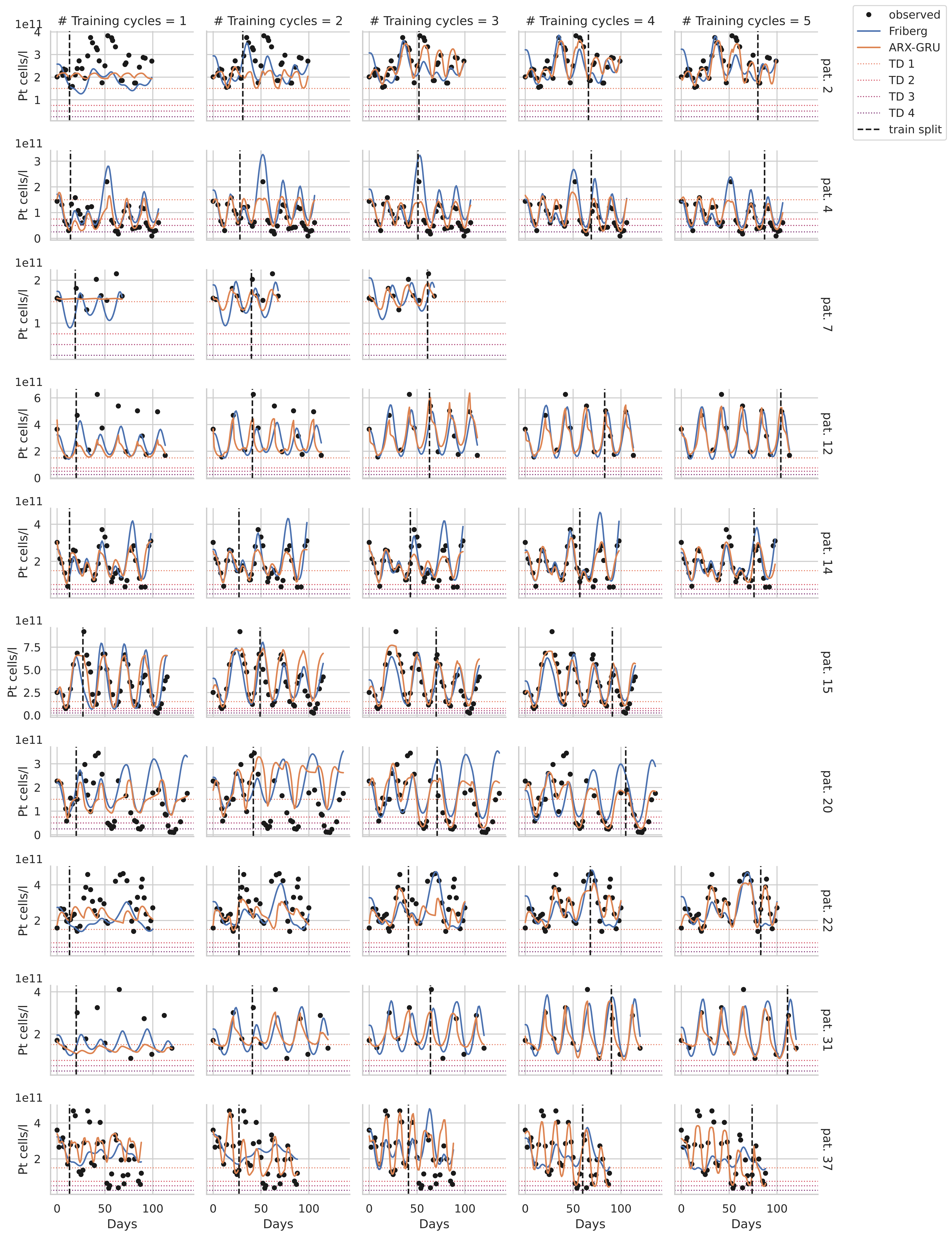

### Training cycles = 1
### Training cycles = 2
### Training cycles = 3
### Training cycles = 4
### Training cycles = 5
observed
Friberg
ARX-GRU
TD 1
TD 2
TD 3
TD 4
train split
Pt cells/l
1e11
Days
pat. 2
pat. 4
pat. 7
pat. 12
pat. 14
pat. 15
pat. 20
pat. 22
pat. 31
pat. 37

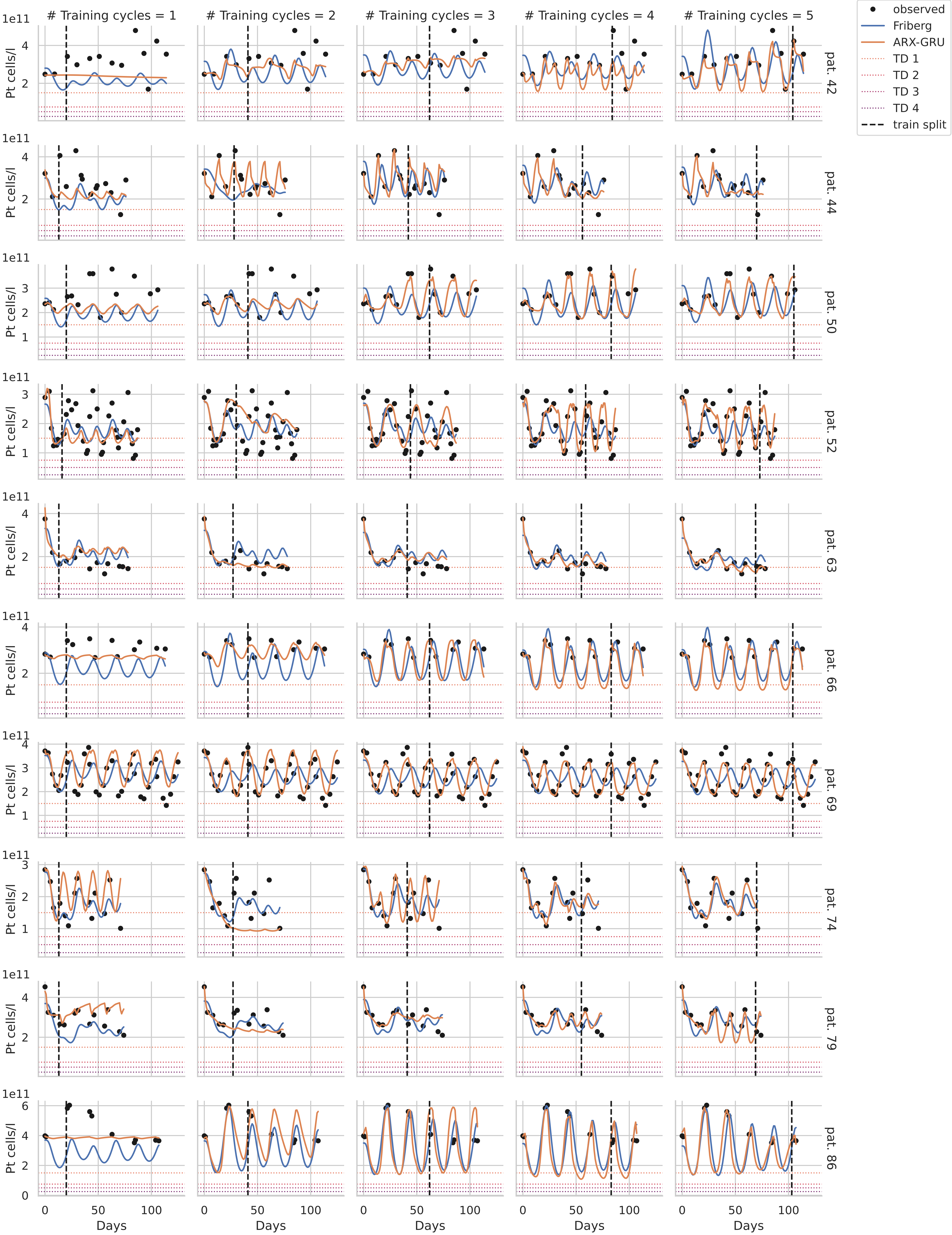

### Training cycles = 1
### Training cycles = 2
### Training cycles = 3
### Training cycles = 4
### Training cycles = 5
observed
Friberg
ARX-GRU
TD 1
TD 2
TD 3
TD 4
train split
Pt cells/l
1e11
Days
pat. 42
pat. 44
pat. 50
pat. 52
pat. 63
pat. 66
pat. 69
pat. 74
pat. 79
pat. 86
0
50
100

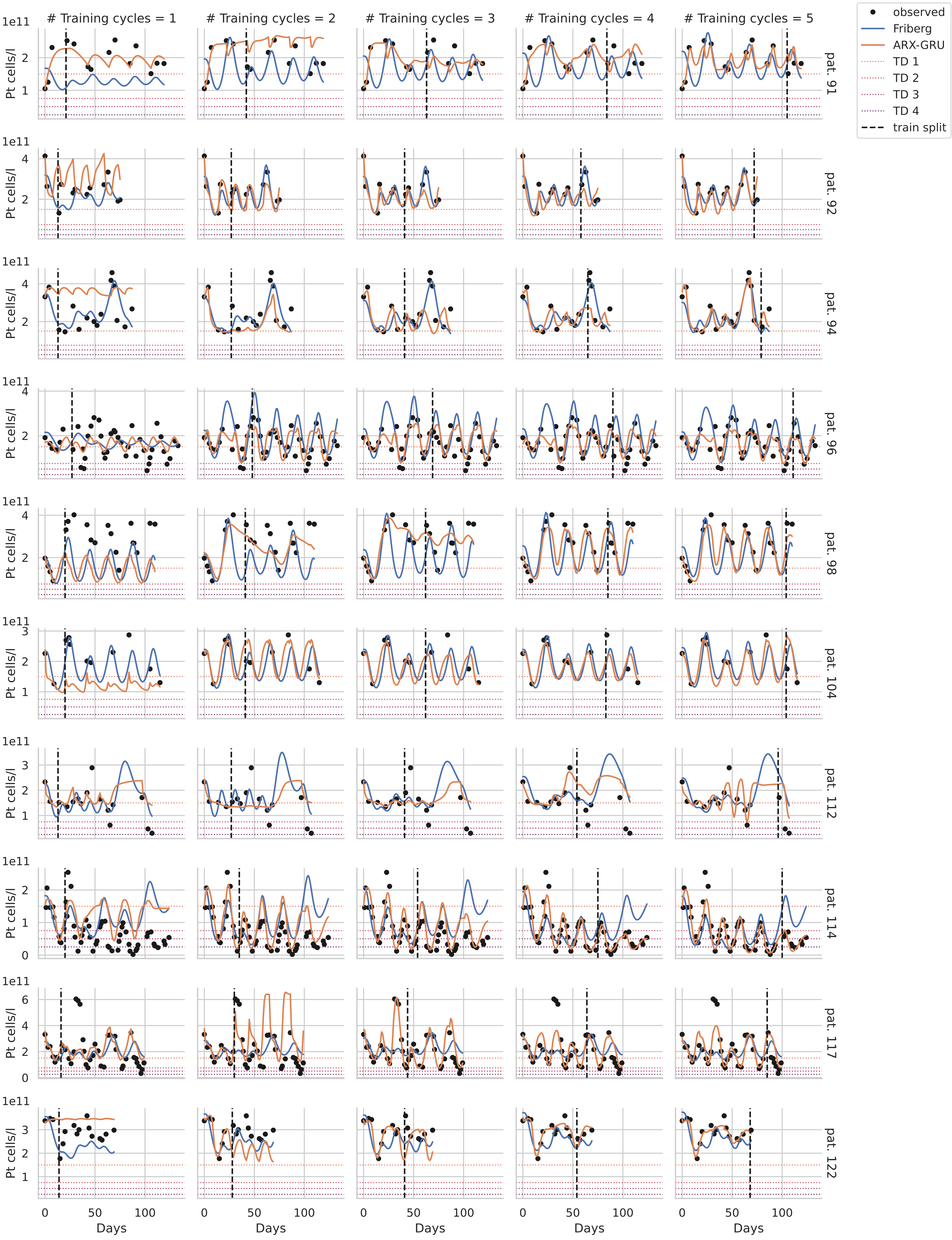

### Training cycles = 1
### Training cycles = 2
### Training cycles = 3
### Training cycles = 4
### Training cycles = 5
observed
Friberg
ARX-GRU
TD 1
TD 2
TD 3
TD 4
train split
Pt cells/l
1e11
pat. 91
pat. 92
pat. 94
pat. 96
pat. 98
pat. 104
pat. 112
pat. 114
pat. 117
pat. 122
Days

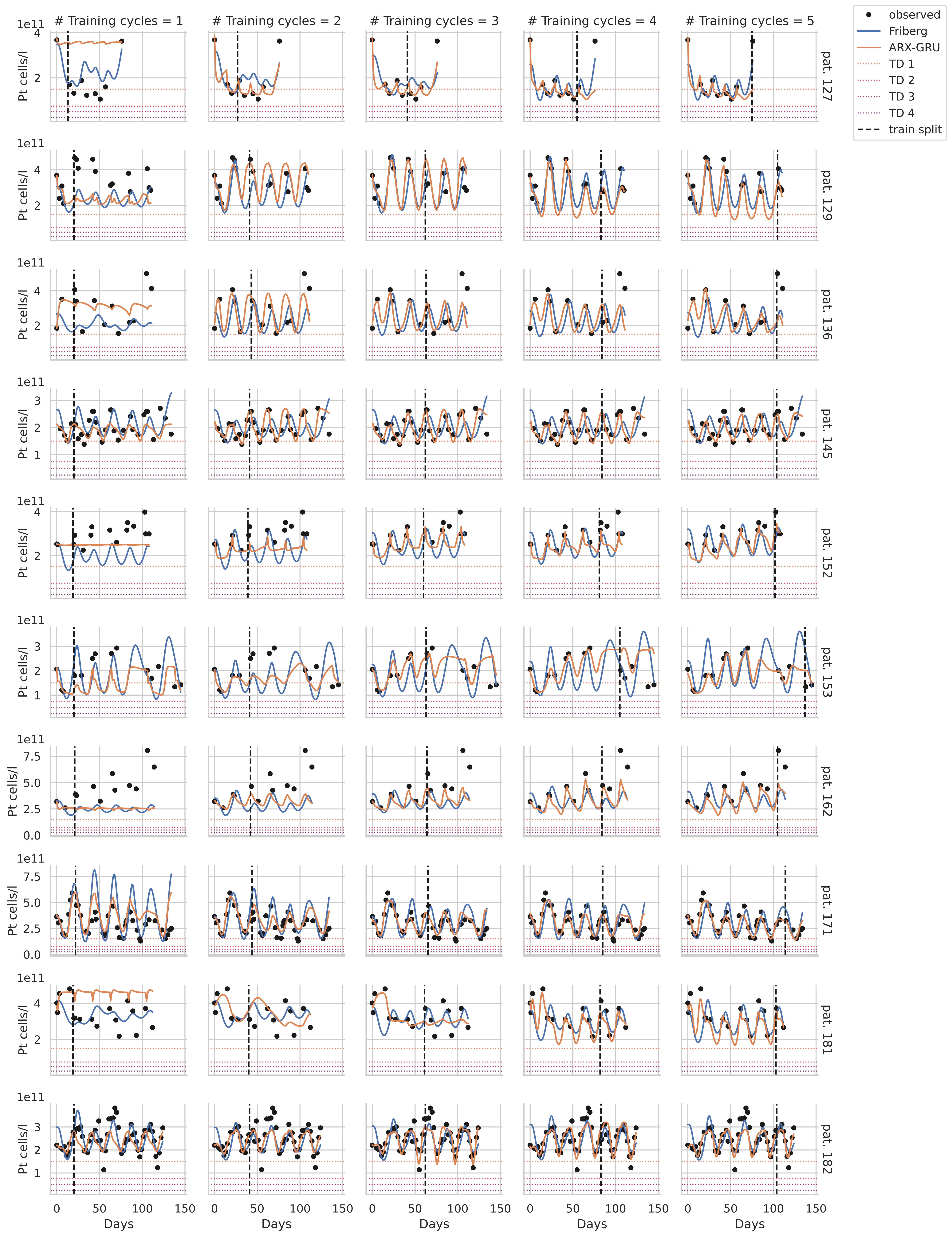

### Training cycles = 1
### Training cycles = 2
### Training cycles = 3
### Training cycles = 4
### Training cycles = 5
observed
Friberg
ARX-GRU
TD 1
TD 2
TD 3
TD 4
train split
Pt cells/l
1e11
Days
pat. 127
pat. 129
pat. 136
pat. 145
pat. 152
pat. 153
pat. 162
pat. 171
pat. 181
pat. 182

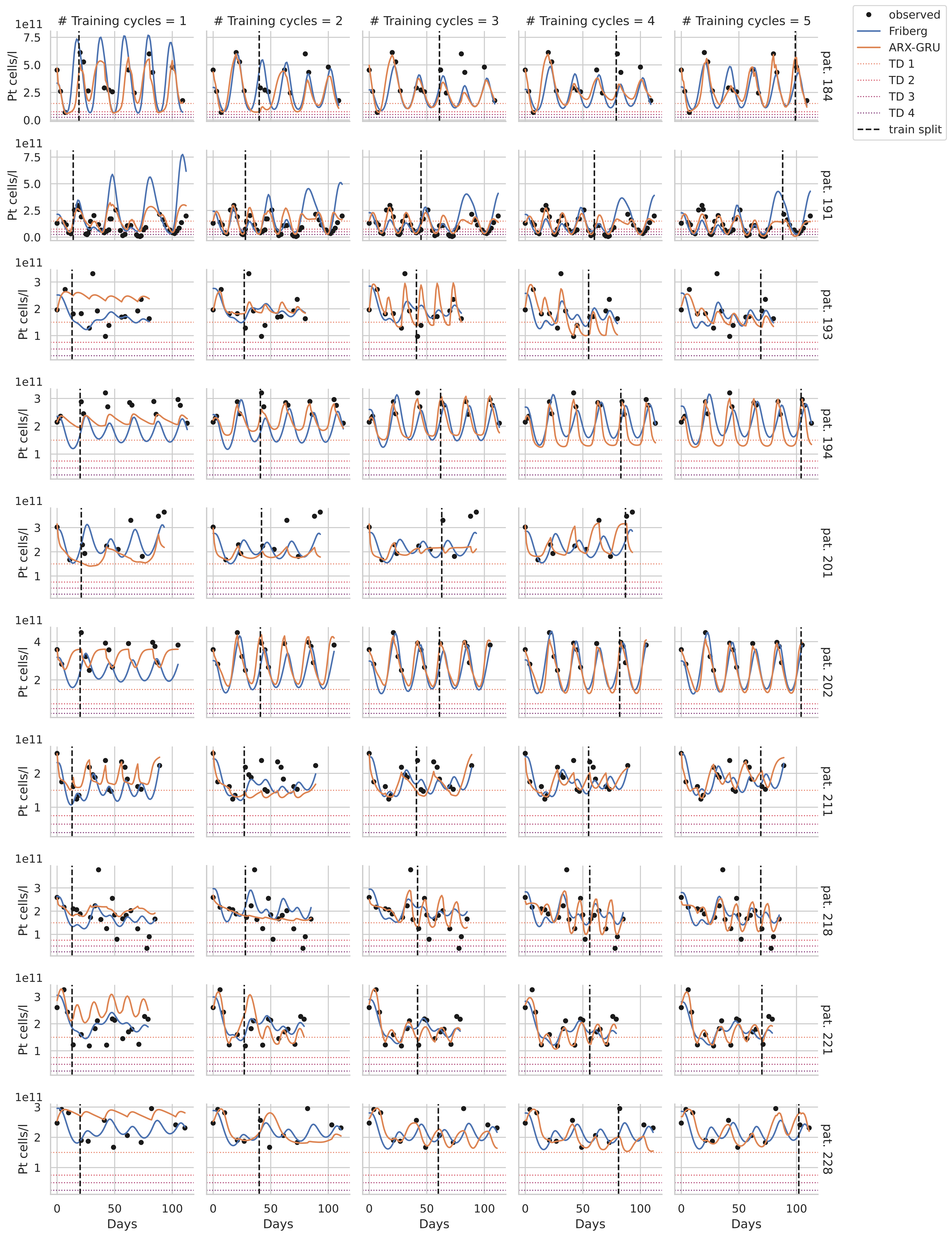

### Training cycles = 1
### Training cycles = 2
### Training cycles = 3
### Training cycles = 4
### Training cycles = 5
observed
Friberg
ARX-GRU
TD 1
TD 2
TD 3
TD 4
train split
Pt cells/l
1e11
pat. 184
pat. 191
pat. 193
pat. 194
pat. 201
pat. 202
pat. 211
pat. 218
pat. 221
pat. 228
Days

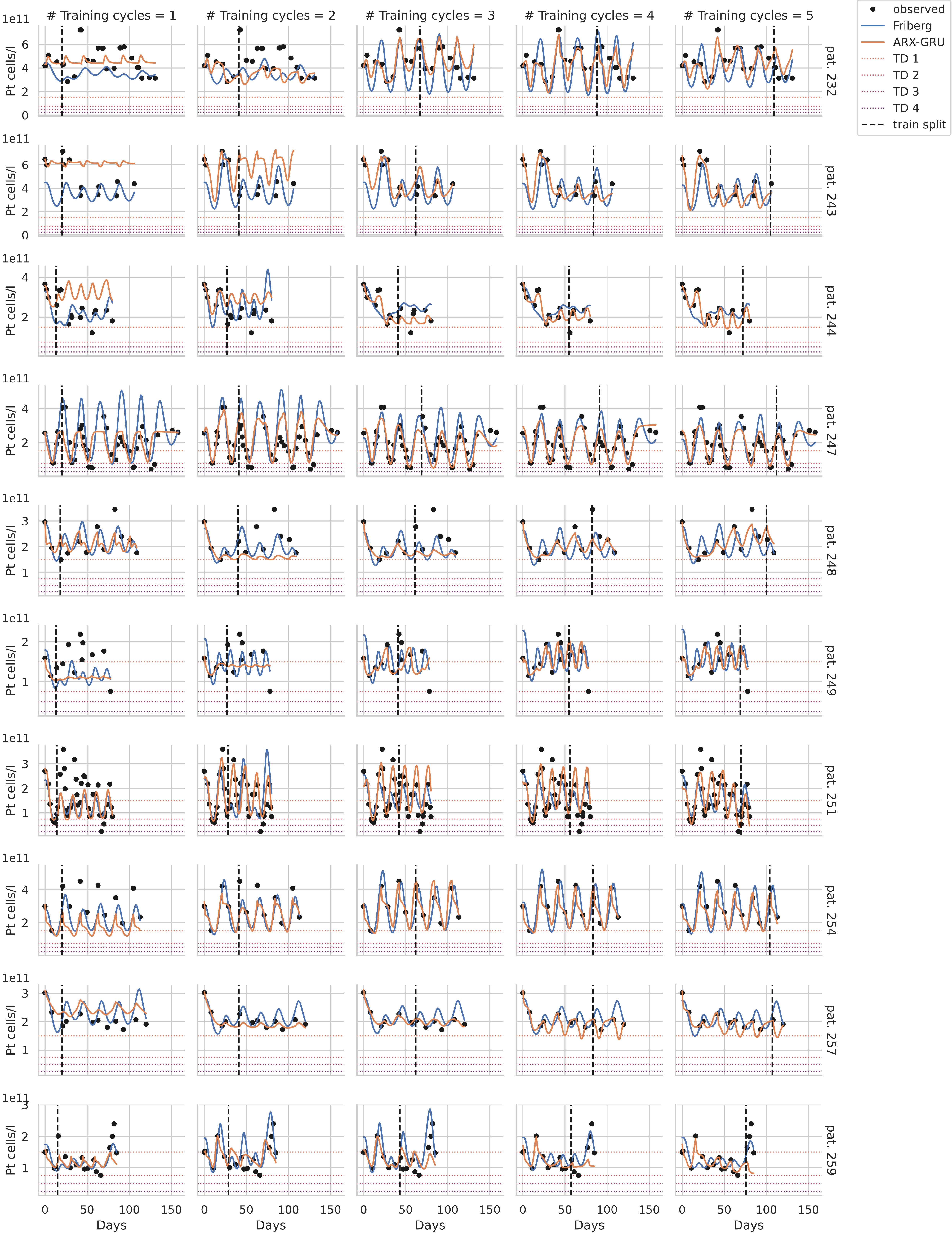

### Training cycles = 1
### Training cycles = 2
### Training cycles = 3
### Training cycles = 4
### Training cycles = 5
observed
Friberg
ARX-GRU
TD 1
TD 2
TD 3
TD 4
train split
Pt cells/l
1e11
pat. 232
pat. 243
pat. 244
pat. 247
pat. 248
pat. 249
pat. 251
pat. 254
pat. 257
pat. 259
Days

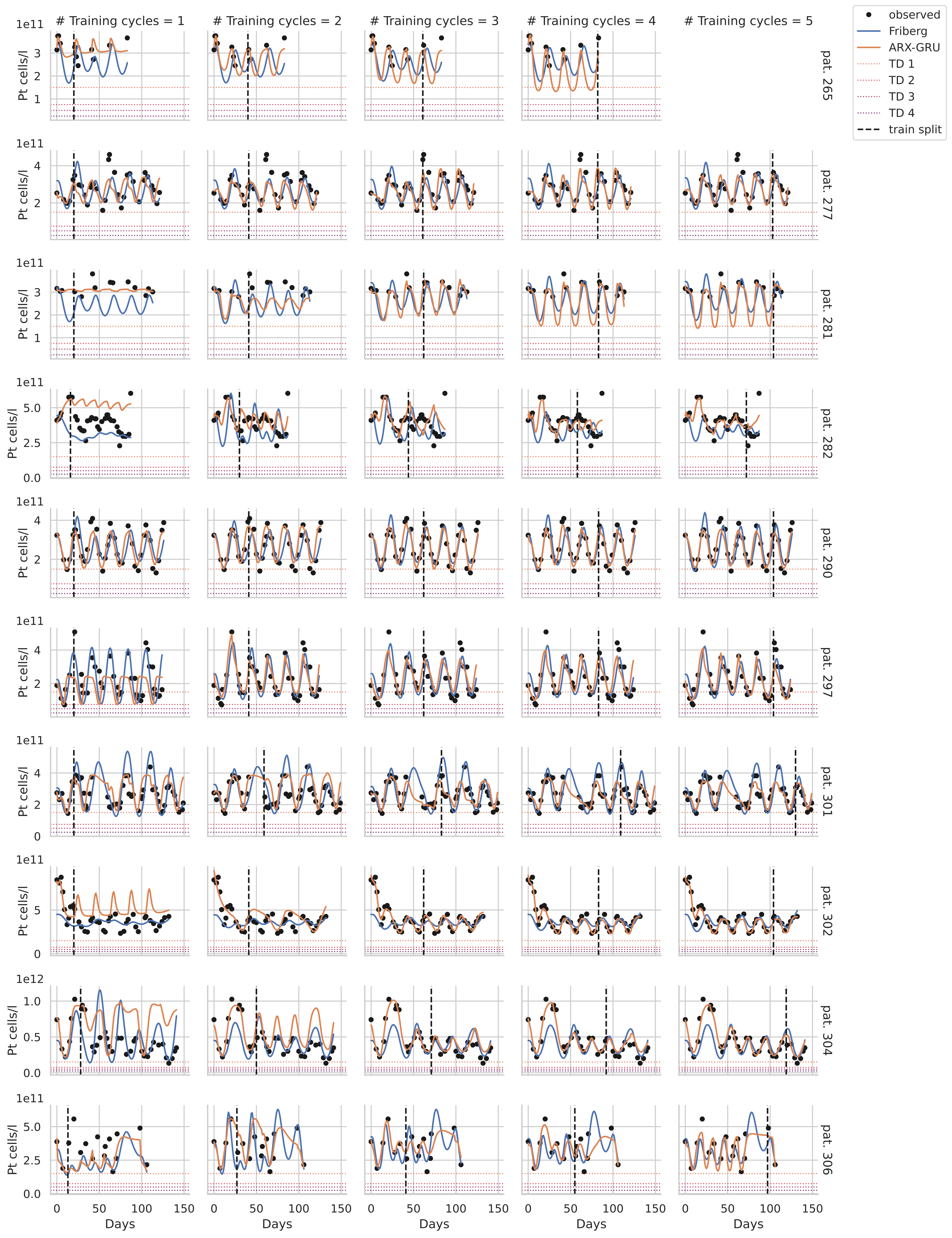

### Training cycles = 1
### Training cycles = 2
### Training cycles = 3
### Training cycles = 4
### Training cycles = 5
observed
Friberg
ARX-GRU
TD 1
TD 2
TD 3
TD 4
train split
Pt cells/l
Days
pat. 265
pat. 277
pat. 281
pat. 282
pat. 290
pat. 297
pat. 301
pat. 302
pat. 304
pat. 306

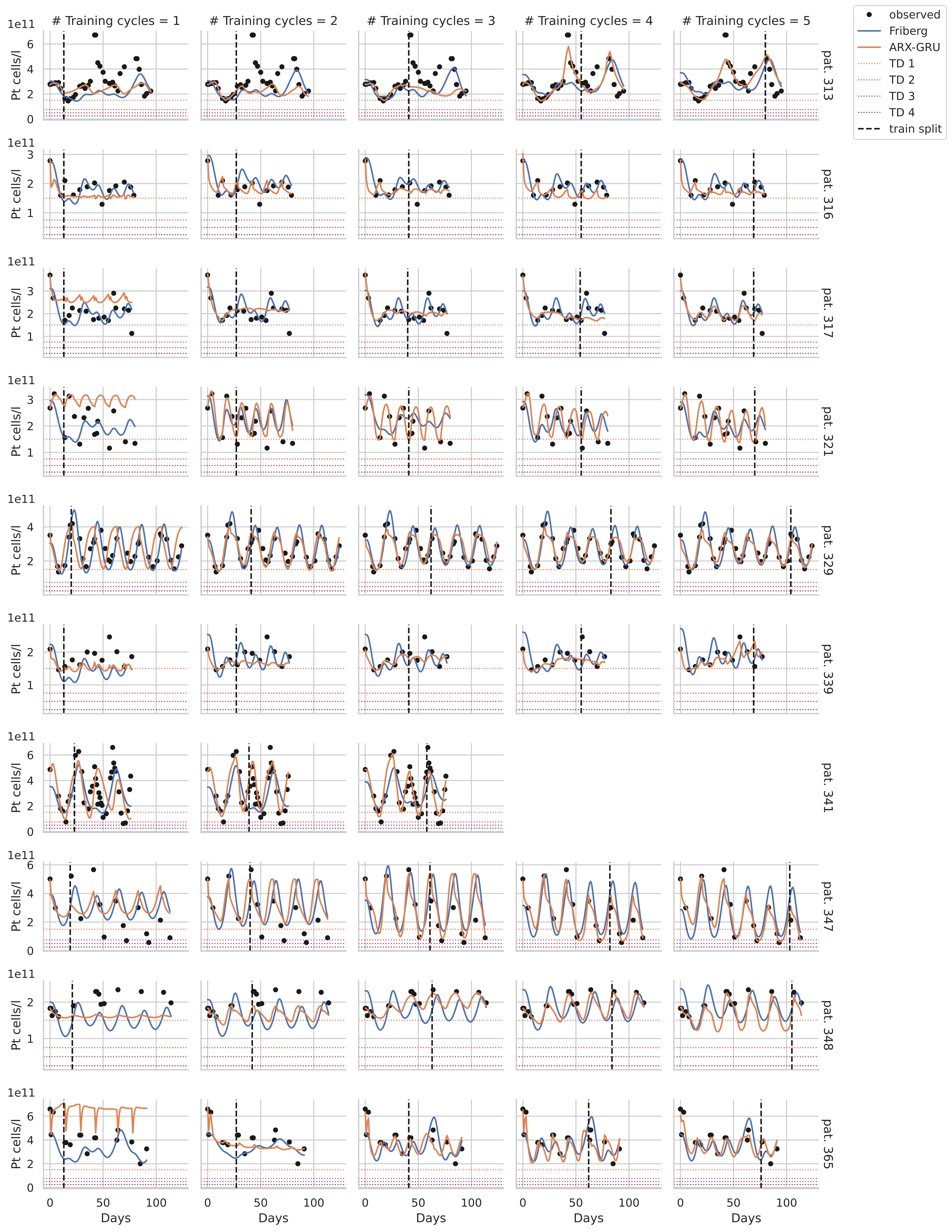

### Training cycles = 1
### Training cycles = 2
### Training cycles = 3
### Training cycles = 4
### Training cycles = 5
observed
Friberg
ARX-GRU
TD 1
TD 2
TD 3
TD 4
train split
Pt cells/l
1e11
pat. 313
pat. 316
pat. 317
pat. 321
pat. 329
pat. 339
pat. 341
pat. 347
pat. 348
pat. 365
Days

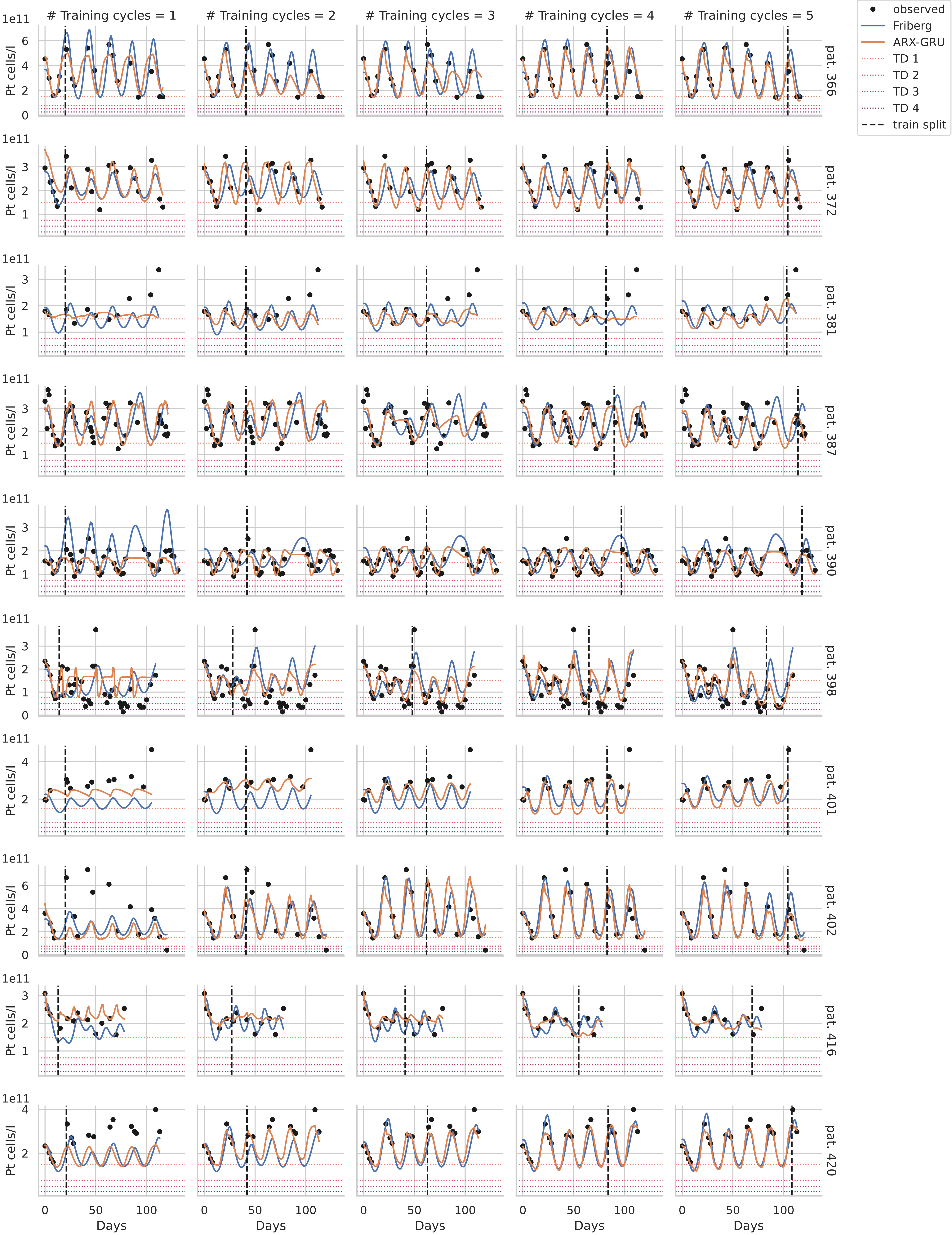

### Training cycles = 1
### Training cycles = 2
### Training cycles = 3
### Training cycles = 4
### Training cycles = 5
observed
Friberg
ARX-GRU
TD 1
TD 2
TD 3
TD 4
train split
pat. 366
pat. 372
pat. 381
pat. 387
pat. 390
pat. 398
pat. 401
pat. 402
pat. 416
pat. 420
Pt cells/l
1e11
Days

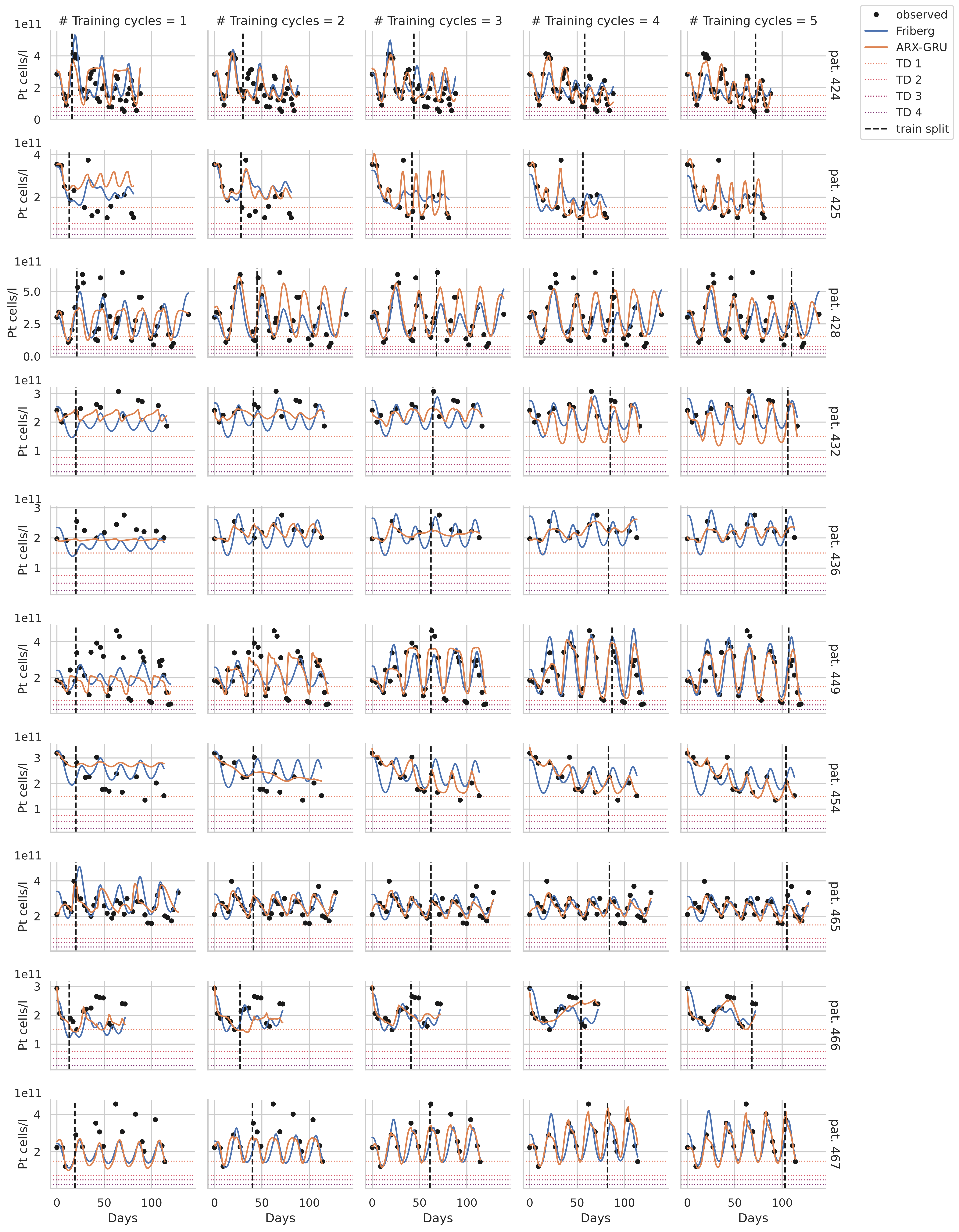

### Training cycles = 1
### Training cycles = 2
### Training cycles = 3
### Training cycles = 4
### Training cycles = 5
observed
Friberg
ARX-GRU
TD 1
TD 2
TD 3
TD 4
train split
Pt cells/l
1e11
pat. 424
pat. 425
pat. 428
pat. 432
pat. 436
pat. 449
pat. 454
pat. 465
pat. 466
pat. 467
Days

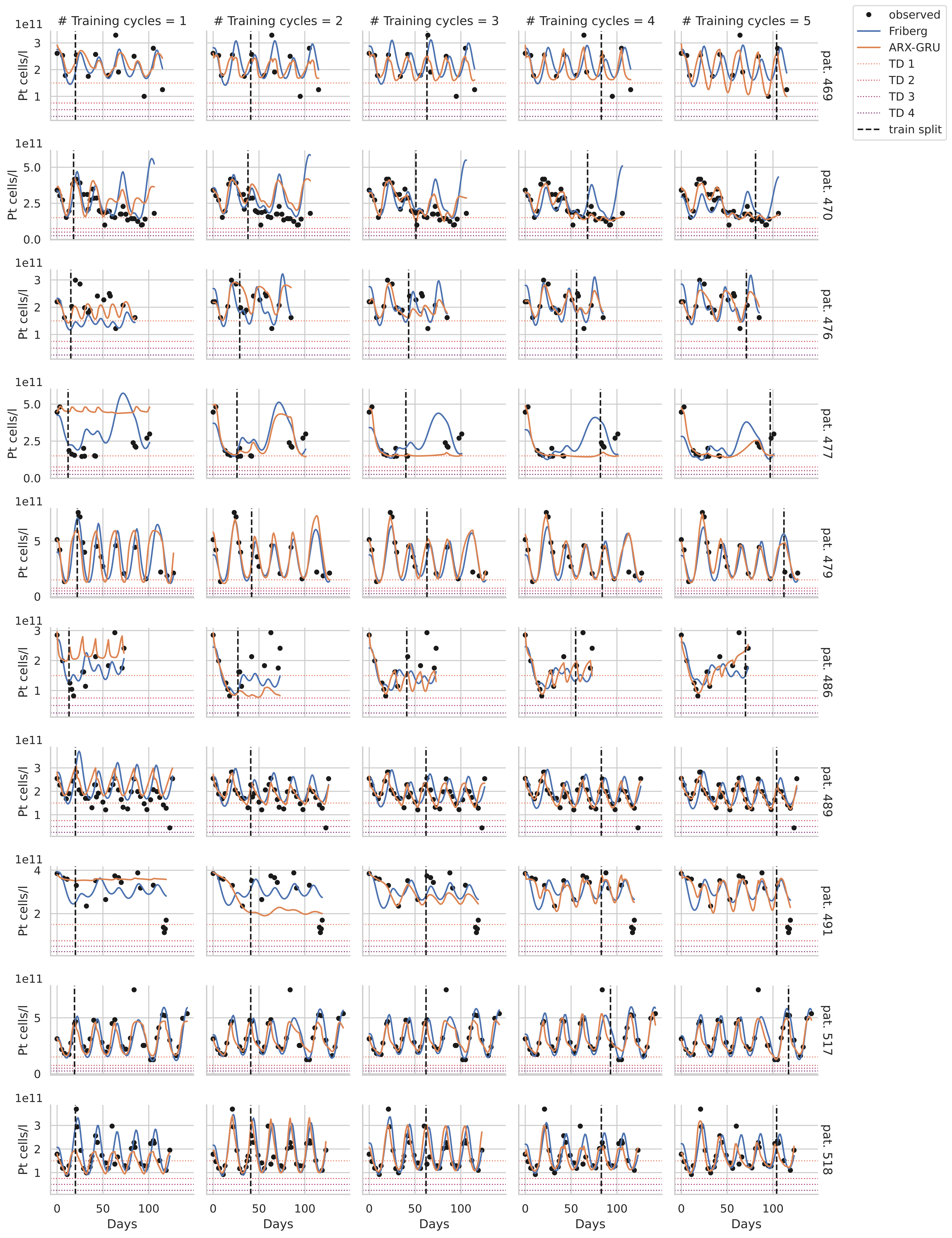

### Training cycles = 1
### Training cycles = 2
### Training cycles = 3
### Training cycles = 4
### Training cycles = 5
observed
Friberg
ARX-GRU
TD 1
TD 2
TD 3
TD 4
train split
Pt cells/l
1e11
pat. 469
pat. 470
pat. 476
pat. 477
pat. 479
pat. 486
pat. 489
pat. 491
pat. 517
pat. 518
Days

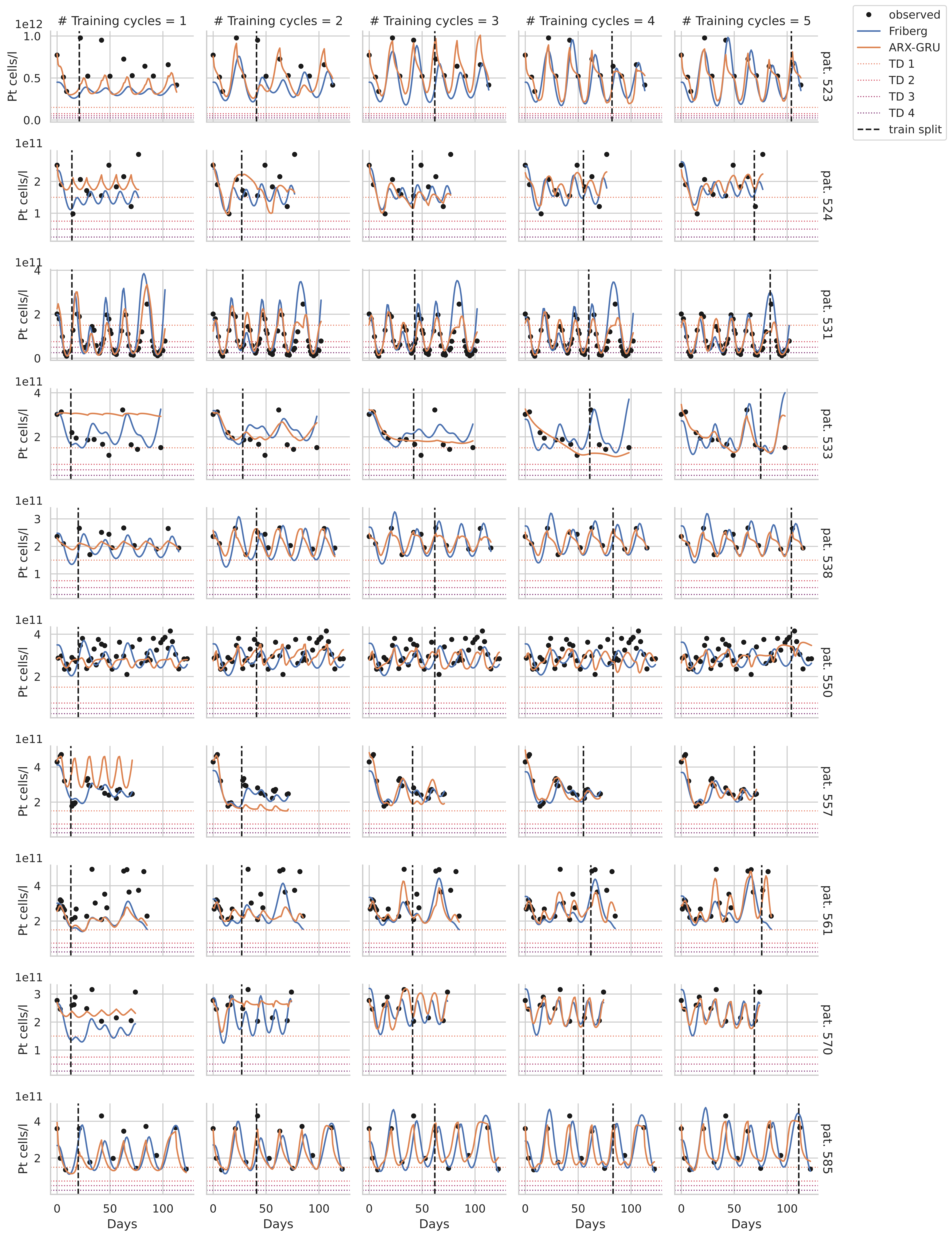

### Training cycles = 1
### Training cycles = 2
### Training cycles = 3
### Training cycles = 4
### Training cycles = 5
observed
Friberg
ARX-GRU
TD 1
TD 2
TD 3
TD 4
train split
Pt cells/l
Days
pat. 523
pat. 524
pat. 531
pat. 533
pat. 538
pat. 550
pat. 557
pat. 561
pat. 570
pat. 585
1e12
1e11

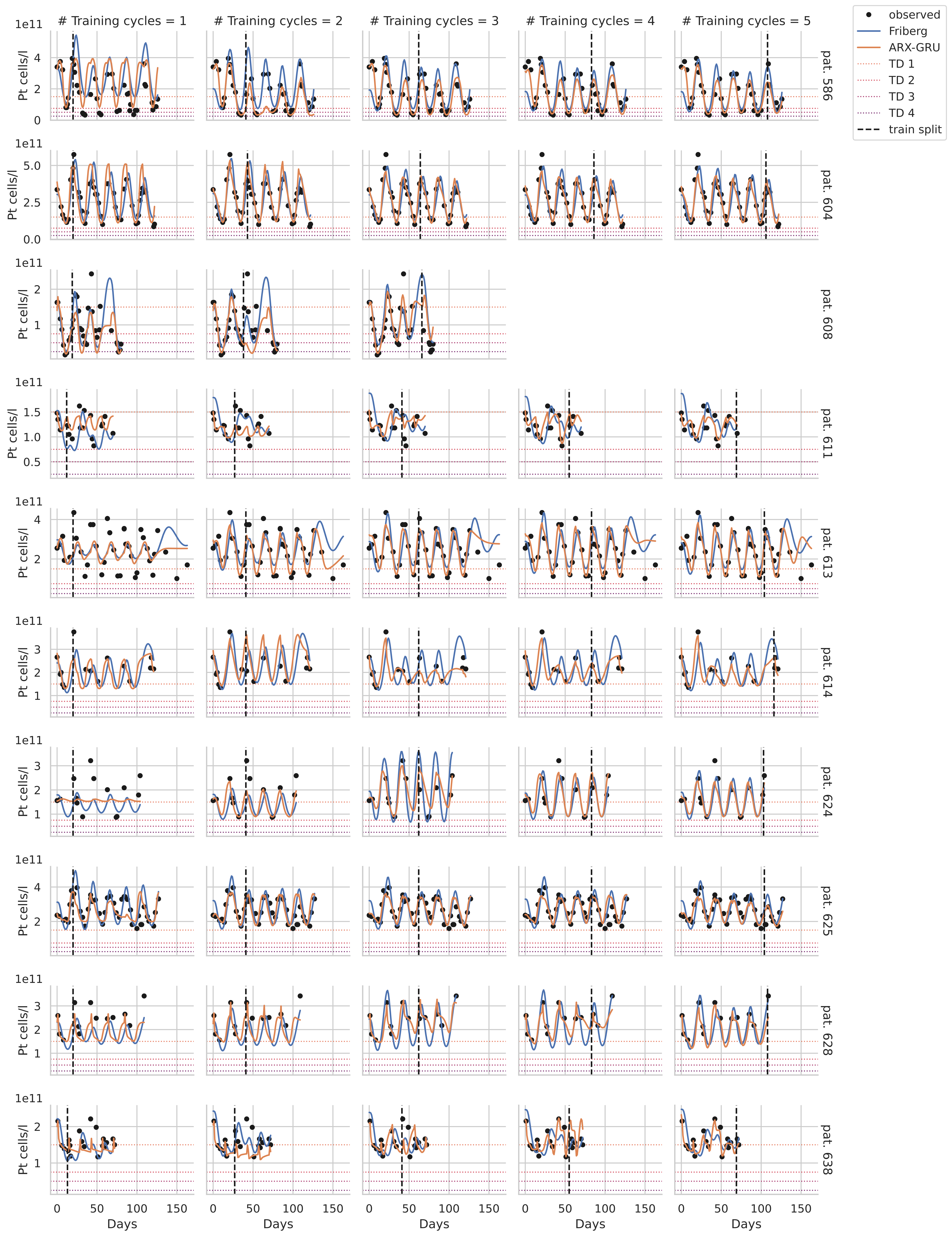

### Training cycles = 1
### Training cycles = 2
### Training cycles = 3
### Training cycles = 4
### Training cycles = 5
observed
Friberg
ARX-GRU
TD 1
TD 2
TD 3
TD 4
train split
Pt cells/l
1e11
pat. 586
pat. 604
pat. 608
pat. 611
pat. 613
pat. 614
pat. 624
pat. 625
pat. 628
pat. 638
Days

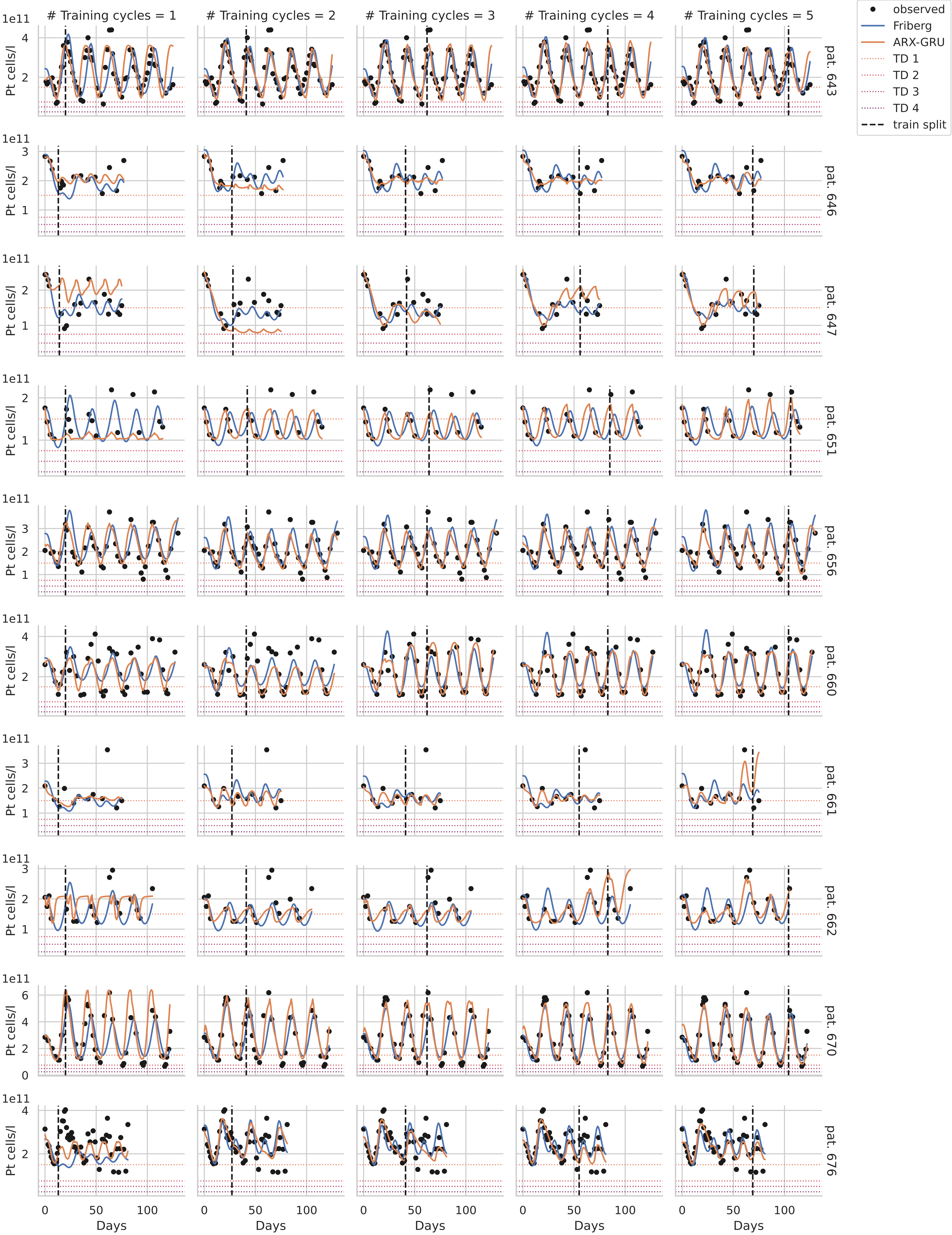

### Training cycles = 1
### Training cycles = 2
### Training cycles = 3
### Training cycles = 4
### Training cycles = 5
observed
Friberg
ARX-GRU
TD 1
TD 2
TD 3
TD 4
train split
Pt cells/l
1e11
pat. 643
pat. 646
pat. 647
pat. 651
pat. 656
pat. 660
pat. 661
pat. 662
pat. 670
pat. 676
Days

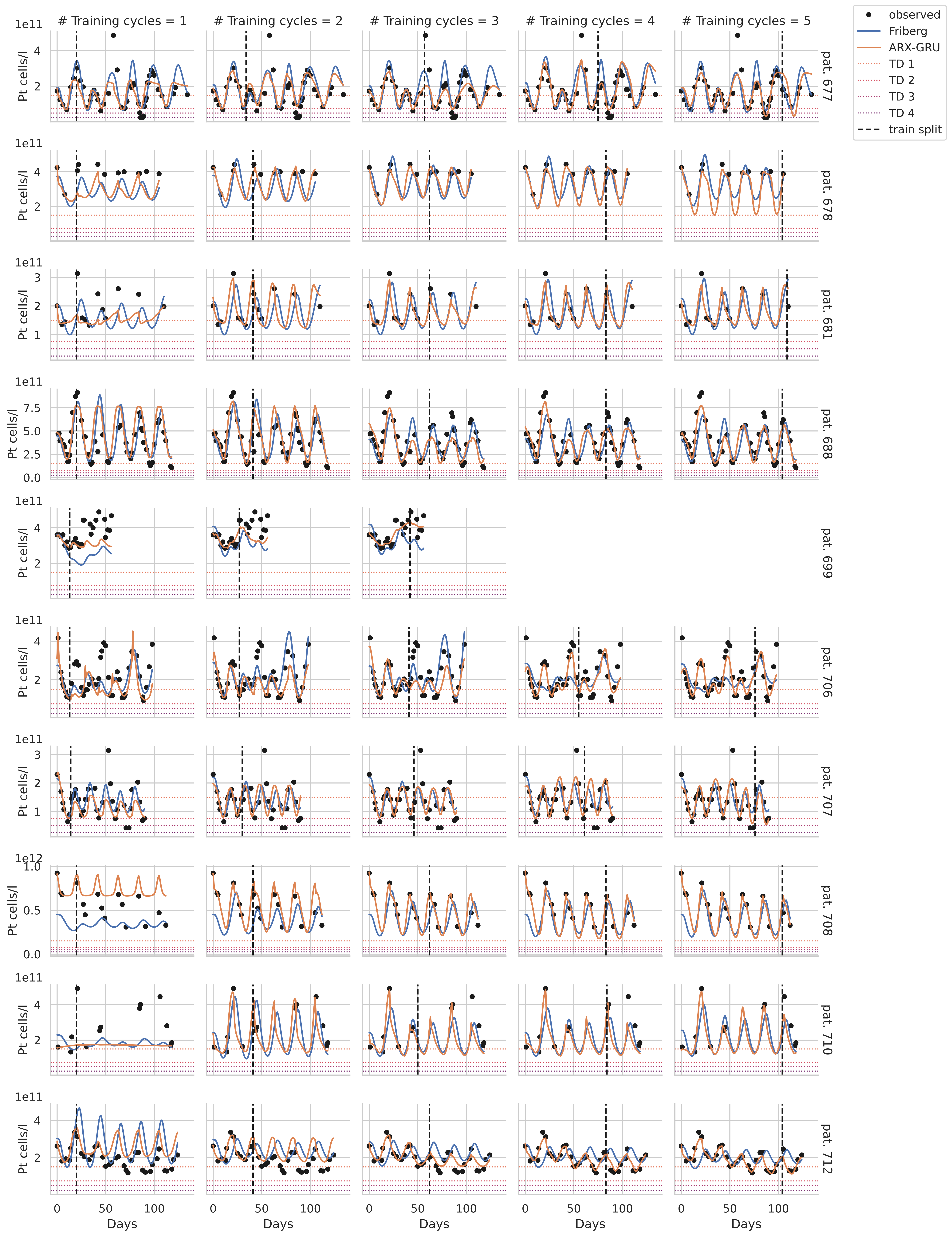

### Training cycles = 1
### Training cycles = 2
### Training cycles = 3
### Training cycles = 4
### Training cycles = 5
observed
Friberg
ARX-GRU
TD 1
TD 2
TD 3
TD 4
train split
Pt cells/l
Days
pat. 677
pat. 678
pat. 681
pat. 688
pat. 699
pat. 706
pat. 707
pat. 708
pat. 710
pat. 712

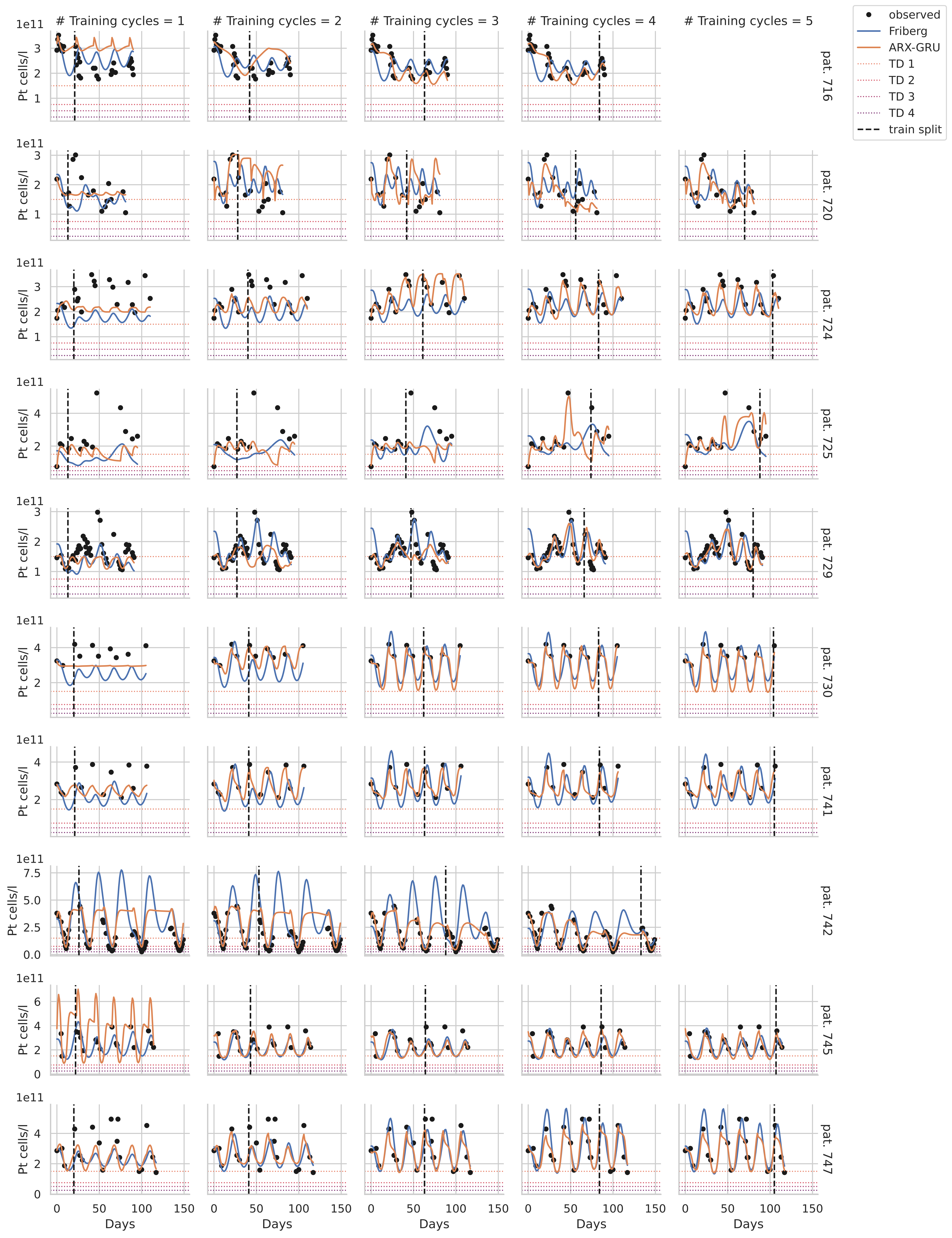

### Training cycles = 1
### Training cycles = 2
### Training cycles = 3
### Training cycles = 4
### Training cycles = 5
observed
Friberg
ARX-GRU
TD 1
TD 2
TD 3
TD 4
train split
Pt cells/l
1e11
Days
pat. 716
pat. 720
pat. 724
pat. 725
pat. 729
pat. 730
pat. 741
pat. 742
pat. 745
pat. 747

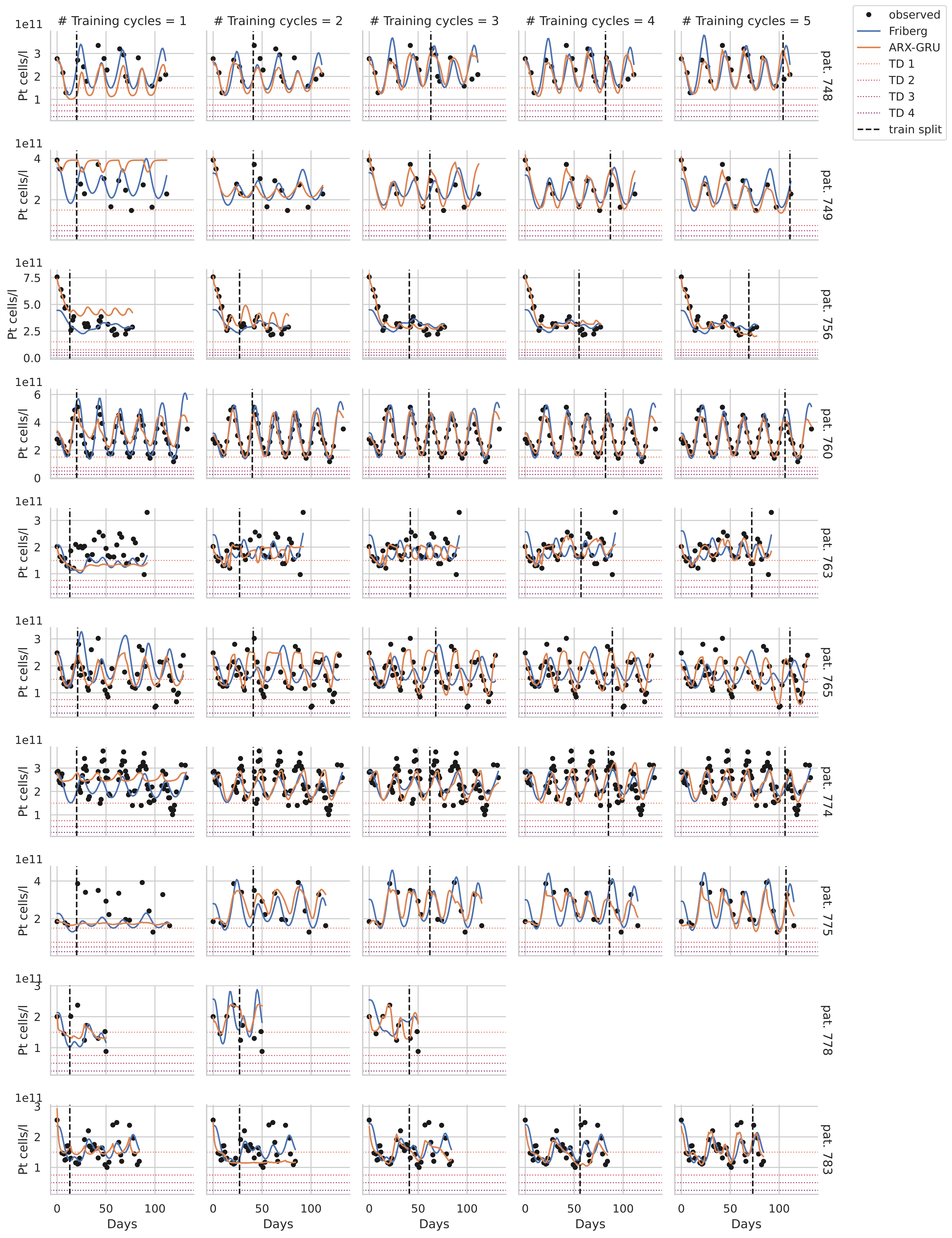

### Training cycles = 1
### Training cycles = 2
### Training cycles = 3
### Training cycles = 4
### Training cycles = 5
observed
Friberg
ARX-GRU
TD 1
TD 2
TD 3
TD 4
train split
Pt cells/l
1e11
Days
pat. 748
pat. 749
pat. 756
pat. 760
pat. 763
pat. 765
pat. 774
pat. 775
pat. 778
pat. 783

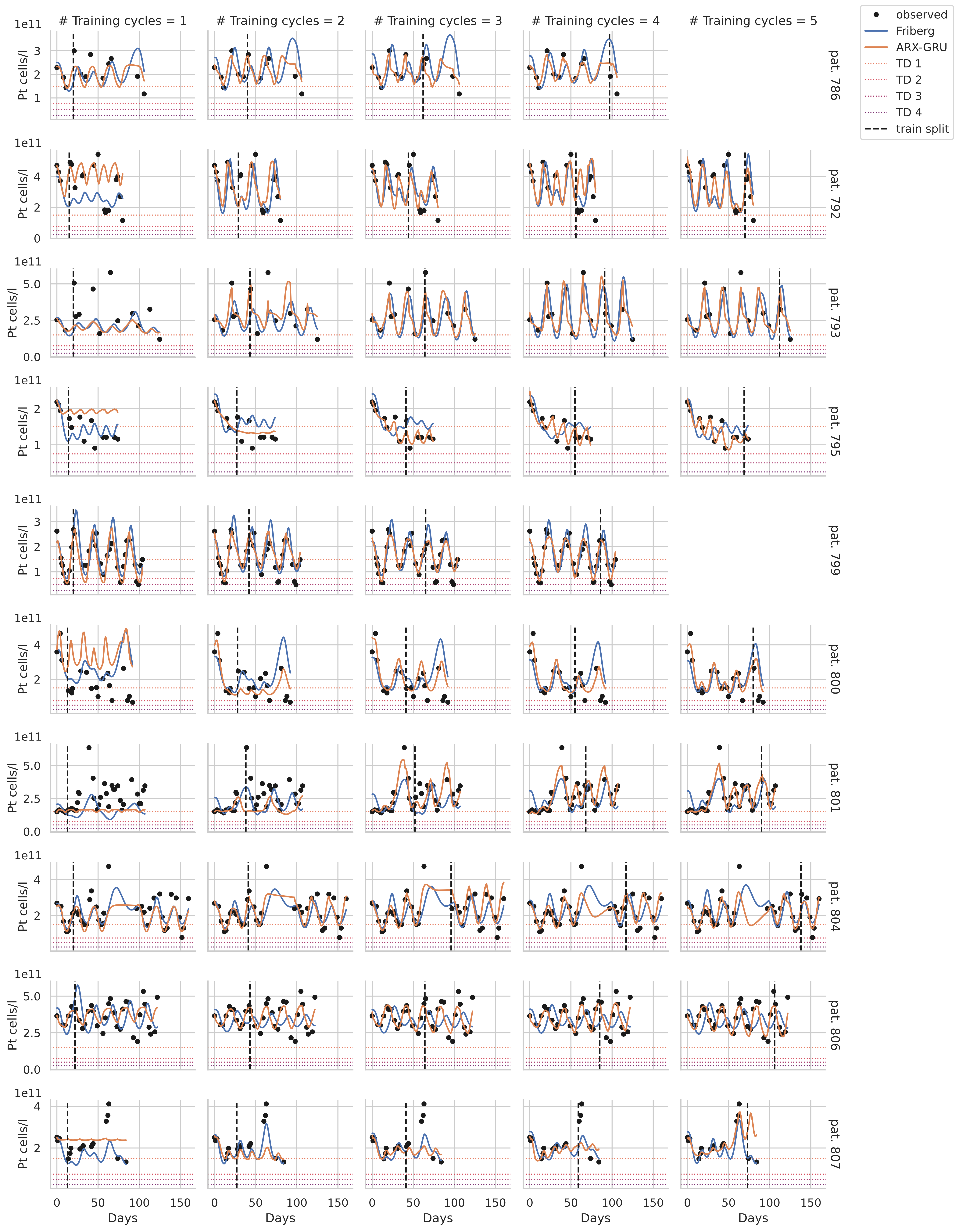

### Training cycles = 1
### Training cycles = 2
### Training cycles = 3
### Training cycles = 4
### Training cycles = 5
observed
Friberg
ARX-GRU
TD 1
TD 2
TD 3
TD 4
train split
Pt cells/l
1e11
pat. 786
pat. 792
pat. 793
pat. 795
pat. 799
pat. 800
pat. 801
pat. 804
pat. 806
pat. 807
Days

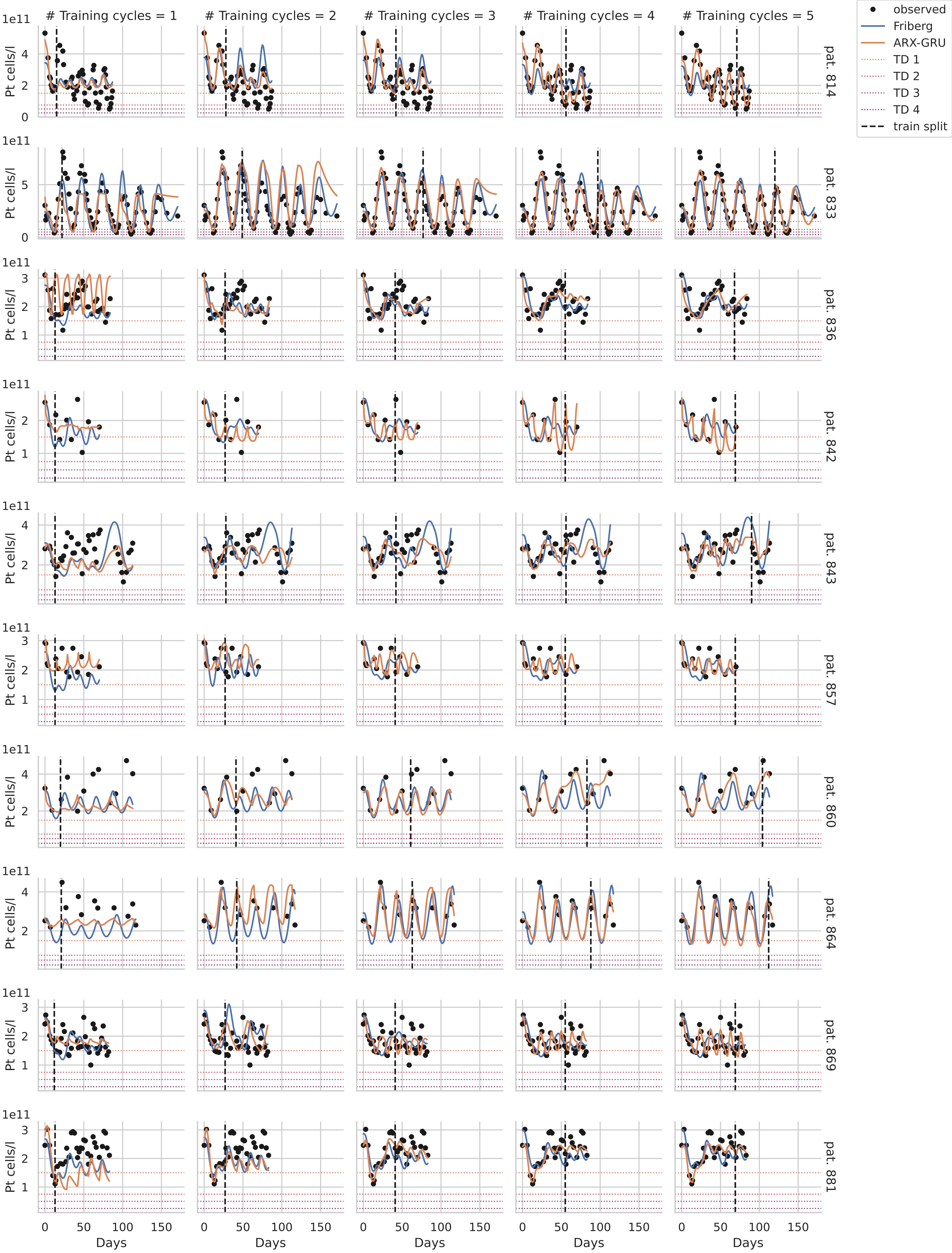

### Training cycles = 1
### Training cycles = 2
### Training cycles = 3
### Training cycles = 4
### Training cycles = 5
observed
Friberg
ARX-GRU
TD 1
TD 2
TD 3
TD 4
train split
pat. 814
pat. 833
pat. 836
pat. 842
pat. 843
pat. 857
pat. 860
pat. 864
pat. 869
pat. 881
Pt cells/l
1e11
Days

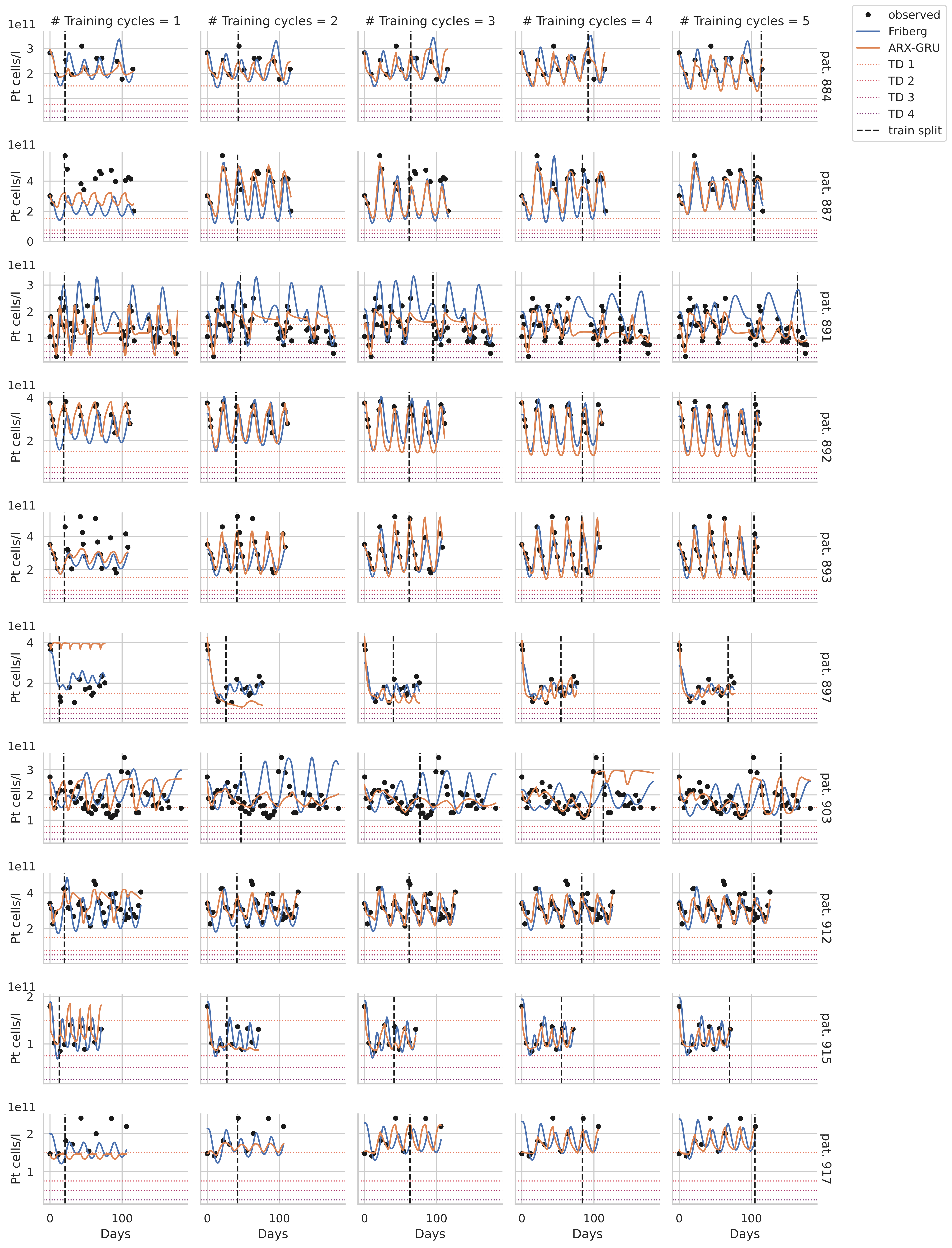

### Training cycles = 1
### Training cycles = 2
### Training cycles = 3
### Training cycles = 4
### Training cycles = 5
observed
Friberg
ARX-GRU
TD 1
TD 2
TD 3
TD 4
train split
Pt cells/l
1e11
Days
pat. 884
pat. 887
pat. 891
pat. 892
pat. 893
pat. 897
pat. 903
pat. 912
pat. 915
pat. 917

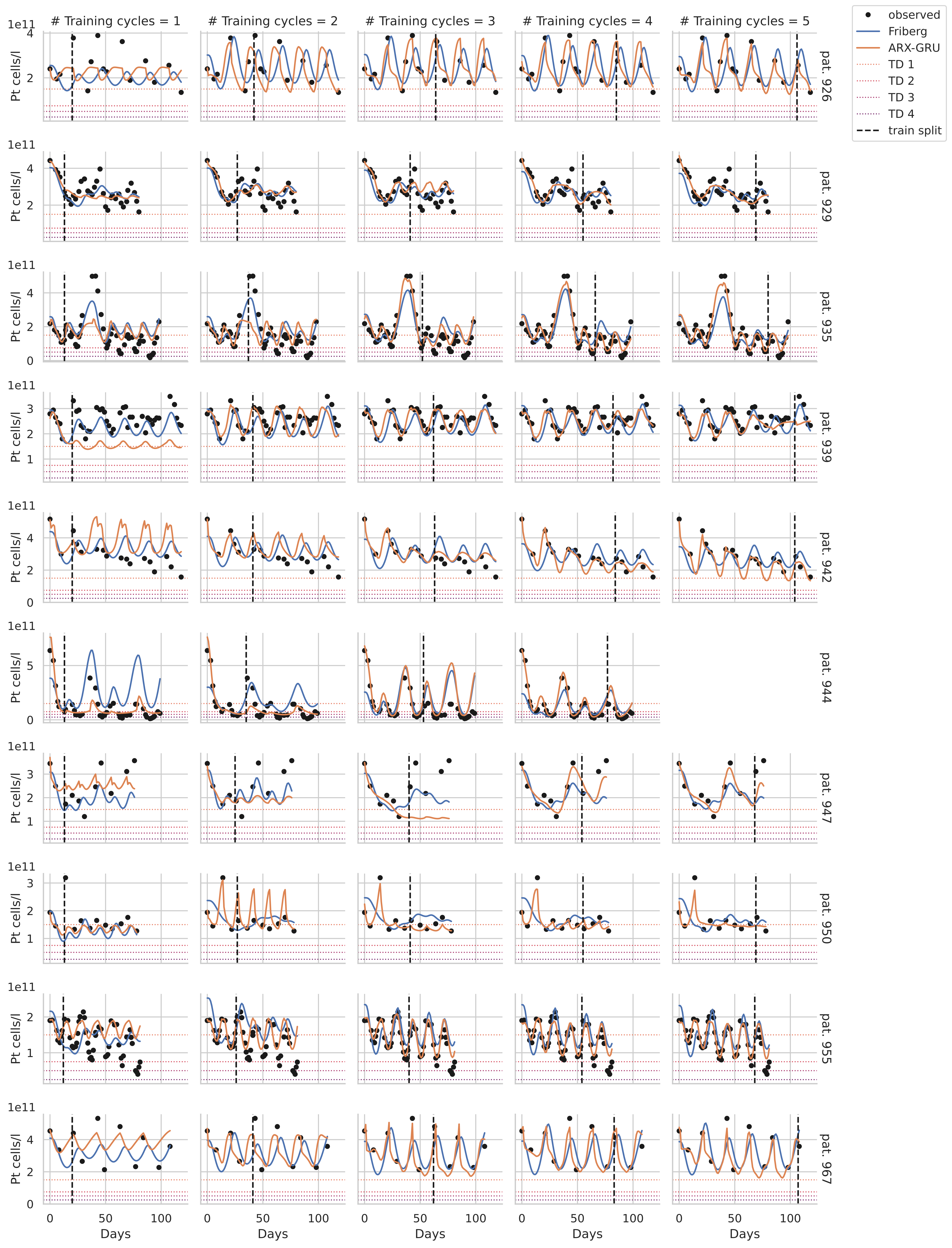

### Training cycles = 1
### Training cycles = 2
### Training cycles = 3
### Training cycles = 4
### Training cycles = 5
observed
Friberg
ARX-GRU
TD 1
TD 2
TD 3
TD 4
train split
pat. 926
pat. 929
pat. 935
pat. 939
pat. 942
pat. 944
pat. 947
pat. 950
pat. 955
pat. 967
Pt cells/l
1e11
Days

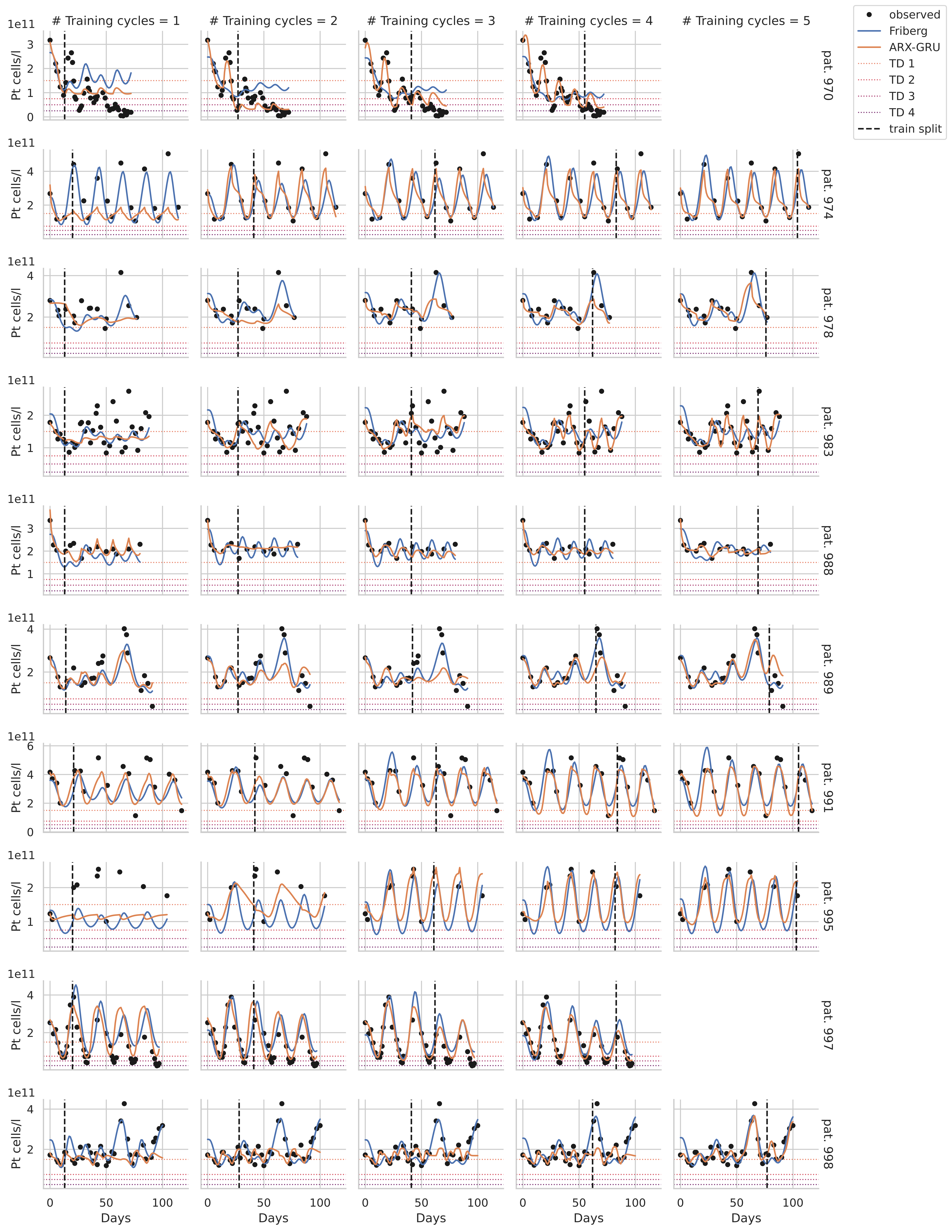

### Training cycles = 1
### Training cycles = 2
### Training cycles = 3
### Training cycles = 4
### Training cycles = 5
observed
Friberg
ARX-GRU
TD 1
TD 2
TD 3
TD 4
train split
1e11
Pt cells/l
pat. 970
pat. 974
pat. 978
pat. 983
pat. 988
pat. 989
pat. 991
pat. 995
pat. 997
pat. 998
Days

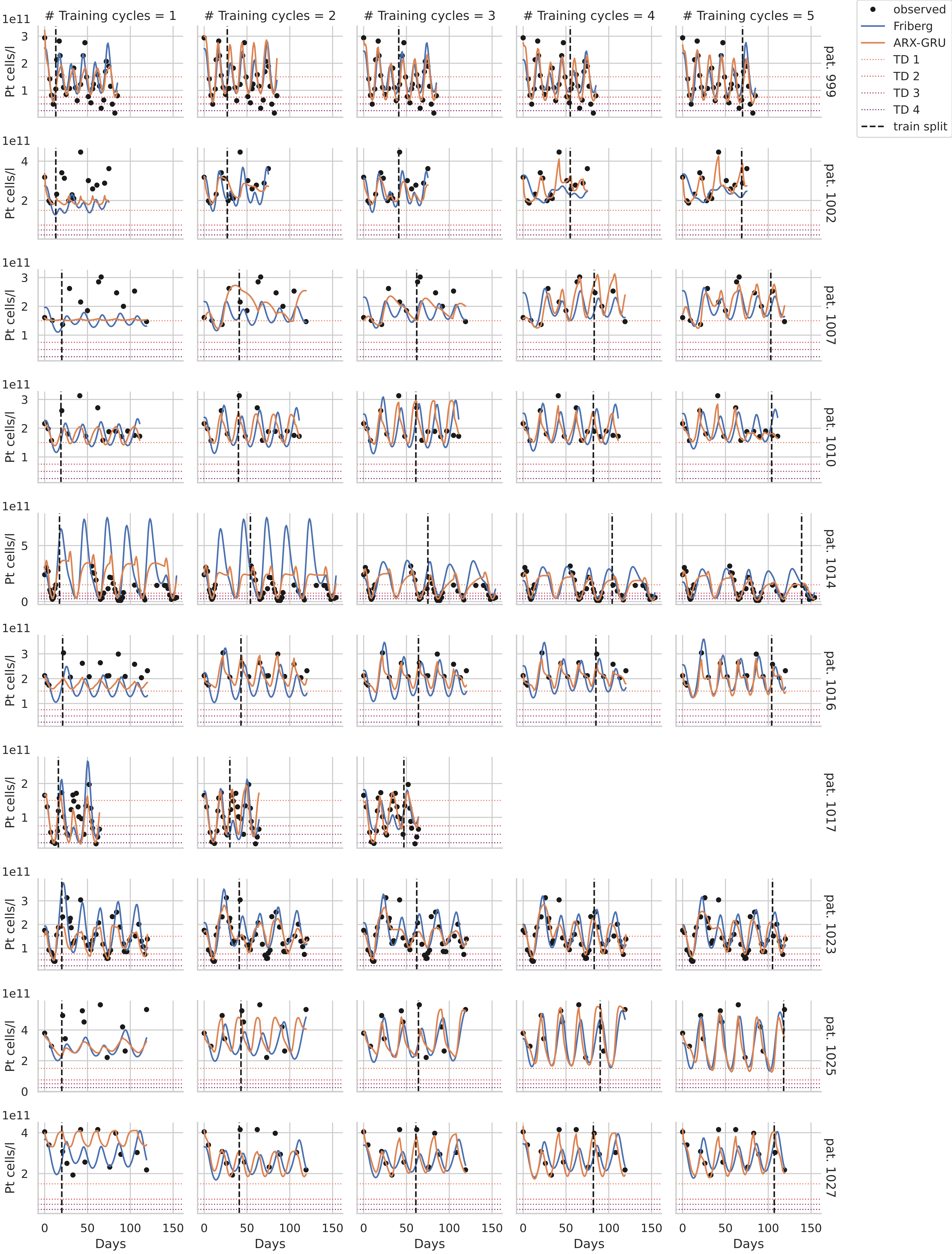

### Training cycles = 1
### Training cycles = 2
### Training cycles = 3
### Training cycles = 4
### Training cycles = 5
observed
Friberg
ARX-GRU
TD 1
TD 2
TD 3
TD 4
train split
Pt cells/l
1e11
pat. 999
pat. 1002
pat. 1007
pat. 1010
pat. 1014
pat. 1016
pat. 1017
pat. 1023
pat. 1025
pat. 1027
Days

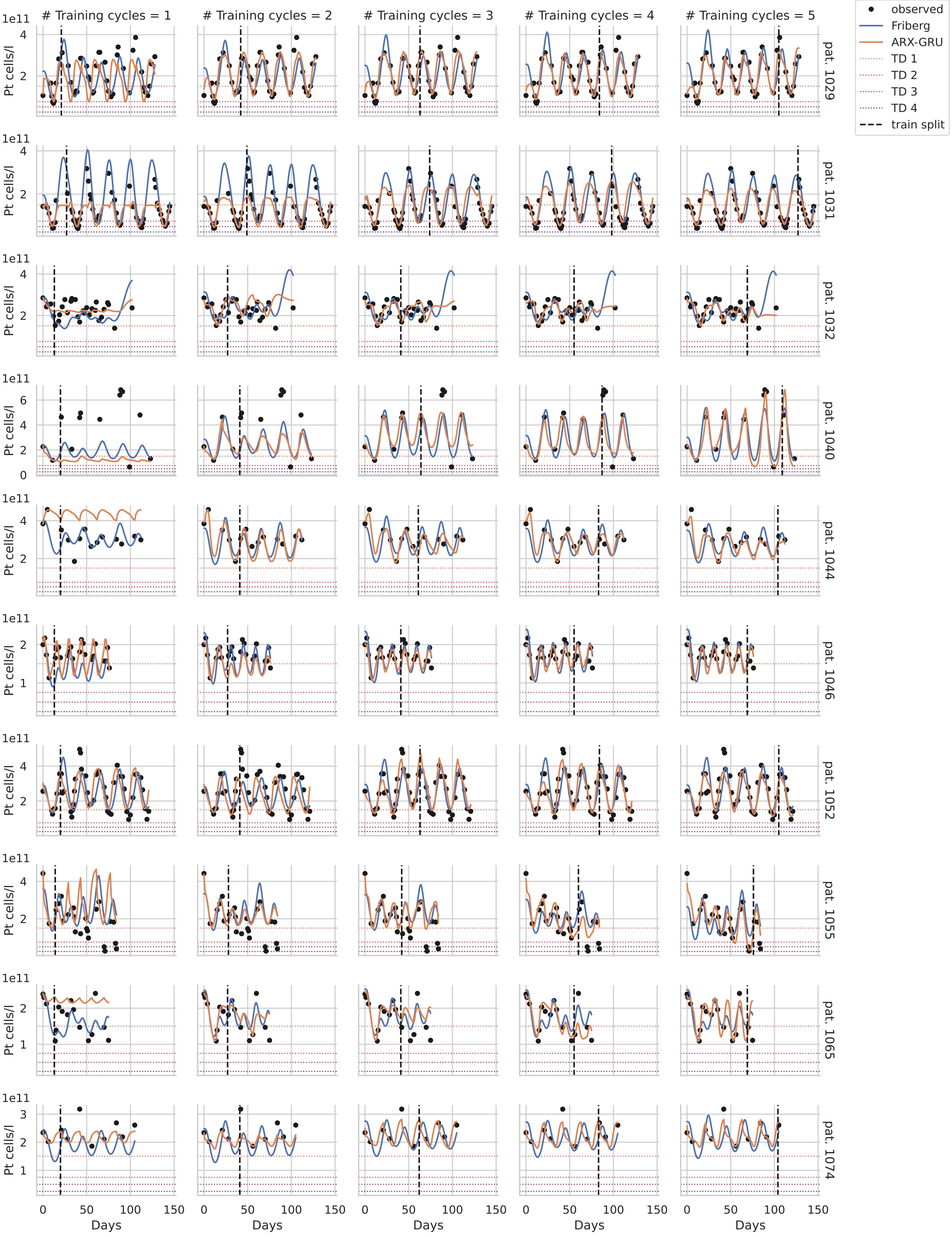

### Training cycles = 1
### Training cycles = 2
### Training cycles = 3
### Training cycles = 4
### Training cycles = 5
observed
Friberg
ARX-GRU
TD 1
TD 2
TD 3
TD 4
train split
Pt cells/l
1e11
pat. 1029
pat. 1031
pat. 1032
pat. 1040
pat. 1044
pat. 1046
pat. 1052
pat. 1055
pat. 1065
pat. 1074
Days

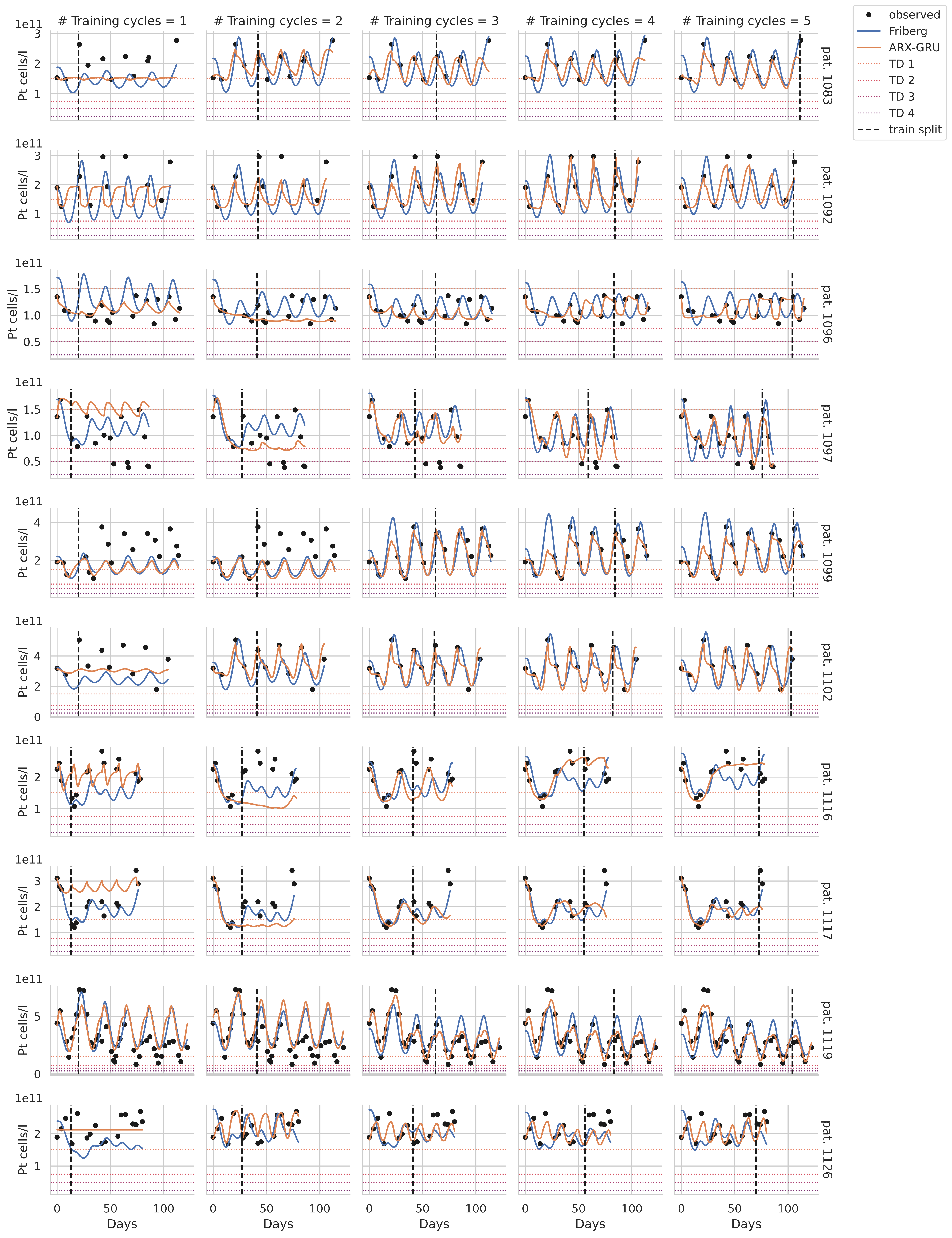

### Training cycles = 1
### Training cycles = 2
### Training cycles = 3
### Training cycles = 4
### Training cycles = 5
observed
Friberg
ARX-GRU
TD 1
TD 2
TD 3
TD 4
train split
Pt cells/l
1e11
Days
pat. 1083
pat. 1092
pat. 1096
pat. 1097
pat. 1099
pat. 1102
pat. 1116
pat. 1117
pat. 1119
pat. 1126

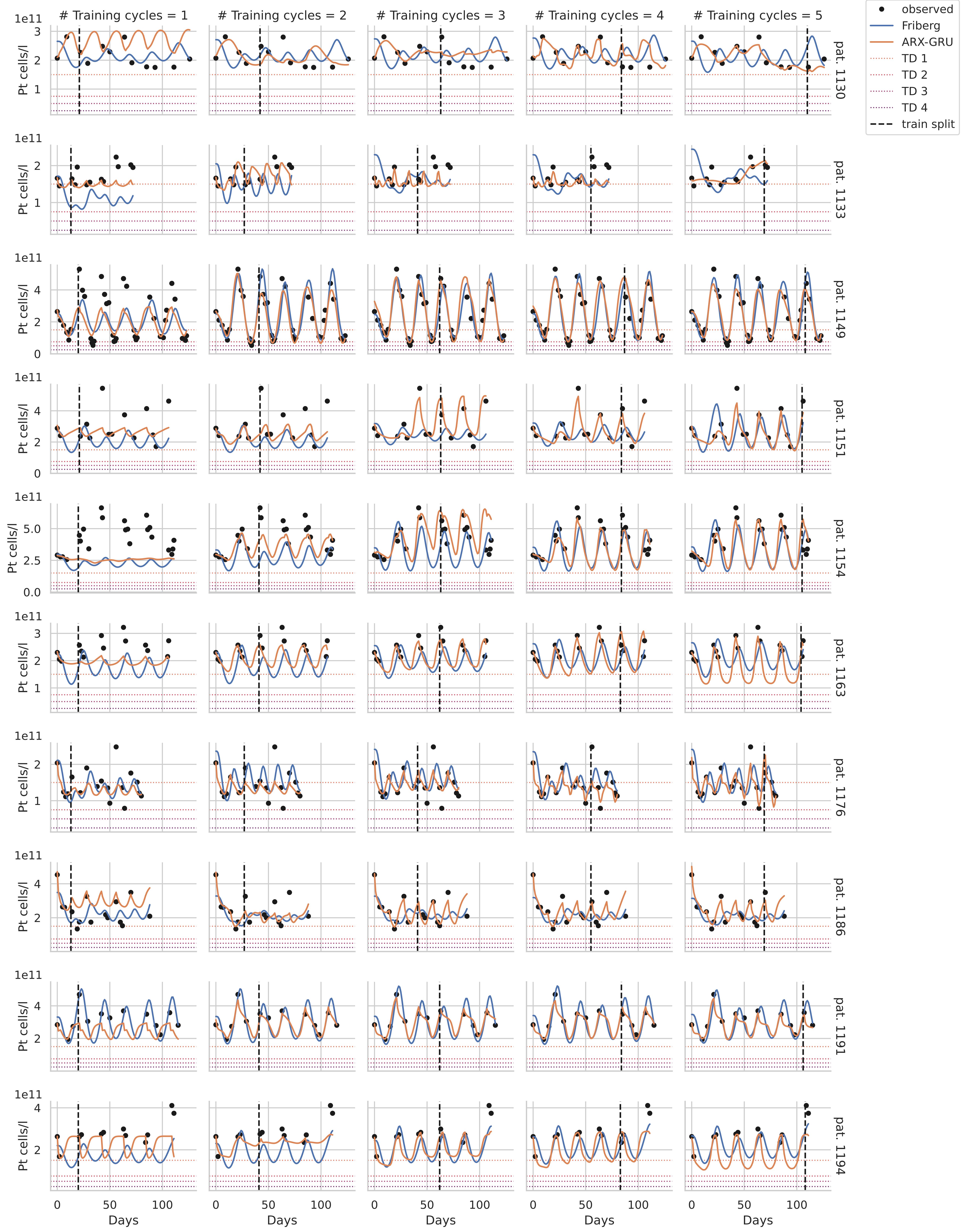

### Training cycles = 1
### Training cycles = 2
### Training cycles = 3
### Training cycles = 4
### Training cycles = 5
observed
Friberg
ARX-GRU
TD 1
TD 2
TD 3
TD 4
train split
Pt cells/l
1e11
pat. 1130
pat. 1133
pat. 1149
pat. 1151
pat. 1154
pat. 1163
pat. 1176
pat. 1186
pat. 1191
pat. 1194
Days
0
50
100

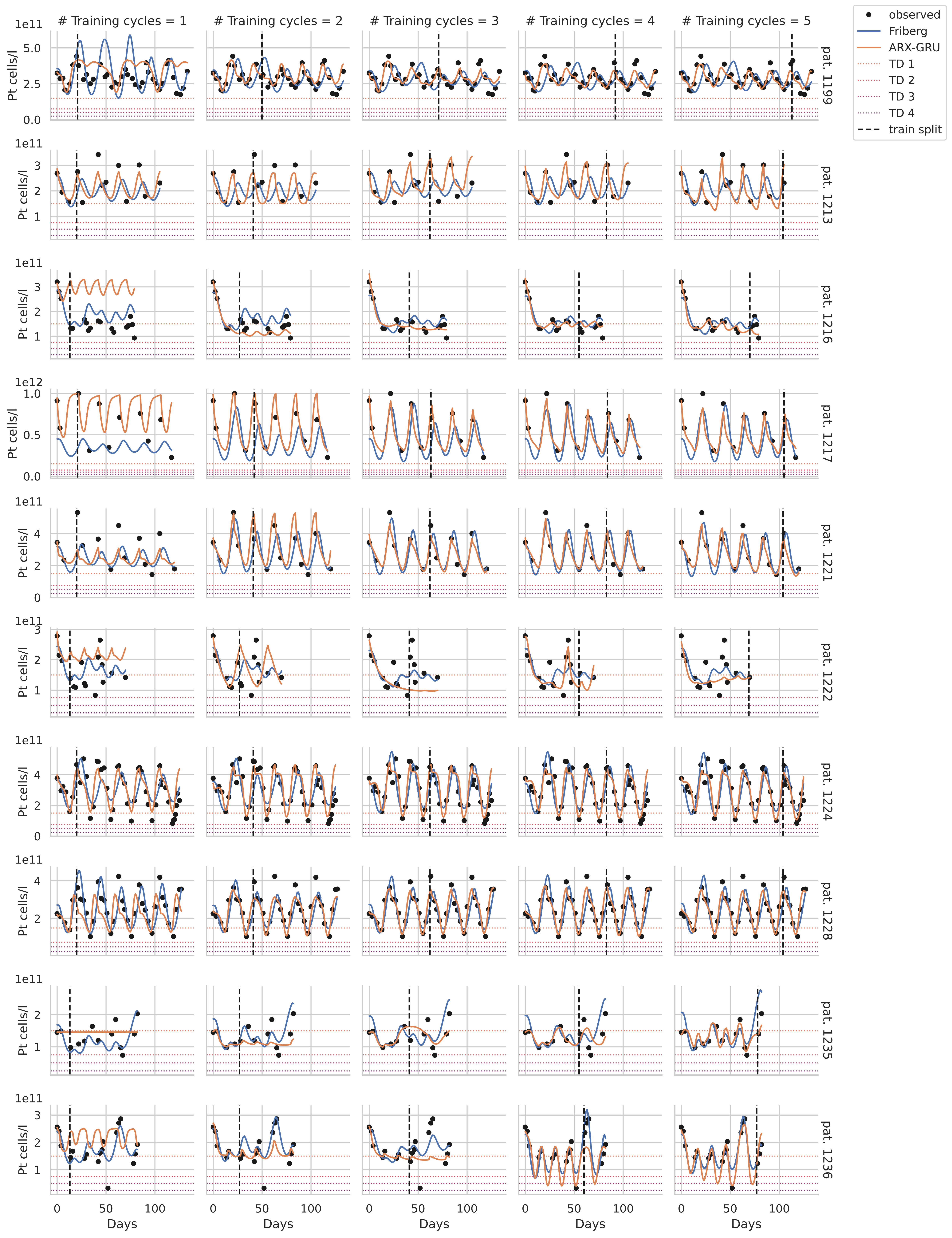

### Training cycles = 1
### Training cycles = 2
### Training cycles = 3
### Training cycles = 4
### Training cycles = 5
observed
Friberg
ARX-GRU
TD 1
TD 2
TD 3
TD 4
train split
Pt cells/l
Days
pat. 1199
pat. 1213
pat. 1216
pat. 1217
pat. 1221
pat. 1222
pat. 1224
pat. 1228
pat. 1235
pat. 1236

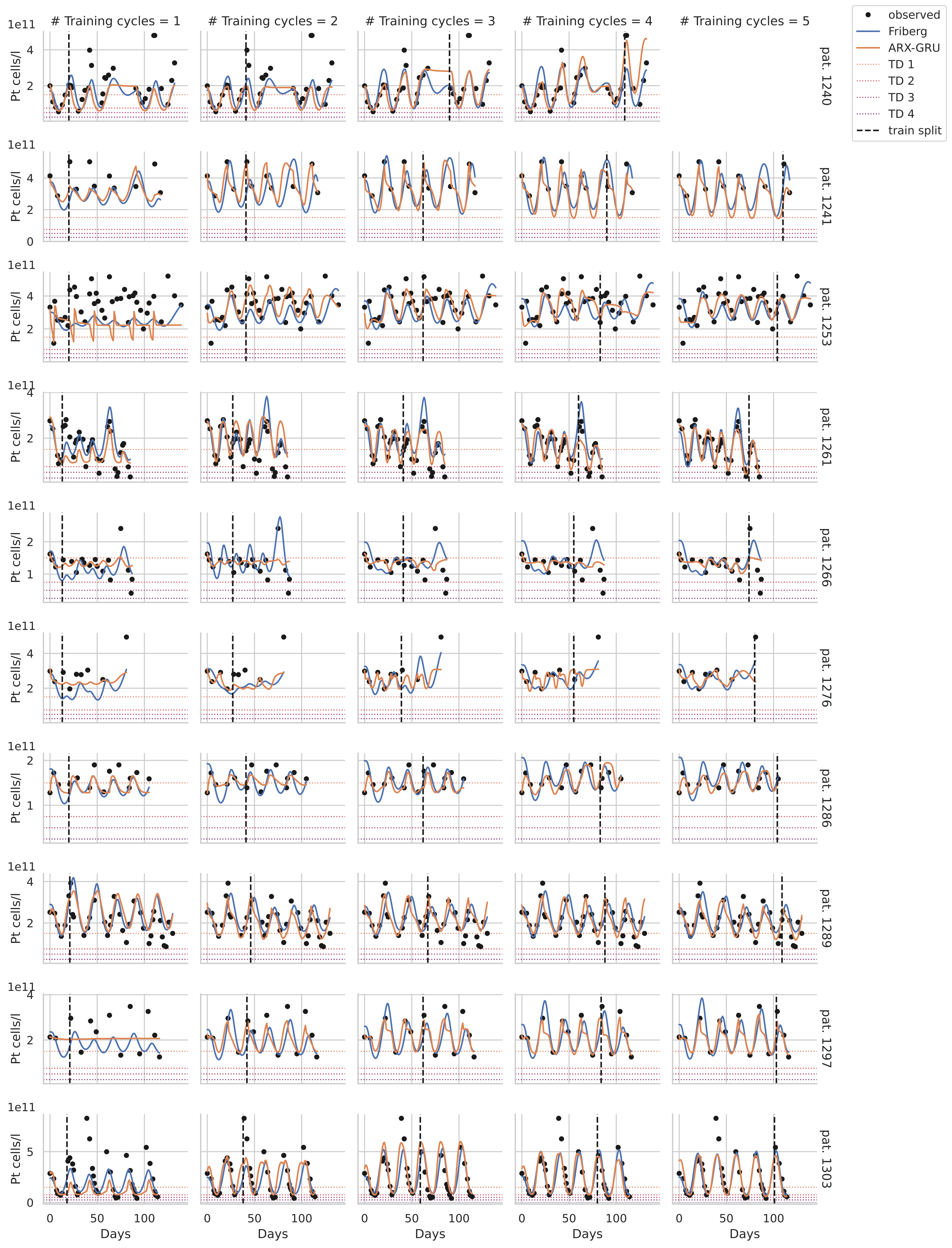

### Training cycles = 1
### Training cycles = 2
### Training cycles = 3
### Training cycles = 4
### Training cycles = 5
observed
Friberg
ARX-GRU
TD 1
TD 2
TD 3
TD 4
train split
Pt cells/l
1e11
Days
pat. 1240
pat. 1241
pat. 1253
pat. 1261
pat. 1266
pat. 1276
pat. 1286
pat. 1289
pat. 1297
pat. 1303

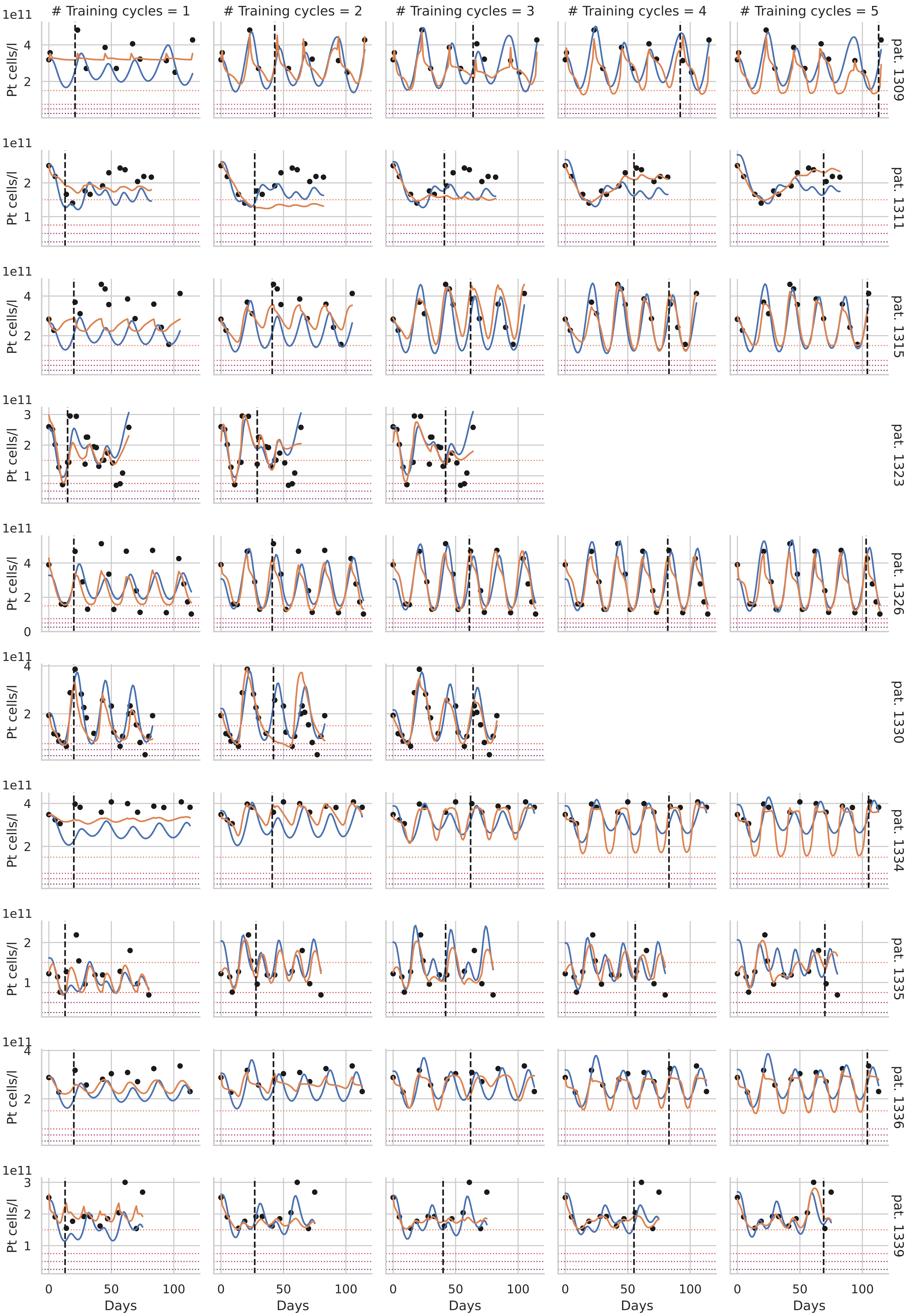

### Training cycles = 1
### Training cycles = 2
### Training cycles = 3
### Training cycles = 4
### Training cycles = 5
Pt cells/l
1e11
Days
pat. 1309
pat. 1311
pat. 1315
pat. 1323
pat. 1326
pat. 1330
pat. 1334
pat. 1335
pat. 1336
pat. 1339

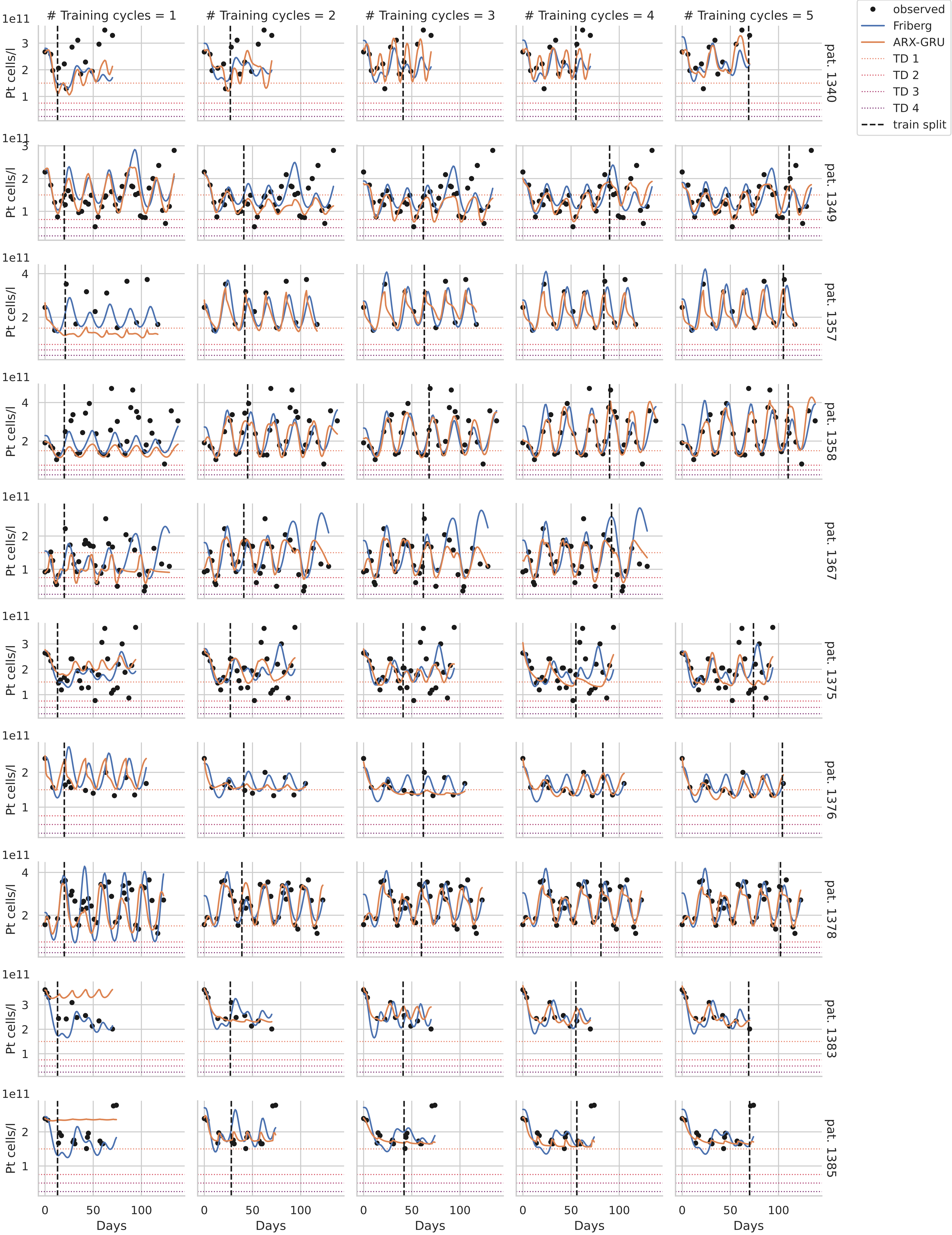

### Training cycles = 1
### Training cycles = 2
### Training cycles = 3
### Training cycles = 4
### Training cycles = 5
observed
Friberg
ARX-GRU
TD 1
TD 2
TD 3
TD 4
train split
1e11
Pt cells/l
pat. 1340
pat. 1349
pat. 1357
pat. 1358
pat. 1367
pat. 1375
pat. 1376
pat. 1378
pat. 1383
pat. 1385
Days
0
50
100

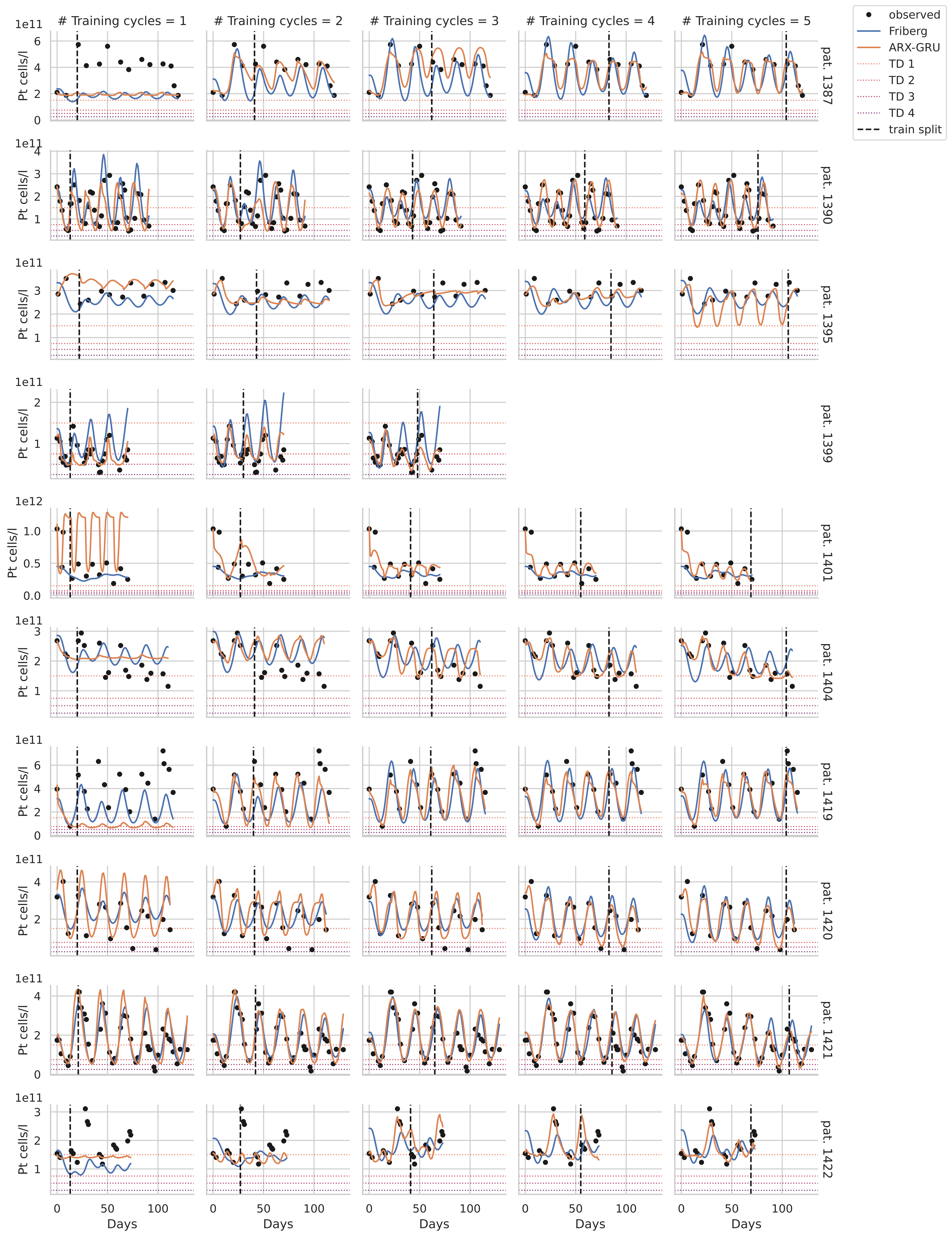

### Training cycles = 1
### Training cycles = 2
### Training cycles = 3
### Training cycles = 4
### Training cycles = 5
observed
Friberg
ARX-GRU
TD 1
TD 2
TD 3
TD 4
train split
Pt cells/l
pat. 1387
pat. 1390
pat. 1395
pat. 1399
pat. 1401
pat. 1404
pat. 1419
pat. 1420
pat. 1421
pat. 1422
Days

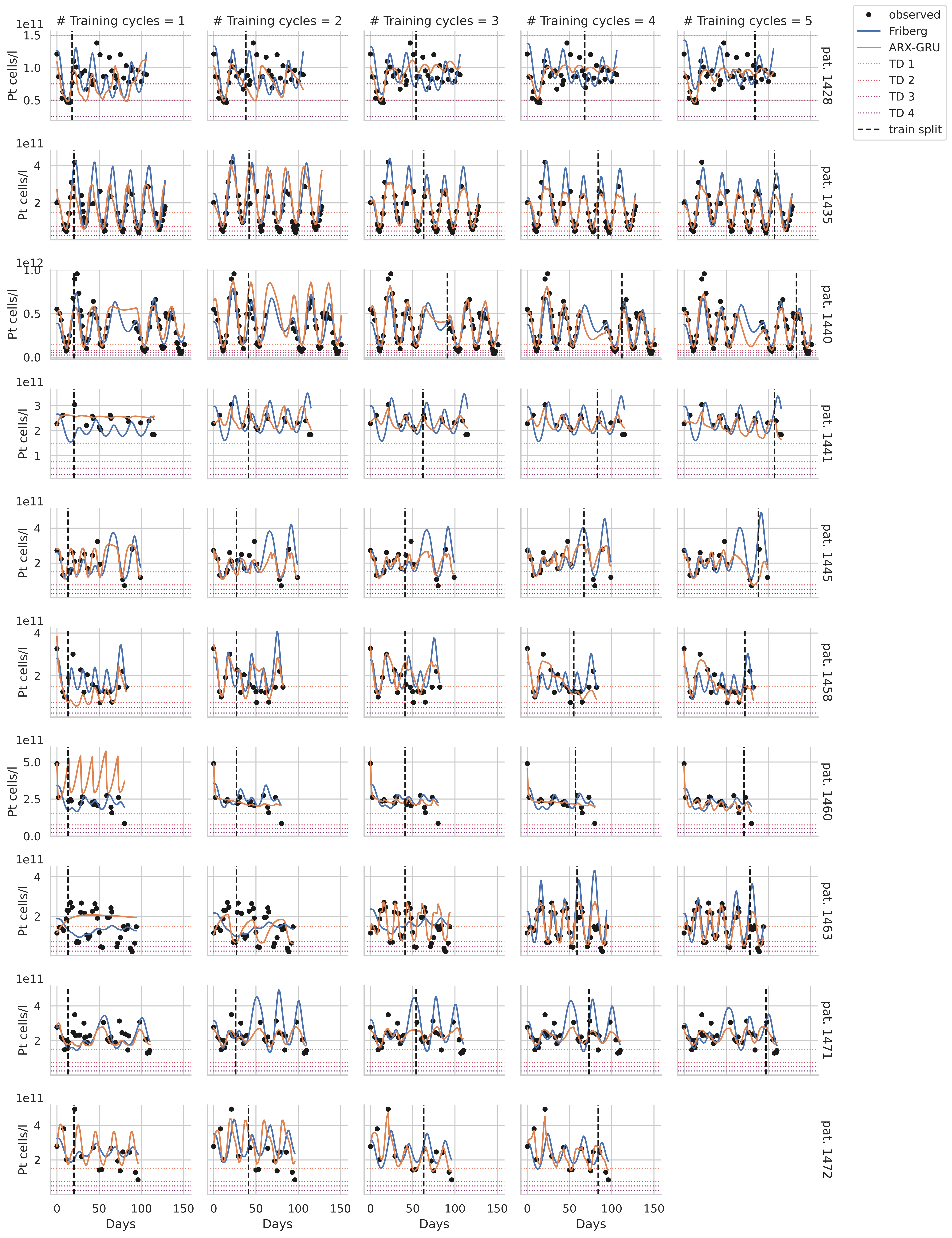

### Training cycles = 1
### Training cycles = 2
### Training cycles = 3
### Training cycles = 4
### Training cycles = 5
observed
Friberg
ARX-GRU
TD 1
TD 2
TD 3
TD 4
train split
pat. 1428
pat. 1435
pat. 1440
pat. 1441
pat. 1445
pat. 1458
pat. 1460
pat. 1463
pat. 1471
pat. 1472
Pt cells/l
Days

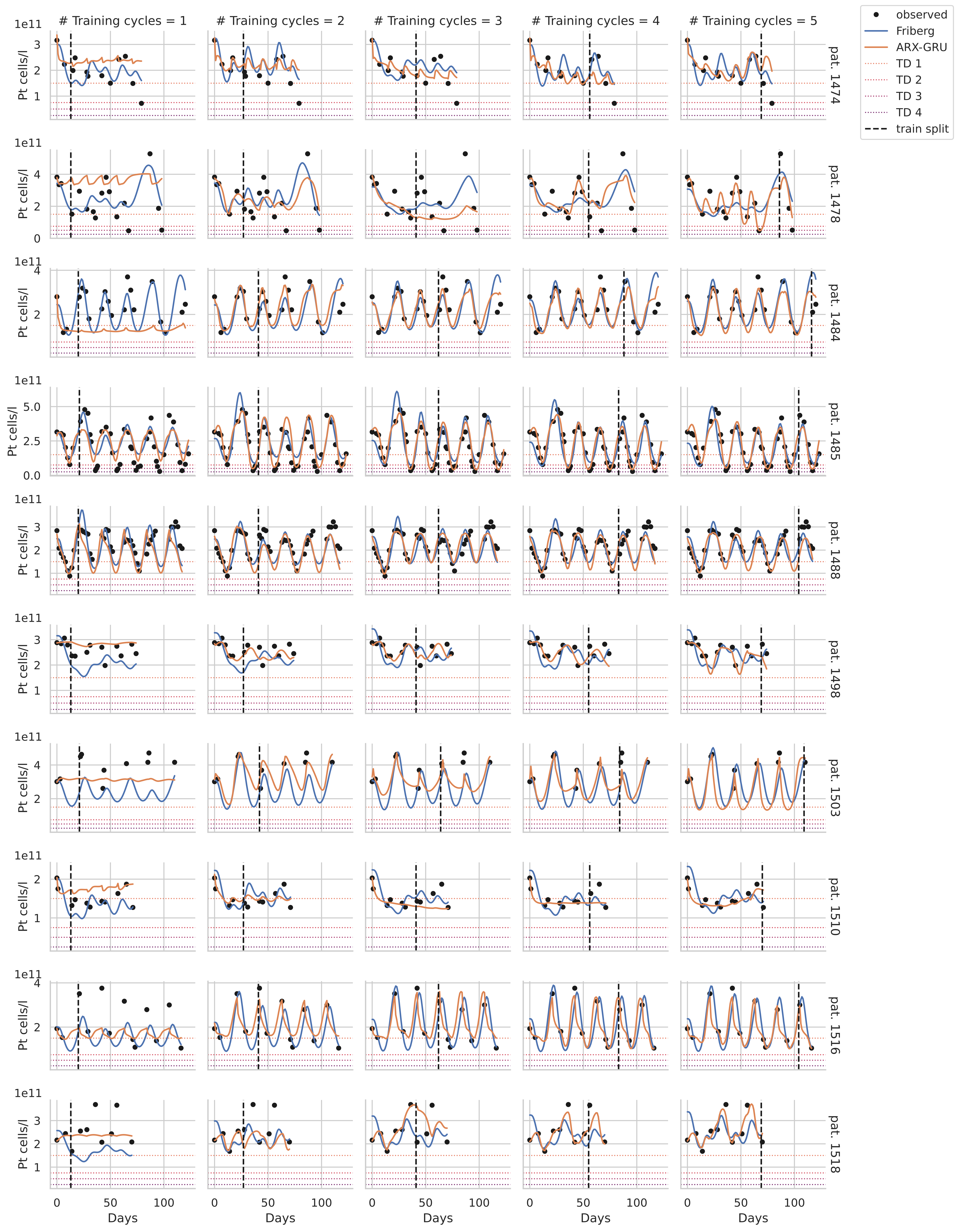

### Training cycles = 1
### Training cycles = 2
### Training cycles = 3
### Training cycles = 4
### Training cycles = 5
observed
Friberg
ARX-GRU
TD 1
TD 2
TD 3
TD 4
train split
Pt cells/l
1e11
Days
pat. 1474
pat. 1478
pat. 1484
pat. 1485
pat. 1488
pat. 1498
pat. 1503
pat. 1510
pat. 1516
pat. 1518

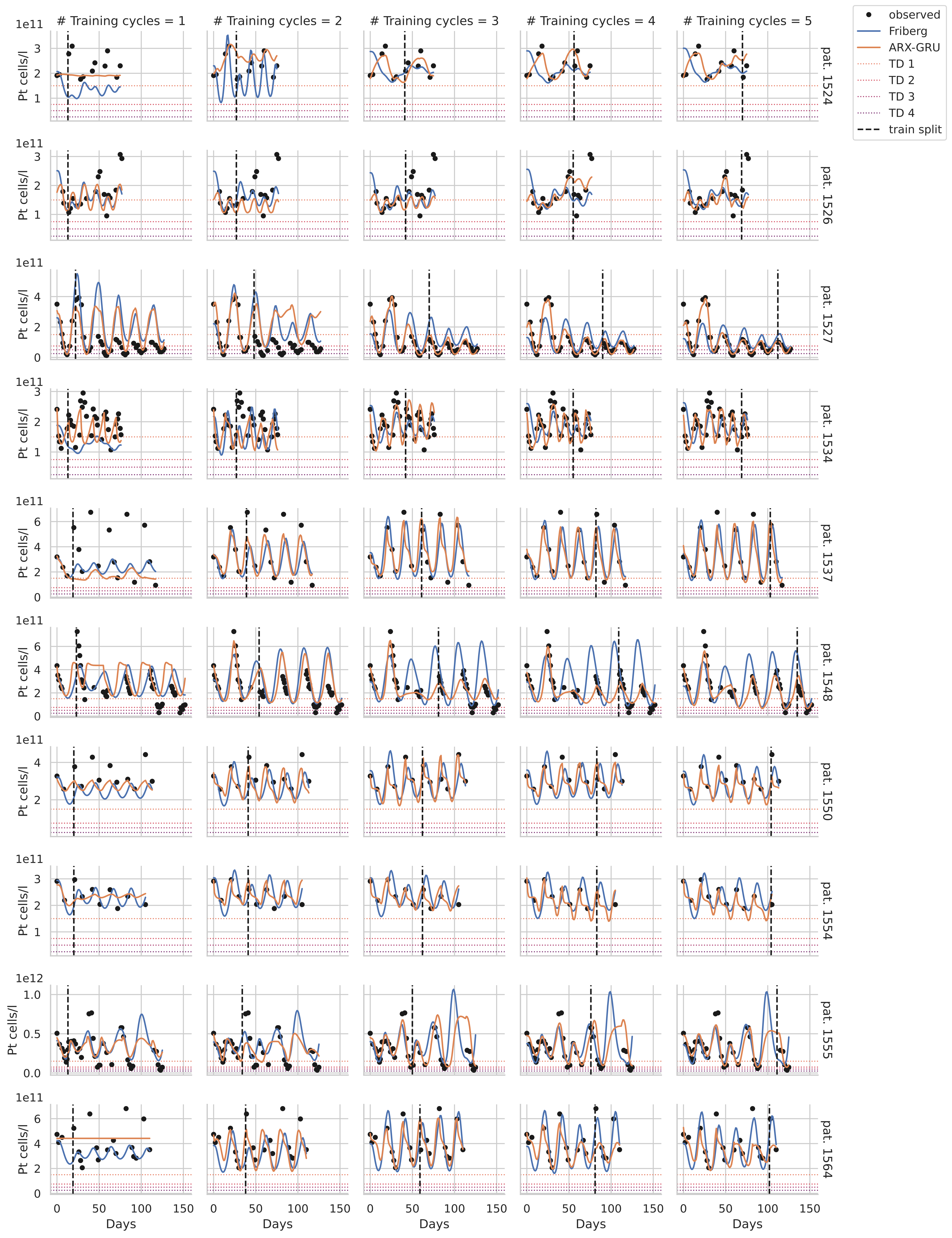

### Training cycles = 1
### Training cycles = 2
### Training cycles = 3
### Training cycles = 4
### Training cycles = 5
observed
Friberg
ARX-GRU
TD 1
TD 2
TD 3
TD 4
train split
Pt cells/l
1e11
1e12
pat. 1524
pat. 1526
pat. 1527
pat. 1534
pat. 1537
pat. 1548
pat. 1550
pat. 1554
pat. 1555
pat. 1564
Days

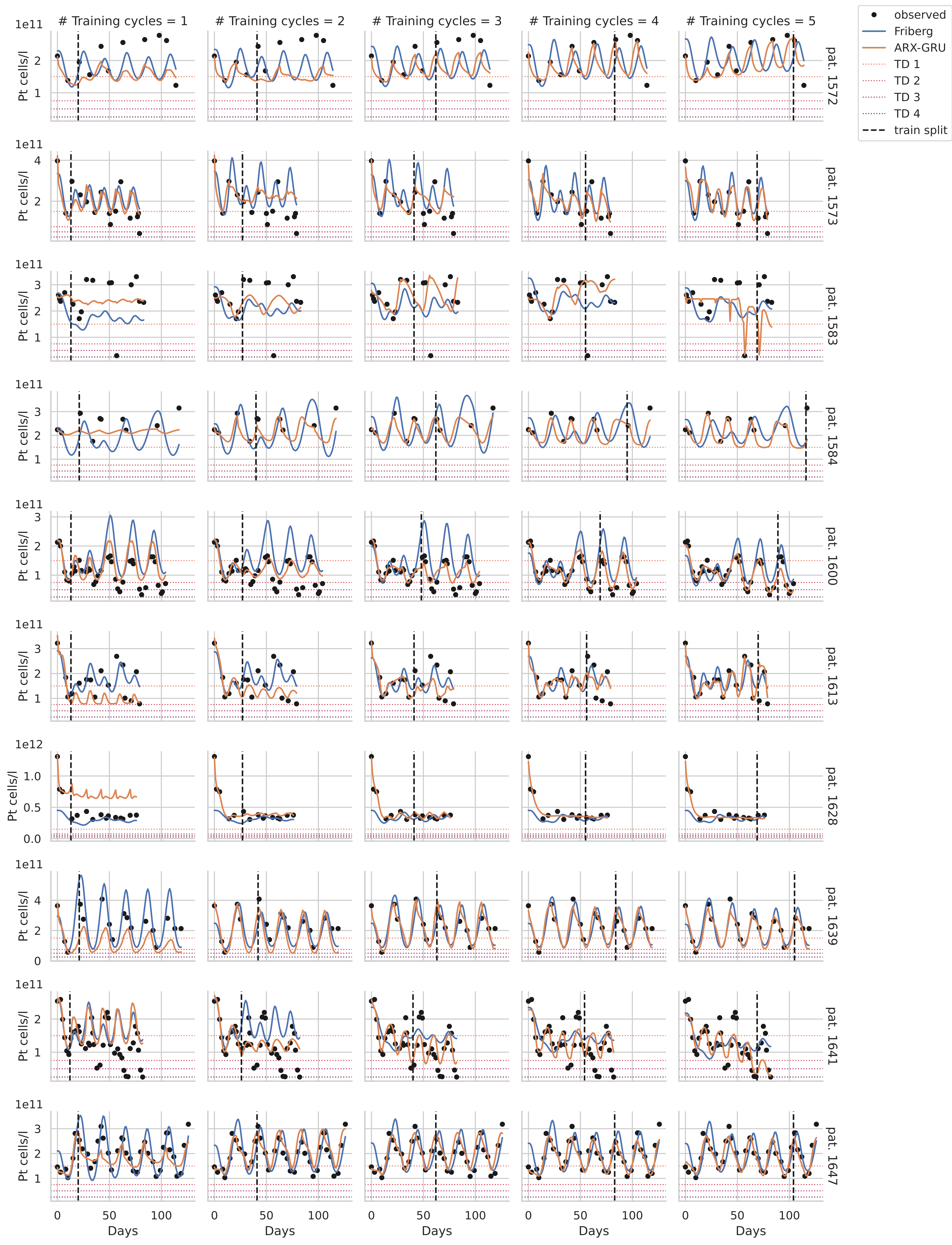

### Training cycles = 1
### Training cycles = 2
### Training cycles = 3
### Training cycles = 4
### Training cycles = 5
observed
Friberg
ARX-GRU
TD 1
TD 2
TD 3
TD 4
train split
Pt cells/l
1e11
1e12
pat. 1572
pat. 1573
pat. 1583
pat. 1584
pat. 1600
pat. 1613
pat. 1628
pat. 1639
pat. 1641
pat. 1647
Days

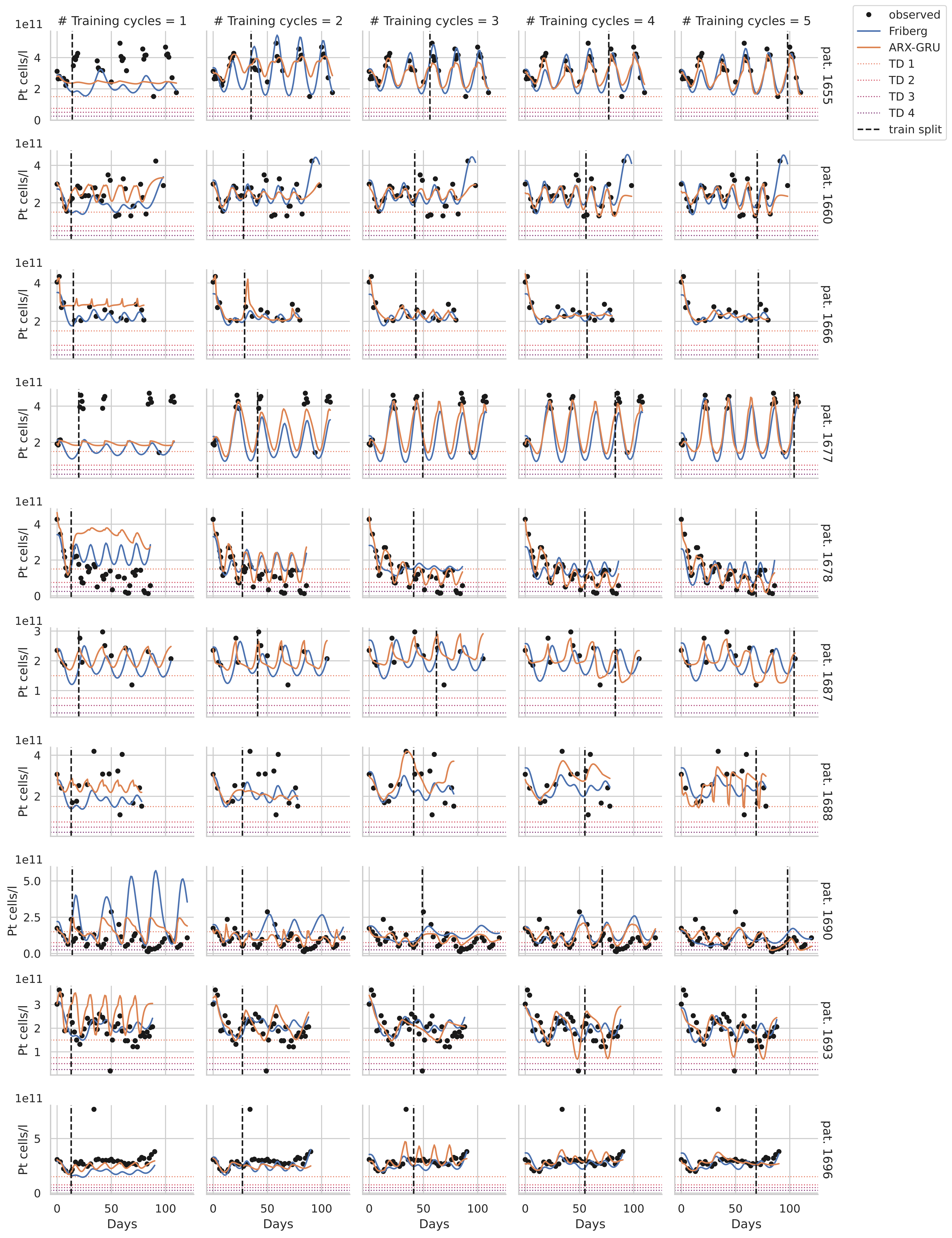

### Training cycles = 1
### Training cycles = 2
### Training cycles = 3
### Training cycles = 4
### Training cycles = 5
observed
Friberg
ARX-GRU
TD 1
TD 2
TD 3
TD 4
train split
Pt cells/l
1e11
pat. 1655
pat. 1660
pat. 1666
pat. 1677
pat. 1678
pat. 1687
pat. 1688
pat. 1690
pat. 1693
pat. 1696
Days
